# Episodic and associative memory from spatial scaffolds in the hippocampus

**DOI:** 10.1101/2023.11.28.568960

**Authors:** Sarthak Chandra, Sugandha Sharma, Rishidev Chaudhuri, Ila Fiete

## Abstract

Hippocampal circuits in the brain enable two distinct cognitive functions: the construction of spatial maps for navigation and the storage of sequential episodic memories. This dual role remains an enduring enigma. While there have been advances in modeling spatial representations in the hippocampus, we lack good models of its role in episodic memory. Here we present a neocortical-entorhinal-hippocampal network model that implements a high-capacity general associative memory, spatial memory, and episodic memory by factorizing content storage from the dynamics of generating error-correcting stable states. Unlike existing neural memory models, which exhibit a memory cliff, the circuit (which we call Vector-HaSH, Vector Hippocampal Scaffolded Heteroassociative Memory) exhibits a graceful tradeoff between number of stored items and detail. Next, we show that structured internal scaffold states are essential for constructing episodic memory: they enable high-capacity sequence memorization by abstracting the chaining problem into one of learning low-dimensional transitions. Finally, we show that previously learned spatial sequences in the form of cortico-hippocampal location-landmark associations can in turn be used as larger scaffolds and associated with neocortical inputs for a high-fidelity one-shot memory, providing the first circuit model of the “memory palaces” used in the striking feats of memory athletes.

## Introduction

As we navigate through life, the hippocampus weaves threads of experience into a fabric of episodic memory. Cross-linked by various contexts, this fabric allows us to revisit scenes and events from only a few cues, like Proust’s famous madeleine^1^. Such cue-driven recall makes memories available in ways relevant to making inferences in the present and planning for the future. The hippocampal complex is responsible for this functionality^2–5^, but it is unclear exactly how the architecture and representations of the hippocampal formation and the adjoining entorhinal cortex and other cortical regions enable it.

The representations and dynamics in substructures of the hippocampal complex have been studied extensively^6–26^, and experimental findings combined with models and model testing have resulted in striking progress in our understanding of local circuit mechanisms^18,27–55^. These works put us in an excellent position to now build our understanding of the combined system, on how the substructures work together to subserve robust, efficient, and high-capacity associative memory storage and recall. A particularly intriguing question centers on the dual role of this structure: the hippocampus underlies both episodic memory and spatial memory. Why are these two forms of memory co-localized? Episodic memory, or the storage of new autobiographical experiences, is famously compromised by damage to the hippocampal complex^56–58^. Spatial memory, which refers to our ability to remember and navigate the layout of our physical environment, involves the hippocampus. Place cells in the hippocampus fire at a particular location in a particular environment and context^59,60^. Grid cells of the entorhinal cortex play a complementary role: they generate invariant representations for spatial displacements across environments^36,61–64^, in the form of triangular grid-like firing patterns^61^. Thus, entorhinal grid cells are hypothesized to generate a generalized coordinate system, while hippocampal cells encode specific bindings of features and locations. Both episodic and spatial memory can be accumulated and accessed over a lifetime without major interference, despite the small size of the hippocampus relative to cortex.

The dual memory function of the hippocampus might be understood by three distinct (but non-exclusive) hypotheses. The first is that the circuit’s architecture and representations are optimized for spatial memory, prioritizing the importance of spatial information content for survival – such as details about where we found certain foods and dangers^65^. In this view, episodic memory is an augmentation of the system and representations are not optimized for it. The second is that the circuit is optimized for episodic memory, with spatial coordinates represented primarily because space is a stable and useful index into episodic memory^66^. The third hypothesis is that the circuit doesn’t merely encode spatial and non-spatial episodic information side-by-side or one in the service of the other. Rather, all of its highly structured architectures, representations, and dynamics are equally optimal for both functions, regardless of whether the memory in question involves space. Thus the third hypothesis is that the *abstract* low-dimensional states in the circuit that might be interpreted as spatial are equally critical scaffolds for linking together (potentially entirely non-spatial) elements of an episodic memory^66–73^.

In this work, we build and extensively characterize a new neocortical-entorhinal-hippocampal memory model. We find that this circuit excels at three kinds of memory: for individual inputs (item memory), for spatial mapping (spatial memory), and for sequences (episodic memory). One of the most interesting properties of the model is that the seemingly spatial representations of the grid cell circuit, specifically the low-dimensional and vectorial nature of the code, play a critical and distinct role even for completely non-spatial episodic memory. In other words, our model supports the third hypothesis about the equal relevance of a shared set of structured representations in the hippocampal complex for spatial and episodic memory functions.

The highly constrained architecture, neural activations (invariant low-dimensional representation in grid cells), and biologically plausible learning rules of Vector-HaSH enable memory without the full erasure (memory cliff) seen in existing neural memory models when adding inputs beyond a fixed low capacity, Fig. 1a. As we will see, two features are critical to the abilities of the circuit: 1) The factorization of the problem of creating dynamical fixed points (for pattern completion and error correction) from that of content storage. Fixed points are created by a prestructured scaffold of rigid grid cell circuits (in accord with our knowledge of that circuit^36,61–64^) interacting in a fixed and random way with hippocampus, while content is stored by (hetero)association of these fixed points with input data. 2) Mapping transition dynamics in episodic memory to a low-dimensional shift operator acting on the grid states.

**Figure 1.**
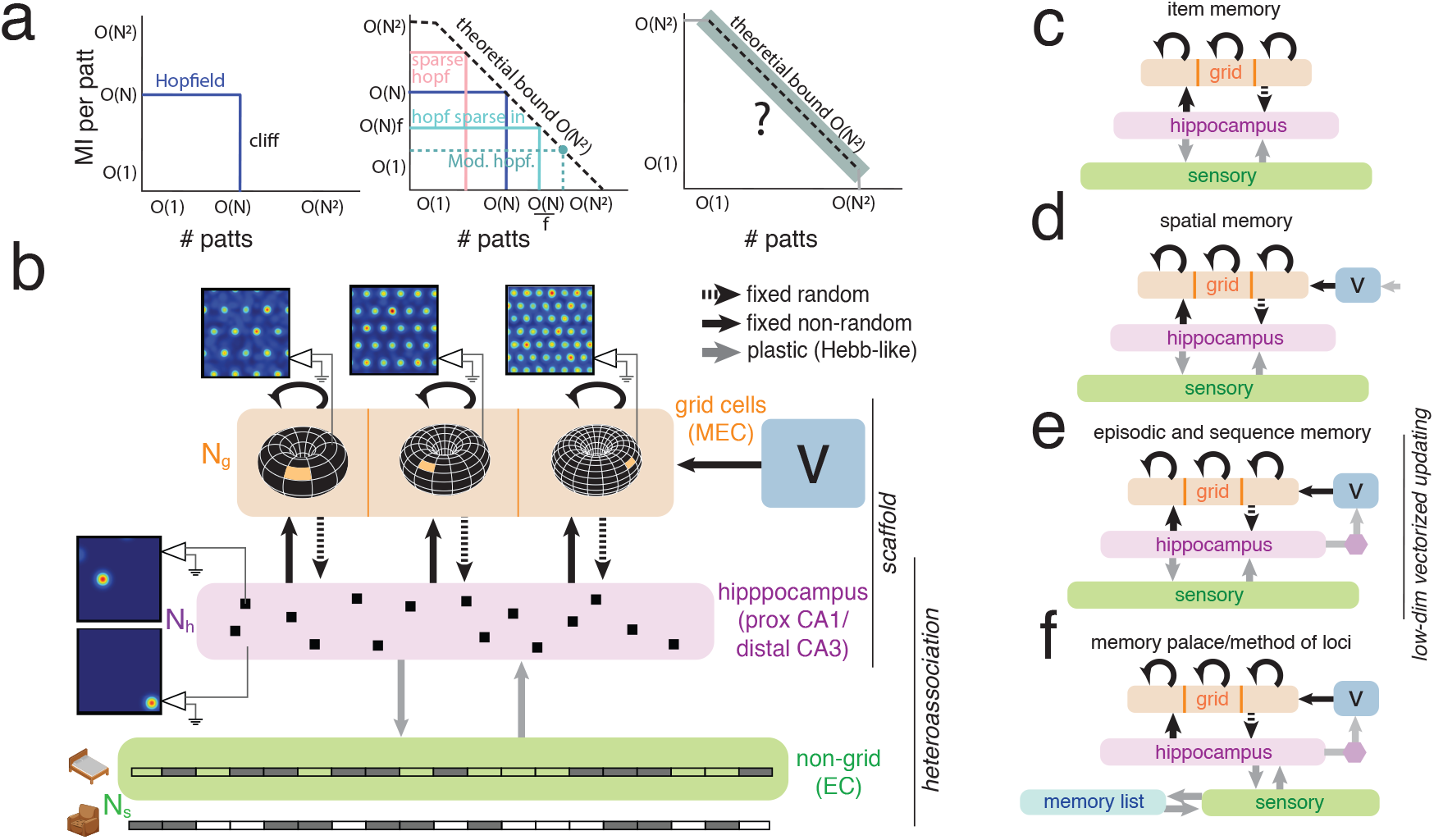
The challenge of biological memory and a biologically informed architecture for general episodic and spatial memory. (a) *Left*: Hopfield networks exhibit a memory cliff: In an ∼ *N*-neuron network, up to *N* patterns (*N* bits each) can be perfectly recovered (recovery quantified through the mutual information (MI) between the stored and recovered patterns, cf. SI Sec. B.2); adding more patterns leads to total loss of all prior patterns. *Center*: All variations of Hopfield networks^74–76^ have a memory cliff. Dashed line: theoretical information bound for neural networks with *N*^2^ plastic synapses^74,75^. *f* = *p* ln(*p*) where *p* is the input pattern sparseness. *Right*: An ideal content-addressable memory would maintain ∼ *N*^2^ bits of total information regardless of the number of stored patterns. No existing model does so. (b) Vector-HaSH consists of multiple grid cell modules (orange) with predefined and rigid connectivity and a low-dimensional velocity or “shift mechanism” (light blue, marked *v*), random fixed projections to hippocampus (pink) with once-learned and fixed projections back to grid cells. This circuit constitutes the fixed “scaffold”. Processed sensory inputs (green) from cortex (including non-grid entorhinal neurons) project to hippocampus with bidirectional connections that are plastic and modified via associative learning. (c-f) Vector-HaSH with small variations or additions to model item memory (c), spatial memory (d), episodic memory (e), and the memory palace circuit for the method of loci mnemonic technique used by memory athletes (f). In the latter, a previously learned spatial memory is repurposed as a scaffold for high-fidelity one-shot memory.

In coding-theoretic terms, for each input pattern, the model assigns a hash code (a unique internal label, given by the internal scaffold state); prestructured recurrent connectivity in the scaffold converts hash states into error-correcting fixed points; and the structure of the scaffold states enables reconstruction of the input patterns with minimal possible interference. The model exploits the fact that grid coding states are ordered within a low-dimensional space to enable efficient sequence memorization through vector transitions. For these reasons, we call our model Vector-HaSH: Vector Hippocampal Scaffolded Heteroassociative Memory.

All code for running the model will be made freely available (upon publication) for others to make and test predictions for future experiments.

## Results

### HaSH architecture for hippocampal associative memory: Factorization of dynamics and content

Our model is based on known and inferred recurrent connectivity between entorhinal cortex and hippocampus^77–80^ and among grid cells in the entorhinal cortex^33^. Processed extrahippocampal inputs enter the hippocampus (Fig. 1b, purple) via direct and non-grid entorhinal inputs (Fig. 1b, green); these inputs carry sensory information from the world, but also internally generated cognitive inputs from other brain regions^66^. The hippocampus also receives inputs from entorhinal grid cells (Fig. 1b, orange). It connects back out to both grid and non-grid cells.

The grid cell circuit consists of multiple grid modules^81^, comprising disjoint groups of cells with grid periods *λ* . Each grid module is assumed to enforce an *invariant* set of low-dimensional states regardless of task. This invariance has been established in an extensive set of studies of the population states and cell-cell relationships of co-modular grid cells across behavioral conditions and states, including navigation in familiar and novel environments, across different spatial dimensions, and across sleep and wakefulness^35,36,62,64^. In spatial contexts, we can describe grid cell modules as coding position as a phase modulo their spatially periodic responses^61,82,83^. In non-spatial contexts, the states of a grid cell module remain the same but can be conceptualized as abstract representations constrained to lie on a 2-dimensional torus.

Connections from grid cells to hippocampus are set as random and fixed. Connections from hippocampus to grid cells are set once (e.g. over development) through associative learning, and are then held fixed. As we will see, the fixed internal grid connectivity and random fixed projections from grid to hippocampal cells are critical for many important properties of the circuit. Connections between hippocampus and non-grid inputs remain bidirectionally plastic and set by associative learning. Activity propagation between regions takes place in discrete time in a sequential manner, a simplification of the idea that oscillations and synaptic latencies gate this information flow. Most simulations involve discrete-valued inputs and grid cell activations, but are shown to generalize to continuous space and activations.

Because the grid cell states are fixed and the grid-hippocampal weights are bidirectionally fixed, we refer to the grid-hippocampal circuit as the fixed *scaffold* of the memory network. This architecture, involving a set of low-dimensional states (grid cell circuit) that is recurrently coupled through high-rank projections to the hippocampus, creates a large bank of well-behaved fixed points, as we will see next. Separately, we refer to the hippocampal-non-grid cortical feedback loop as the *heteroassociative* part of the circuit. In this circuit, a separate set of connections than those generating fixed points heteroassociatively attach sensory data to the scaffold. Unlike standard associative memory models like the Hopfield network^84^ in which the recurrent weights stabilize and associate content directly, here the two are separated: Vector-HaSH *factorizes* recurrent dynamics from content.

We next explore the theoretical and empirical properties of this circuit architecture and its extensions for content-addressable memory in various settings, from spatial to non-spatial memory to sequential episodic memory, Fig. 1c-f.

### Generation of vast library of robust fixed points in an invariant scaffold

The grid cell circuit consists of a few (*M*) independent grid cell modules: the population states of the neurons in each module are constrained to lie on a 2-dimensional torus. Each grid module can express just one state on the torus at a time, independent of the other modules. The *i*^th^ module can take one of *K*_*i*_ states, thus together they express ∏_*i*_ *K*_*i*_ ∼ ⟨*K*⟩ ^*M*^ many, or exponentially many, distinct states. If all co-active grid cells across modules are bidirectionally coupled to a hippocampal cell, that grid-hippocampal state becomes a stable fixed point, enabling intrinsic error correction^83,85^. However, the hippocampus does not possess enough cells to convert each grid state into an attractor in this way.

In the scaffold network, grid cells project with fixed random weights – a high-rank random projection – to hippocampal cells, which threshold and rectify their inputs. The return projection is learned once through simple Hebb-like (associative) learning to reinforce the input grid cell state, then held fixed (Methods).

#### Random fixed scaffold converts exponentially many grid states into stable fixed points

Remarkably, the random grid to hippocampal projections combined with associatively learned return projections in the scaffold converts *all* the grid states into stable fixed points, or attractors (defined as fixed points that correct hippocampal errors of norm 0.25 times the average hippocampal state norm), of the entorhinal-hippocampal circuit, Fig. 2b-c, for a sufficiently large hippocampal network. Adding noise to a hippocampal state derived from any grid state, then running the dynamics of the circuit, exactly restores the correct (denoised) hippocampal state for the original grid state.

**Figure 2.**
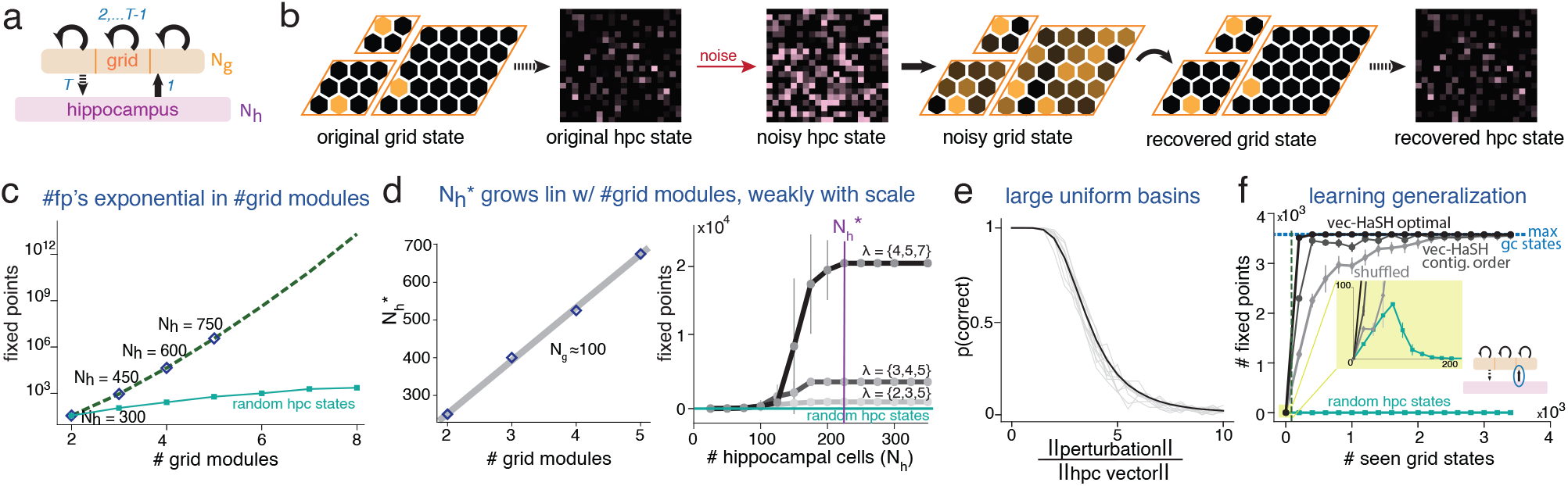
Scaffold generates exponentially many fixed points with large equal-sized basins. (a) The scaffold consists of prestructured grid cells, random fixed grid projections to hippocampus, and fixed non-random associatively learned return projections. 1 · · · *T* indicate the flow of dynamics (order of updating) in the circuit. (b) Left two panels: a hippocampal state that is a projection of a grid state. Next four panels: Initializing the scaffold with a noisy version of this hippocampal state results in a cleaned up state after one round-trip pass through the scaffold. (c) Number of stable fixed points of scaffold dynamics grows exponentially with number of grid modules^82,83^, provided the hippocampus is large enough (diamonds: numerical simulation results; green dashed line: analytical results, derivation in SI C.1). Error bars not visible. Light green: grid-hippocampal weights learned bidirectionally (scaffold not built from fixed random projection of grid cell states). (d) Left: Minimal hippocampal size 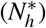 to achieve exponentially many fixed points grows lineary with the number of grid modules. Right: 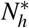 is nearly independent of the grid periods, *λ* . Light green: as in (c). (e) Probability that a noisy hippocampal state is returning to the correct noiseless state, as a function of noise magnitude (divided by hippocampal state norm). Black: average over 3600 fixed points, 100 random noise realizations per fixed point. Grey curves: five random individual fixed points. (f) Strong generalization in setting the return hippocampal-to-grid projections: number of stable fixed points as a function of number of states over which the hippocampus-to-grid weights are learned. Horizontal dashed line: total number of possible grid states. All grid states become fixed points after learning from a near-zero fraction of all states. Black: Vector-HaSH with optimal sequence of seen grid states. Dark gray: generic contiguous sequence of grid phases. Light gray: random set of grid phases. Light green: as in (c). Vertical dashed line: theoretical minimum number of grid states for all grid states to become fixed points.

The minimum number 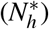 of hippocampal cells required to convert the exponentially many grid states into stable fixed points is small (Fig. 2c). In fact, this number scales only linearly with the number of grid modules (Fig. 2d, left; SI Fig. S1; proof in SI C.1) and is nearly independent of the scale (periodicity) of the grid modules (Fig. 2d, right, SI Fig. S2; proof in SI Sec. C.1). Thus, the number of stable states generated by the scaffold (∼ ⟨*K*⟩^*M*^ ) is exponential in the total number of scaffold (grid and hippocampal) neurons (*N*_*h*_ + *M*⟨*K*⟩ ∼ *M*(*c* + *K*), where *c* is a constant^1^).

If, instead of using random projections from grid states to dertermine hippocampal states, the same set of hippocampal states are randomly assigned to the grid states and the grid-hippocampal weights are bidiretionally set through associative learning for self-consistency, the number of fixed points in the scaffold collapses, (Fig. 2c-d (light green curves; also see SI Fig. S3). A similar collapse occurs if the hippocampal-to-grid weights are randomly set and the grid-to-hippocampal weights are associatively learned for self-consistency, SI Fig. S4.

We find theoretically that there are no spurious stable states in the scaffold, thus the entire hippocampal state space is devoted to forming large basins for the grid cell states. We also find theoretically that the basins are all convex and essentially identical in size across fixed points, SI Sec. C.2, Sec. C.3. We numerically test these analytical results by computing the probability that a noisy hippocampal state will be corrected to the true noiseless state, as a function of noise amplitude, Fig. 2e. We find that noise that is several times larger than the magnitude of the hippocampal state norm is reliably corrected, and that all basins have identical probability curves. This result confirms the convexity and uniformity of the scaffold fixed point basins, and demonstrates the magnitude of robustness.

The construction of exponentially many large-basin fixed points does not violate fundamental capacity bounds for Hopfield-like recurrent networks. By these bounds, a network of 𝒪 (*M*) neurons and 𝒪 (*M*^2^) synapses can support at most 𝒪 (*M*) user-defined points as stable states of the dynamics, or 𝒪 (*M*^2^) bits of information^74,75^, since the fixed points are pre-determined states and cannot be arbitrarily defined. The specific structure of the pre-determined grid states, randomly projected to the hippocampus, creates well-spaced robust grid-hippocampal attractors with large even-sized basins.

#### Strong generalization property of scaffold

The scaffold network possesses another key property dependent on grid coding, which we call strong generalization. The hippocampus-to-grid weights are set by visiting the grid states, then determining the hippocampal states via the grid-to-hippocampal random projection (with thresholding) and applying Hebb-like associative learning. However, learning these weights does not require visiting all ∼ 𝒪 (*K*^*M*^) grid states. Instead, we find that all grid states become stable fixed points of the iterated dynamics after visiting a small fraction of the states, Fig. 2f.

We derive an analytical proof in SI Sec. C.4, Fig. S5 that the number of visited states to make all grid states fixed points is 𝒪 (*MK*_*max*_) states, where *K*_*max*_ is the number of states in the largest module. When *M* and *K* are large, this is a miniscule fraction of all the grid states. In other words, learning the weights to make a very small number of visited grid states stable fixed points of the scaffold results in strong generalization to the entire set of possible grid states, converting them all into scaffold fixed points.

Strong generalization involves visiting a small number of contiguous grid states. Visiting random grid states instead results in some generalization, but requires seeing more states before all grid states become stable fixed points (Fig. 2f). (However, learning from certain special sets of non-contiguous locations can also lead to strong generalization SI C.4, SI Fig. S6.) Further, if we replace the grid states by fixed patterns of matching sparsity as in MESH^86^ (e.g., by shuffling each grid coding state), there is almost no generalization. (proof in SI Sec. C.4). The strong generalization property is only marginally affected by noise during learning, indicating that strong generalization does not require idealized circumstances (cf. SI Fig. S7) and could be achieved in biological settings.

Biologically, the strong generalization property implies that a little spatial exploration in a small environment by a juvenile is sufficient to generate the entire vast set of scaffold fixed points, which can be used for the rest of the lifetime (as described next).

### Content-addressable item memory through heteroassociation of inputs onto scaffold

A content-addressable memory must store and recall arbitrary (user-defined) input patterns based on partial or corrupted patterns. Scaffold states are not themselves memory states because they are not user-defined. However, the scaffold states can be used for content-addressable memory via *heteroassociation* of external cues with the scaffold.

Consider external inputs to the hippocampus, which arrive directly from neocortex^87–89^ or via non-grid entorhinal cells, Fig. 3a (green). We will call all these sensory inputs for short. An incoming sensory input is ‘assigned’ to a randomly chosen scaffold fixed point via Hebb-like one-shot learning between the input and the hippocampal state by a biologically plausible online Hebb-like implementation of the pseudoinverse rule^90,91 2^. The goal of these weight updates is self-consistency: the drive from hippocampus back to the sensory states should attempt to generate the same sensory pattern that activated the hippocampal state. As additional sensory inputs are received, they are assigned to other scaffold states, and the weights between sensory inputs and the hippocampus are correspondingly updated. Scaffold states can be selected in any order for association with the sensory inputs. Patterns can be stored in an online way (patterns presented once, in sequence) or offline (all weights updated at the same time), with no difference in the corresponding weights and thus performance.

**Figure 3.**
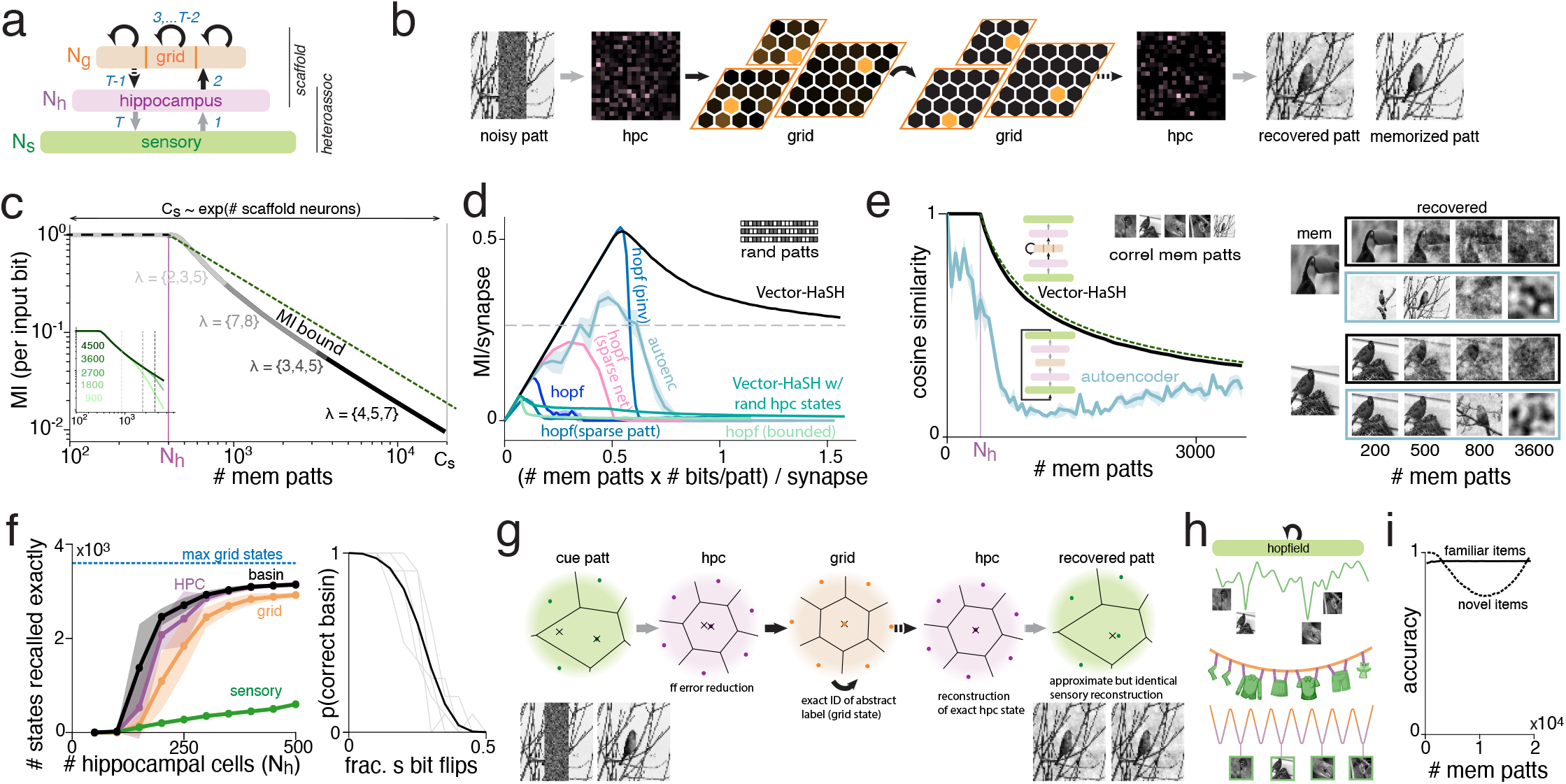
High-capacity content-addressable item memory via heteroassociation with scaffold. (a) Vector-HaSH architecture: scaffold receiving processed sensory input; inputs attached to scaffold fixed points via bidirectional associative learning. Numbers indicate update order for retrieval. (b) Circuit completes a corrupted version of a previously seen sensory input. (c) Mutual information per input bit (MI/number of patterns/bits per pattern) between memorized and recovered patterns versus number of memorized patterns. *C*_*s*_: number of scaffold fixed points; *N*_*h*_: hippocampus size and memory cliff location in Hopfield network. MI per input bit degrades inversely with the number of memorized patterns. Dashed line: theoretical upper bound. Inset: curves for various *N*_*s*_, indicated numbers. See SI Fig. S9. (d) MI (per synapse) versus total number of input bits (per synapse). Even asymptotically, Vector-HaSH recovers a constant MI per synapse (dashed gray line), scaling as the theoretical optimum; for other models, MI per synapse drops to zero as the input information grows. (e) Performance comparison of Vector-HaSH and a fully-trainable autoencoder of matched architecture and size, with identity output-to-input tail-biting weights. Dashed line: analytical capacity result. Right: reconstructions after variable number of stored patterns (black outline: Vector-HaSH; blue: autoencoder). (f) Left: Number of grid (orange), hippocampal (purple), or sensory states (green) recalled perfectly (0-error) when cued with corrupted sensory inputs (2.5% noise). Beyond a threshold *N*_*h*_, most grid and hippocampal states are exactly recalled; sensory state recall is partial and improves continuously with *N*_*h*_. However, the recalled sensory state falls into the correct basin (black curve) when grid and hippocampal recall is accurate. Right: Probability that the recalled sensory state falls in the correct basin versus number of bit flips in cue. (Black: average across sensory patterns; gray: individual patterns; result is for 500 memorized patterns). (g) Schematic of recovery in sensory, hippocampal and grid state spaces. Left to right: Cued and recovered patterns (bottom) are points (top, x’s) in the state spaces (dots: noiseless stored pattern states; lines separate different pattern basins). (h) Energy landscape schematic: Top: Hopfield network minima width, depth, and positions depend on pattern content. Vector-HaSH’s scaffold generates well-spaced minima, with arbitrary content “hooked” on, like a clothesline onto which any clothing may be hung. (i) Accuracy of a simple decoder of familiar and novel patterns based on deviations from the narrow distribution of mean hippocampal firing rates for familiar patterns. In (c-f) errorbars (shaded) are SD over 5 runs.

Once acquired, memories are reconstructed from partial sensory cues as follows: these inputs drive a hippocampal state, which then settles toward a valid fixed point through hippocamapl-grid updating via the scaffold dynamics; finally, the sensory is reconstructed via the hippocampal-to-sensory weights. Vector-HaSH thus behaves as a content addressable memory (CAM) network, Fig. 3b.

#### Associative memory continuum via graceful item number-information tradeoff

Memory recall is perfect up to *N*_*h*_ memorized patterns (the circuit recovers all *N*_*s*_ bits per pattern correctly, corresponding to MI/input bit = 1, where *N*_*s*_ is the size of the sensory input). Beyond *N*_*h*_ stored input patterns, instead of a memory cliff the recovered information per pattern scales inversely with the number of patterns: there is a graceful tradeoff^92^ between number of stored patterns and recall richness. The mutual information (MI) per input bit between recalled and stored patterns remains parallel to the theoretical upper bound (the square of the number of synapses) even as the network is loaded with a number of patterns beyond *N*_*h*_, up to the number given by the exponentially many scaffold fixed points, Fig. 3c. Increasing the sensory input dimension itself does not decrease the information per input bit recovered by the network, because as the pattern size grows, so do the number of heteroassociation weights for storing the additional information Fig. 3c (inset). We call the smooth tradeoff of information per pattern with number of patterns a memory “continuum”, in contrast to the memory cliff of other CAM models^86^.

#### Comparison with existing memory models

We can compare the performance of Vector-HaSH with Hopfield models of various varieties: the classical version, with pseudoinverse learning, with sparse weights, or with sparse patterns. To fairly compare across models, we normalize by the number of synapses in each model, thus using a MI per synapse metric. Up to a threshold number of patterns (linearly proportional to network size), all models perfectly recover their memories. In Vector-HaSH, each synapse conveys a finite amount information per synapse, approaching a fixed constant value per synapse regardless of the number of stored patterns (up to the scaffold number capacity, *C*_*s*_). The recovered information per synapse approaches a constant value up to the scaffold number capacity, Fig. 3d. By contrast, beyond the threshold number of patterns, Hopfield networks exhibit a “cliff” such that the mutual information between recalled and memorized patterns drops to zero for all patterns, including those previously memorized (Fig. 3d). Other memory models also exhibit a similar cliff^93–99^, or can only store a specific number of patterns for a fixed network architecture^76^.

Once all scaffold states have been associated with an input there are three possibilities. The first is that the memory is saturated and no further patterns can be stored. The second involves rewriting a specific memory with the new input, either randomly or based on sensory overlap. The third involves gradual decay of all heteroassociative weights so that older memories are lost and those scaffold states are identified for reuse. The third possibility leads to an incorporation of forgetting within Vector-HaSH, with the oldest memories forgotten first, similar to a recency effect^100^.

#### Comparison with end-to-end trained deep networks

Vector-HaSH can be unfolded for interpretation as an autoencoder^99^, Fig. 3e (left, architecture schematic; black arrows are fixed weights, gray are learnable weights). However, it is a highly constrained one: encoding in the bottleneck layer is fixed, with predefined recurrent dynamics within the layer. The weights from the encoder to the bottleneck and bottleneck to decoder layers are fixed, and all remaining weights are set through biologically plausible associative learning. For comparison, we consider an unconstrained autoencoder of the same dimensions with fully trainable gradient-optimized weights learned end-to-end (via backprop), Fig. 3e (left, architecture schematic), with the addition of a tail-biting connection (identity weights from autoencoder output to its inputs) to enable iterative reconstruction^99^ (Fig. 3e (left)). Strikingly, Vector-HaSH substantially outperforms this autoencoder despite the latter’s much larger potential flexibility, Fig. 3e, mirroring the significant advantage of MESH over the same autoencoder^86^. The tail-biting autoencoder exhibits a memory cliff (Fig. 3e, left); example reconstructions of a sample pattern, after storage of increasing numbers of patterns illustrate the continuum aspect of Vector-HaSH’s memory (graceful degradation of pattern recall, images in black box, Fig. 3e, right) and the cliff of the tail-biting autoencoder (sudden switch in what is being recalled, images in blue box, Fig. 3e, right). SI Fig. S10 shows that Vector-HaSH also outperforms both tail-biting and non-iterated optimized autoencoders when cued with noisy sensory cues.

In sum, the fixed attractor states in the scaffold appear to provide a key inductive bias for robust high-capacity memory that learning with backpropagation on an unconstrained architecture cannot find or achieve.

#### Mechanisms of the associative memory continuum

As we have seen, when a partial or corrupted sensory state is presented to Vector-HaSH, it retrieves an item from memory with the precision of recall depending on the number of stored patterns. We probe the dynamics of the circuit to understand its performance. As *N*_*h*_ is varied (with a fixed number of stored patterns), reconstruction precision is different in different parts of the circuit (Fig. 3f, left): grid and hippocampal states are recalled exactly almost always (beyond a threshold number of hippocampal cells). The sensory state is usually recalled only approximately, with a continuous dependence on the size of the hippocampal area. Even when sensory reconstruction is approximate, the retrieved sensory state falls in the correct basin (the Voronoi region corresponding to the memorized sensory pattern, Fig. 3f, left, black). This is true even when the fraction of errors in the cue is large (Fig. 3f, right, SI Fig. S11).

Consider the dynamics of the network when cued by a highly corrupted memory pattern or by the true memory pattern itself (Fig. 3g, left). The projection onto the scaffold via the sensory-to-hippocampal weights maps the input closer to the appropriate scaffold state (Fig. 3g, green to pink), and scaffold dynamics recover the exact scaffod state (Fig. 3g, pink to orange to pink) even deep in the memory continuum, which we show analytically in SI Sec. D.1. This happens because of the fixed-point dynamics of the scaffold, and their large-basin property. Finally, the hippocampal to sensory projection reconstructs an approximate sensory pattern corresponding to the retrieved scaffold state, with the approximation degrading with interference in the heteroassociatively learned weights across patterns. The reason for this continuous variation is because the reconstructed readout from the stabilized scaffold fixed points happens in a single feedforward pass rather than via iteration, SI Sec. D.2 (see also SI Sec. D.5 for an equivalent analysis for Hebbian learning). In other words, the factorization of fixed point creation from content storage enables both recursive error correction/pattern completion and the graded memory precision behavior of Vector-HaSH.

An interesting and important (as we will see when modeling memory palaces) property of Vector-HaSH gleaned from this dynamics is that though inexact, sensory recovery is reliable: all cues that fall into the same basin converge to a common recovered state (Fig. 3g, right): both noiseless and corrputed patterns map to the same point in sensory space, some distance from the memorized pattern state. As the number of stored patterns is varied, this recalled sensory state typically moves further from the stored pattern within the Voronoi cell. In other words, the *precision* of sensory reconstruction systematically decreases with the number of stored patterns — this accounts for the memory continuum — but the reconstruction is *reliable*: regardless of the cue (which might be noiseless or corrupted), the reconstructed pattern is the same.

We prove in SI Sec. D.2 that associative content-addressible memory with perfect recall for the first *N*_*h*_ states and a near-optimal precision-pattern number tradeoff are possible when the set of grid-driven hippocampal states are constructed as a random projection from grid cells after passing through some nonlinear transformation (in fact, almost every nonlinear transformation is sufficient, without any fine-tuning, SI Figs. S13,S14). In sum for item memory, conventional (e.g. Hopfield) autoassociative memory networks perform poorly because their fixed point locations, basin widths and depths are governed by pattern content, leading to highly uneven and small basins sizes with many spurious minima. In Vector-HaSH, the landscape is set by the scaffold, which has large and well-spaced basins, and content is simply “hooked” onto these prestructured states. The analogy is with a clothesline (the scaffold), to which any clothes (sensory patterns) can be attached (via heteroassociation), Fig. 3h.

### High-capacity recognition memory

Hippocampal states for familiar patterns are constructed from a fixed nonlinear projection of valid grid states (whose statistics remain fixed: the number of modules and the number of activated cells in each module are the same for every valid pattern), thus they are drawn from the same narrow distribution with highly similar firing rates. When a novel sensory input drives the hippocampal state, its projection into grid cells and their return projection back to hippocampus create a pattern far out of the usual distribution for both areas (SI Fig. S15). A simple decoder with two hidden units and thresholds that are fixed from the time of scaffold formation (they do not depend on any details of the sensory inputs or the number of memories stored; SI Sec. D.6) can compare the mean firing rate of the current hippocampal response with its usual mean rate, and respond when the firing rate deviates in either direction from the usual mean. This simple decoder acts as a reliable detector of novelty and familiarity, Fig. 3i.

This novelty-familiarity classification can be used to determine whether a given sensory input should be cleaned up (pattern completed) to recall the closest stored memory, or if it should instead trigger the formaton of a new memory (association of the input with a fresh scaffold state).

### Spatial memory and inference

We now consider how this circuit performs *spatial* memory. Here, the metric or *vector* ordering of grid states, which was not necessary for pattern memory, becomes critical. When self-motion signals during spatial navigation are allowed to drive transitions between the metrically ordered grid states, Fig. 4a, we find below that the architecture and dynamics of Vector-HaSH support spatial memory without catastrophic forgetting and zero-shot spatial inference along novel paths.

**Figure 4.**
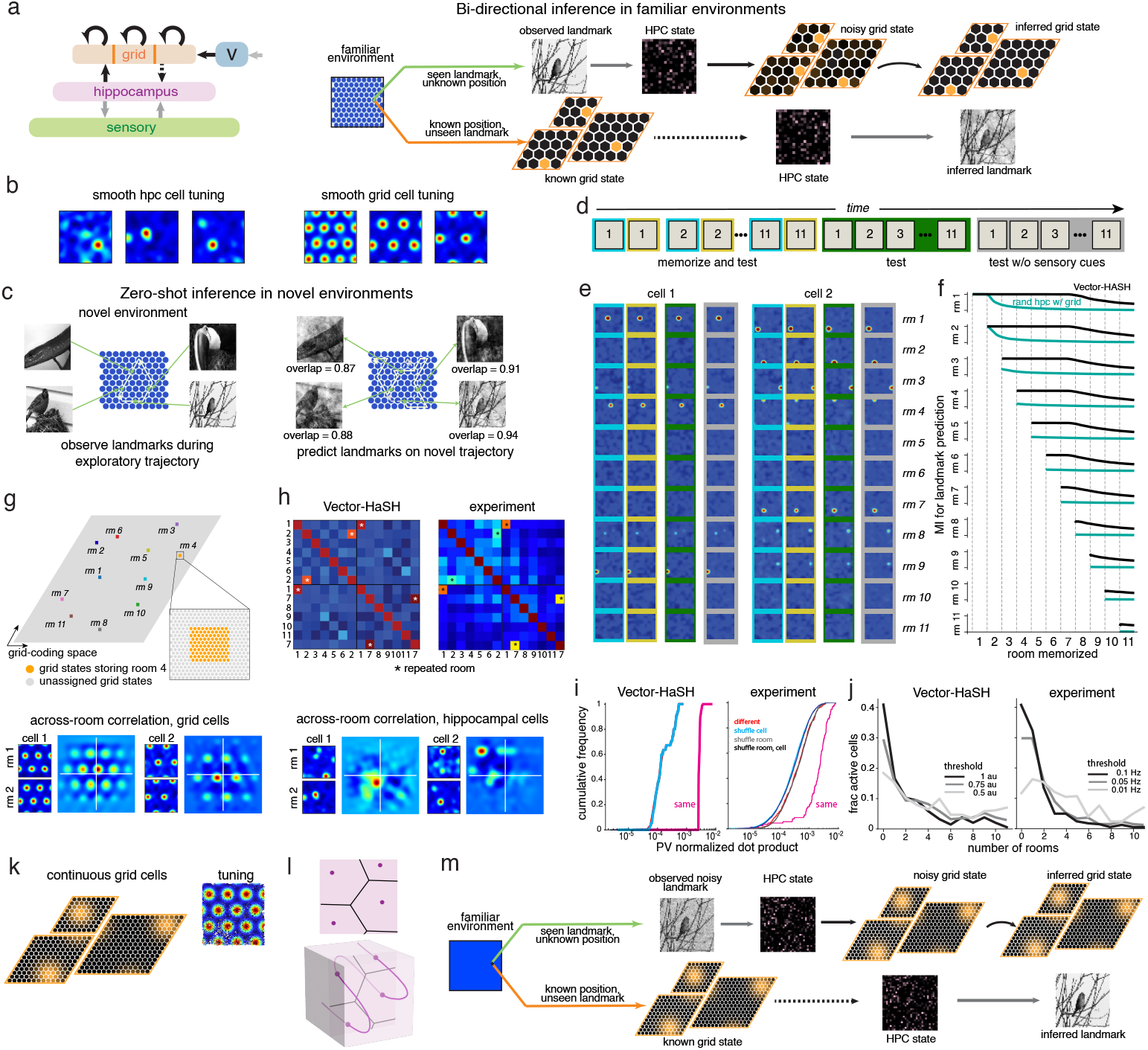
Spatial memory, inference, and lifelong learning without catastrophic forgetting. (a) *Left:* Vector-HaSH with a path integration mechanism such that velocity inputs drive shifts in grid phase. *Right*: In a familiar environment (after associative-like learning), bidirectional inference of position (internal grid states) from sensory cues (top), or prediction of sensory landmarks from a grid state (bottom). (b) For one environment: smoothed tuning of random hippocampal cells and one random cell from each grid module (raw tuning curves in SI Fig. S17). (c) *Left:* First-time traversal of an environment along only shown trajectory, with memorization over the trajectory. *Right:* Vector-HaSH prediction of landmarks on novel path starting from known location. (For realism to fall along memory continuum, Vector-HaSH previously memorized 596 other landmark-scaffold associations (mimicking learning other environments).) (d-e) Sequential learning protocol (d) and spatial maps (two hippocampal cells) (e) for 11 rooms: Vector-HaSH is steered on a random path in each room (blue outline), and memorizes each room once in sequence. Immediately after memorization of room *i*, it is tested there (mustard outline; no learning) on landmark-driven location (scaffold state) inference (thus testing landmark-to-scaffold associations) on a random trajectory. All rooms are re-tested (dark green; no learning) for landmark-driven location inference, then tested for path integration-driven landmark inference in the “dark” (no cues after the starting position). (f) Black: Landmark reconstruction accuracy for room *i* after learning rooms 1 · · · *i*− 1. Recall of room 1 after learning room 11 (top right) is no worse than recall of room 11 immediately after learning room 11 (bottom right). Cyan: baseline model as in blue-green curves of Figs. 2 and 3. (g) Top: Gray diamond represents the set of all grid phase combinations across modules, with side-length given by the exponentially large coding range per dimension of the grid code. Random assignment of the starting grid phase results in non-overlapping grid representations. Bottom: the shift in grid phase going from room 1 to 2 is different across modules (left vs right). (h) Top: Hippocampal population activity similarity matrix across rooms (with repeated exposures to some rooms, white asterisks): Vector-HaSH (left) and experiments^101^ (right). Bottom: total remapping of hippocampal cells across rooms. (i) Hippocampal representation similarity for same versus different rooms and shuffle controls for Vector-HaSH and experiments: representations of different rooms are as orthogonal as the shuffles for both. (j) Distribution of active hippocampal cells in *R* rooms versus *R*: most cells are active in only a few rooms. (k) Vector-HaSH extention to continuous-valued grid cells and locations in space. *Left*: Population vector. *Right*: Raw (unsmoothed) spatial tuning curve of a grid cell from random trajectory. (l) *T*op: Scaffold basins in discrete Vector-HaSH; *Bottom*: depiction of the same for a continuous version (1D line represents 1D continuous represented variable; the actual continuous manifold would be a folded 2D sheet in 3D with basins in higher dimensions). (m) Similar to (a),*right*, continuous Vector-HaSH can perform bidirectional location (grid) inference from noisy landmarks, and landmark prediction from grid states.

In a novel room, grid module phases are randomly initialized at a landmark (a room corner could suffice). Velocity inputs then update the grid phases by path integration^102^. Vector-HaSH learns associations between the grid-driven updating scaffold states and any sensory cues in the room, building a map of the space. This learned map allows for bi-directional recall of grid states from sensory cues and vice-versa (Fig. 4a). If a familiar room is traversed without access to sensory inputs (in the “dark”, or in the interval between landmarks in the light), Vector-HaSH continues to perform path integration to update grid (and thus hippocampal) states. If it then encounters a landmark, Vector-HaSH updates the hippocampal state through the sensory-hippocampal weights (pathway shown in Fig. 3a-b), resetting any path integration errors. Grid cells and hippocampal cells exhibit realistic spatial tuning curves, including the spatially localized and typically single-bump tuning of place cells (Fig. 4b, S16).

After even very sparse exploration in a novel room (Fig. 4c, left), Vector-HaSH is able to predict expected sensory observations when taking an entirely novel route through the room (Fig. 4c, right). This zero-shot spatial inference ability arises from the path-invariance of velocity integration^103–105^: the initial grid phases are updated based on velocity to generate accurate grid states for locations even along novel paths, which can then drive reconstruction of the sensory cues associated with those states.

Next, we consider the ability to sequentially learn a series of different rooms, Fig. 4d. For each room, Vector-HaSH learns a reliable spatial map, as assessed by testing sensory-cued grid state inference (without path integration) right after it has learned that room. The hippocampal maps are distinct and reliably recovered for each room, Fig. 4d-e.

To assess sequential memory and the extent of overwriting of memories of one room by the subsequently learned room(s) (catastrophic forgetting), after Vector-HaSH learned the *i*th room, we tested recall in all prior rooms (Figs. 4d-e. Recall of the hippocampal and grid cell activations (maps) from sensory cues remains unchanged in all prior environments despite the subsequent acquisition of up to 10 new rooms. Notably, there is a total absence of catastrophic forgetting in grid and hippocampal recall. This lack of catastrophic forgetting does not require replay or consolidative associative learning to refresh prior memories. It is due to the specific architecture and exponential scaling capacity of Vector-HaSH, in which random grid phase initializations result in maps that are well-separated in the coding space, Fig. 4g. Next, we assess reconstruction of the sensory state in the “dark” without visible landmarks, where the the initial grid state is specified for each room, and phases are updated by path integration, Fig. 4f. The model is able to recall a large amount of information (one cue at every location in every room) over 11 rooms, though the sensory cues in all rooms are recalled less vividly after learning 11 rooms (rightmost part of all black curves, Fig. 4f) because of the memory continuum regime for sensory recall.

The spatial response properties resemble observed responses in the brain, including the similarity of hippocampal responses for repeated visits to the same room and orthogonal representations of different rooms, Fig. 4i-j. Additional properties of the hippocampal response, including the distribution of probabilities that a hippocampal cell has a field in multiple rooms, matches experimental data, Fig. 4k.

Finally, we implement a continuous-activation version of Vector-HaSH (continuous grid coding and continuous space; Methods). In the continuous limit, the discrete fixed points of the scaffold network become a folded two-dimensional continuous plane of fixed points in the *N*_*h*_ dimensional space of hippocampal states, but the basins remain large (now *N*_*h*_ −2 dimensional instead of *N*_*h*_ dimensional; Fig. 4m illustrates this for a one-dimensional continuous manifold). Qualitatively, the associative robust memory retrieval property of Vector-HaSH carries through, as does bidirectional inference, Fig. 4l-n.

### High-capacity sequence scaffold via vector updating of grid states

Sequence memory is typically modeled with asymmetric Hopfield networks^106–108^, resulting in similar capacity limitations as standard Hopfield networks^74,99,109^. Remarkably, it is possible to construct a massive sequence memory in Vector-HaSH in a similar way as item memory and spatial memory: by factorizing the problem into a high-capacity abstract scaffold sequence and then affixing content via heteroassociation. We first explore how to construct these high-capacity sequence scaffolds, based on the velocity-shift mechanism of grid cells.

Memory networks perform poorly when user-defined patterns determine the attractor states (Fig. 2e-f). The equivalent problem in sequence memory is that user-defined patterns determine and drive the next user-defined pattern: a Hopfield-like auto-associative network with asymmetric weights quickly fails (within ∼ 50 steps) to reconstruct even an approximation of the pattern, Fig. 5b.

**Figure 5.**
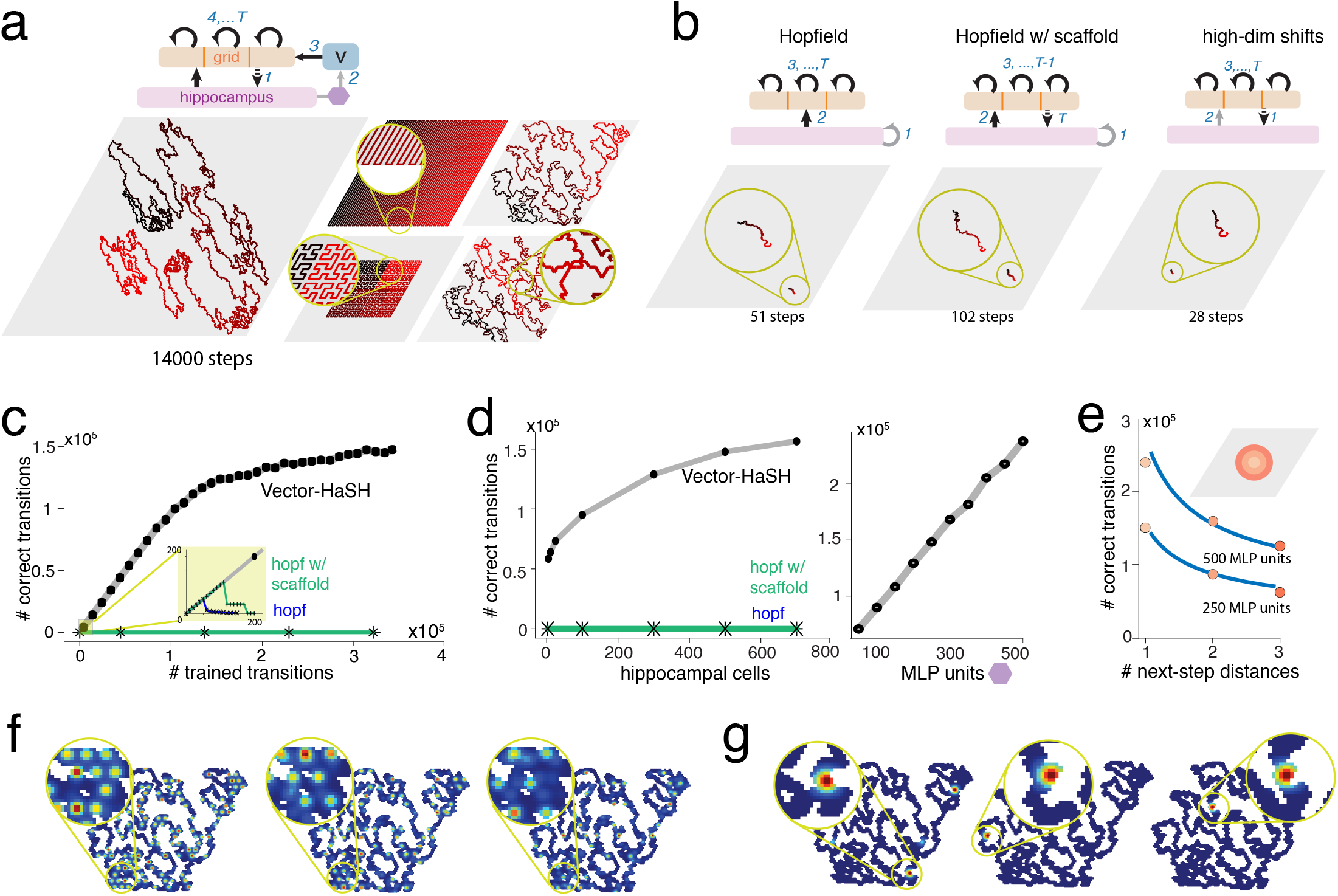
High-capacity scaffold for sequences via low-dimensional velocity update mechanism in grid cells. (a) *Top*: Architecture of the sequence scaffold: transitions of grid states are driven by the hippocampus via the 2D grid cell velocity shift mechanism. A small feedforward network (single hidden layer MLP, *N*_*M*_ = 250; violet) maps the current hippocampal state to a velocity state. *Bottom*: Recall of a 14000-step length self-avoiding random walk (Lévy flight), a hairpin sequence spanning all the grid coding space, a Hilbert curve that spans every point in a subset of the grid coding space, and uniform random walks; curves colored temporally as red-to-black (left-most: parameters as in (b); last four, *λ* = {3, 4, 5} with a 3600 sized state space). (b) Architectures (*top*) and recall performance (*bottom*) of hippocampus as an asymmetric Hopfield network (left), as an asymmetric Hopfield network assisted by the scaffold (center), and as a pure scaffold with hippocampal-to-grid connections driving the next grid state (right). In all cases, network state is visualized via the grid state (*bottom*, a 585 ×585 state space, *λ* = {5, 9, 13} ) and*N*_*h*_ = 500 nodes, and the sequence being memorized is the same as (a)*bottom-left*. Number of steps indicates the longest length of sequence that could be learned by each model before catastrophic failure. (c) Each grid state is assigned a random (one-step) next-state shift direction; Vector-HaSH successfully recalls these state-associated shifts while the Hopfield baselines fail to do so. (d) *Left*: accurately recalled sequence length increases exponentially with the number of hippocampal cells (trained on all 585× 585 random shifts). *Right*: Recalled sequence length increases exponentially with the size of the MLP that maps hippocampal states to a 2-dimensional velocity vector (trained on all 585 ×585 random shifts. (e) Increasing the information load of next-state generation by increasing the range of step-sizes. Blue curves proportional to 1 over the logarithm of the number of distinct possible shift vectors (f-g) Sample grid and hippocampal cell tuning in the abstract 2D space defined by the grid velocities. This space is abstract: it need not correspond to physical space.

We reasoned that if, to a Hopfield-like asymmetrically recurrently coupled hippocampal network, we added bidirectional interactions with grid cell modules as in the scaffold network, it might support high-capacity sequence reproduction by denoising and pattern-completing otherwise inaccurate next-step patterns generated within just hippocampus. Doing so roughly doubled the sequence capacity of the circuit (to ∼ 100 steps), but did not fundamentally alter the scaling of capacity with network size, Fig. 5b.

Next, we reasoned that, in the full spirit of the scaffold network, learning an abstract sequence of scaffold states rather than user-defined hippocampal states might be the solution. Hippocampal states were determined by random grid state projections, and we tested the performance of learning transitions from one abstract grid state to the next via the return projections from hippocampus to grid cells. (The return projections from hippocampus were set associatively to be consistent with the next grid cell state, rather than with the current one.) With the full benefit of the scaffold architecture, the sequence capacity remained qualitatively similar to Hopfield networks, Fig. 5b, with failure within ∼ 30 steps. This happened possibly because even abstract grid states are large and specific activity patterns, which the previous state must sufficiently specify to reconstruct. This failure and hypothesized reason gave us the critical insight that learning the *input to the velocity shift mechanism*, which requires specifying merely a 2-dimensional vector to specify the next grid state given the current one, would minimize the information required from the previous state and potentially alleviate the capacity limitation for sequence reconstruction.

We therefore used the previous hippocampal state to cue the next grid state, but via the drastic dimensionality- and complexity-reduction of the velocity-shift mechanism: the previous hippocampal state specifies a 2-dimensional velocity that signals where to move in the grid coding space for the next state. We built these associations via a simple feedforward network (MLP), Fig. 5a (top) that associated the previous grid state, via the hippocampus, with a 2-dimensional velocity vector. This architecture resulted in the accurate reconstruction of scaffold sequences of 1.4 × 10^4^ states, using the same (small) number of cells in the scaffold network as before, Fig. 5a (left).

To quantify sequence memory, we took a statistical approach (avoiding the need to build specific sequences): we assessed how well the circuit could recall random velocity (shift) vectors assigned to each grid state, Fig. 5c. The circuit memorized and perfectly recalled ∼ 1.5 ×10^5^ state-velocity associations with a scaffold of *N*_*h*_ = 500 and *N*_*g*_ = 275, with grid periods 5, 9 and 13 (and hence a total of ∼ 3.4 ×10^5^ total scaffold states); longer sequences were reproduced with modest decreases in recall performance. The dependence on the number of hippocampal cells is again smaller than logarithmic, similar to scalings for item memory, Fig. 5d (left), while the dependence on the number of units needed to learn the dimension-reducing mapping from state to velocity vectors is linear with a very small coefficient ( ∼ 10^−3^), Fig. 5d (right). .

In sum, recalling a long abstract sequence of grid states can be achieved by converting it into the task of recalling a sequence of simple abstract two-dimensional vectors, each of which points from one grid state to the next. We hypothesize that this enables much longer sequence reconstruction because the information the hippocampus must reconstruct for each step in the sequence is a mere two-dimensional vector, not the much more complex grid pattern state, Fig. 6f. We verify this hypothesis by parametrically varying the amount of information the network must recall at each step to arrive at the next, and quantifying how that affects the recalled sequence length. To parametrically vary information in a 2D velocity, we increased the range of lengths of possible 2D vectors to be recalled. As a result, the recalled sequence fraction decreased systematically (Fig. 5e; the theoretically expected scaling, in which sequence length decreases as the inverse of the number of bits required to specify the next step, is shown in blue). Thus, constraining the sequence recall dynamics to a low-dimensional manifold where only low-dimensional velocity tangent vectors rather than the manifold states themselves must be reconstructed, results in vast increases in sequence length.

**Figure 6.**
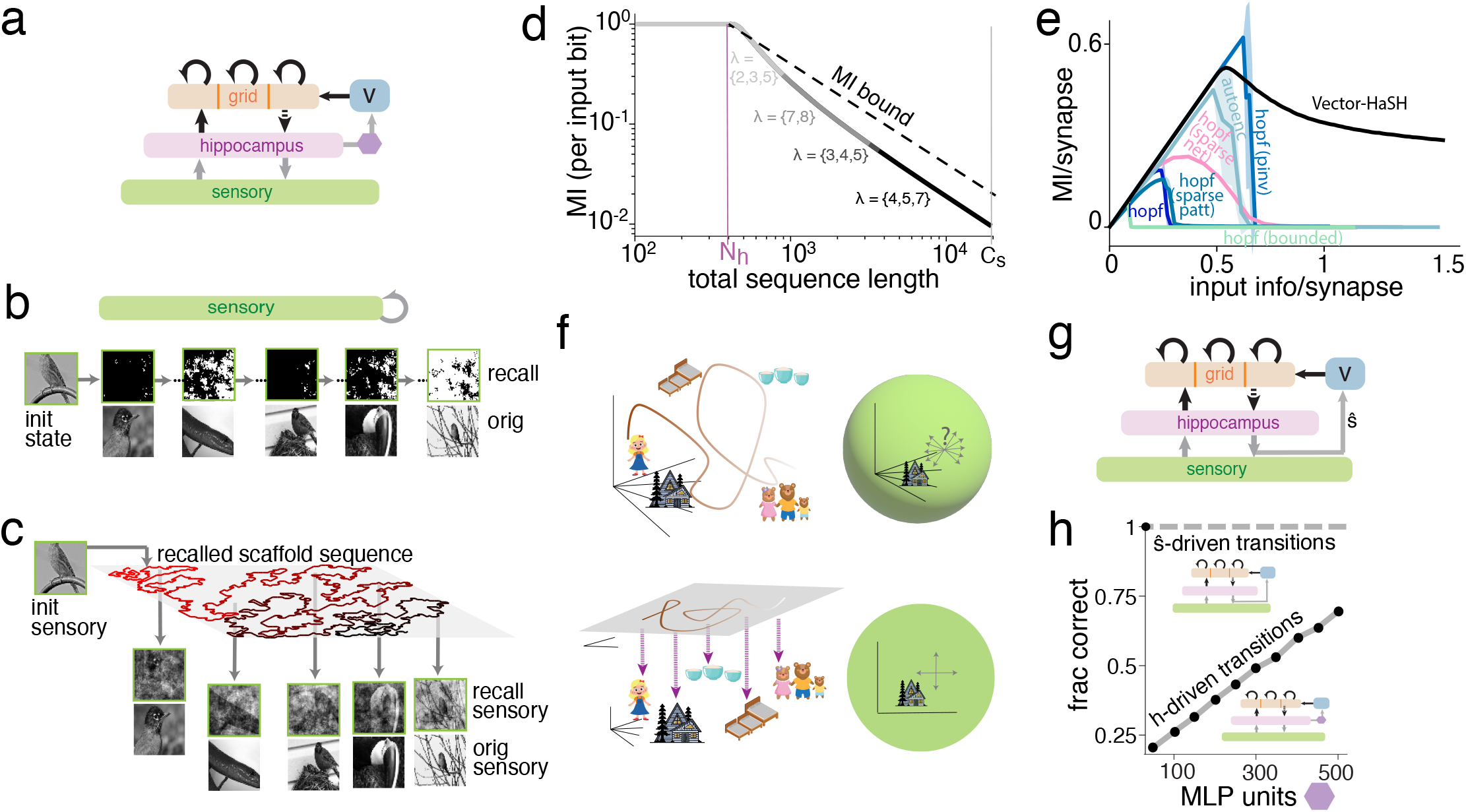
High-capacity episodic (sequence) memory via heteroassociation with sequence scaffold. (a)Architecture of Vector-HaSH for episodic memory. Hippocampal states determine a low-dimensional (2D) shift vector 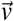 for grid cells, via a decoding network (3-layered MLP, purple hexagon). (b-c) Example recalled trajectory and sensory states in an asymmetric Hopfield network (b) and Vector-HaSH (c) with the same number of synapses (d) for `bw-mini-imagenet` sensory inputs (see Methods; periods {3,4,5}, *N*_*g*_=50,*N*_*h*_=400,*N*_*s*_=3600 for Vector-HaSH, and *N*_*s*_=3600 for Hopfield). (d) MI (per input bit) between stored and recalled sensory states as a function of sequence length for Vector-HaSH, similar to the item memory curve of Fig. 3c. (e) MI (per synapse) for sensory recall as a function of episode (sequence) length when storing episodic memories, across models. (Sensory states in d-e are random binary patterns.) Vector-HaSH asymptotically approaches a constant amount of information per input bit. (f) *Top*: In conventional sequence memory models, the previous state must reconstruct the entire content of the next state. *Bottom*:In Vector-HaSH, the previous state must reconstruct a mere 2-dimensional vector, requiring far less information in the recurrent loop. (g) Alternative architecture: the reconstructed sensory state (*ŝ*) determines the 2-dimensional shift vector 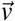 (this resembles route learning, SI Fig. S18). (h) While the accuracy of the hippocampus-driven velocity shift circuit depends linearly on the size of the decoding network (MLP), the accuracy of sensory recall-driven velocity shifts can be consistently high without any MLP because of the size of the sensory network and because sensory recall is reliable even when inaccurate (Fig. 3g). As in Fig. 5d, the fraction of correct shifts is computed on a mapping from *all* scaffold states to random shifts.

Because velocity-driven transitions are 2-dimensional even if the represented space is not, we can plot the grid and hippocampal states as a function of these 2-dimensional transitions, and obtain periodic grid responses, Fig. 5f; hippocampal cells exhibit more-sparse and more-localized tuning than grid cells, Fig. 5g.

In sum, the path integrability of the grid cell code can support not only highly efficient spatial inference and mapping but also a scaffold for sequence memory, even if the sequences do not involve physical navigation in real spaces.

### Episodic memory

Just as high-capacity item memory involved hooking content via heteroassociation onto a fixed-point scaffold, Vector-HaSH supports high-capacity episodic memory by hooking external inputs experienced over an event onto a sequence scaffold, Fig. 6a. The abstract scaffold sequences described above can be formed and learned concurrently with the learning of the heteroassociative weights linking the sensory input to the scaffold. Alternatively, inputs could be affixed to a pre-existing scaffold sequence.

Our control models are Hopfield networks with asymmetric weights and tail-biting autoencoders trained on a sequence of sensory inputs^99^, Fig. 6b. Within very few steps, recall of these models diverges from the trained state sequence, so that recalled states do not even approximately resemble the trained states (the sequential version of the memory cliff).

In Vector-HaSH, sensory states are recalled with varying accuracy, depending on the total amount of stored sequence content, without a memory cliff, Fig. 6c (images). Over the long trajectory shown, the internal grid sequences are recalled with essentially perfect fidelity while sensory states are also recalled perfectly in terms of identity, but recalled only approximately in content (as for item memory, Fig. 3f). This follows from the model architecture (Fig. 6a) and sequence scaffold results (Fig. 5): even when the scaffold is coupled to the sensory network, grid sequences are generated entirely within the scaffold and thus do not depend on the fidelity of sensory reconstruction.

Quantitatively, the quality of sensory recall (MI per input bit) degrades as the total sequence content grows, Fig. 6d, but the information per synapse remains finite, approaching a constant asymptotically, Fig. 6e. By contrast, the MI per synapse drops to zero for other models. These results closely resemble the item memory results above.

The conceptual basis for the failure of conventional sequence memory models is that the previous state and its recurrent projections must carry all the information in them to reconstruct the full N-dimensional next state; while in Vector-HaSH, the previous state and its recurrent projections must reconstruct merely the next 2-dimensional velocity vector. The N-dimensional vector is then reconstructed via the feedforward projection to the sensory areas.

An alternative Vector-HaSH architecture for sequential or episodic memory is one that is not fully factorized: next-step velocity is driven by the previous *recalled* (not actual) sensory state instead of the recalled hippocampal state (Fig. 6g, SI Fig. S18). This alternative works equally well, even in the memory continuum where sensory state recall is only approximate. Because the velocity transition is based on the reconstructed sensory state rather than the sensory cue, and sensory reconstruction is reliable even when approximate, a consistent mapping is possible. In fact, the mapping from sensory state to velocity in this architecture does not require an MLP because of the large number of sensory cells (Fig. 6g-h, proof in SI Sec. D.7). Thus, the fidelity of sequence transitions in this alternative model do not, as in the original sequence scaffold model, depend on MLP size, Fig. 6h.

For episodic memories with content that is not explicitly spatial, the scaffold trajectory can be arbitrarily chosen — in our numerical simulations examining the maximal extent of sequential memory in Vector-HaSH (Figs. 6e-f), we choose a space-filling “hairpin” trajectory in scaffold space.

In sum, the hippocampal-entorhinal circuit in Vector-HaSH is able to store episodic memory of arbitrary input sequences (or several such sequences) with high capacity and maintains a total amount of information that scales in the same way as the theoretical bound, regardless of the total length of stored sequences. It does so by exploiting the low-dimensional vector updating property of grid cell states, even in the absence of any spatial content in the memory.

### Vector-HaSH reproduces multiple aspects of entorhinal and hippocampal phenomenology

Vector-HaSH exhibits considerable flexibility as a general-purpose memory circuit. At the same time, it exhibits several properties of grid and place cells.

Because it is based on continuous attractor grid modules, it generates similar predictions about invariant structure. A prediction of continuous attractor grid cell models that holds for grid cells in Vector-HaSH is that from the very first part of exploration in a novel environment, grid cells exhibit bump-like activations, Fig. 7a (top); and when the trajectory is sufficiently dense to see a pattern, the first-observed activity bumps align with the bumps of the formed pattern, Fig. 7a (bottom). Next, grid cells in Vector-HaSH demonstrate grid-like responses (at least over short time-windows, where path integration errors do not blur out the pattern) in environments without any sensory cues (“dark” exploration), Fig. 7b (left). These observations are consistent with experiments^110^ and contrast with alternative models for the hippocampal complex that require multiple traversals of an environment and the presence of external sensory cues before grid cell-like tuning can emerge^43,51^.

**Figure 7.**
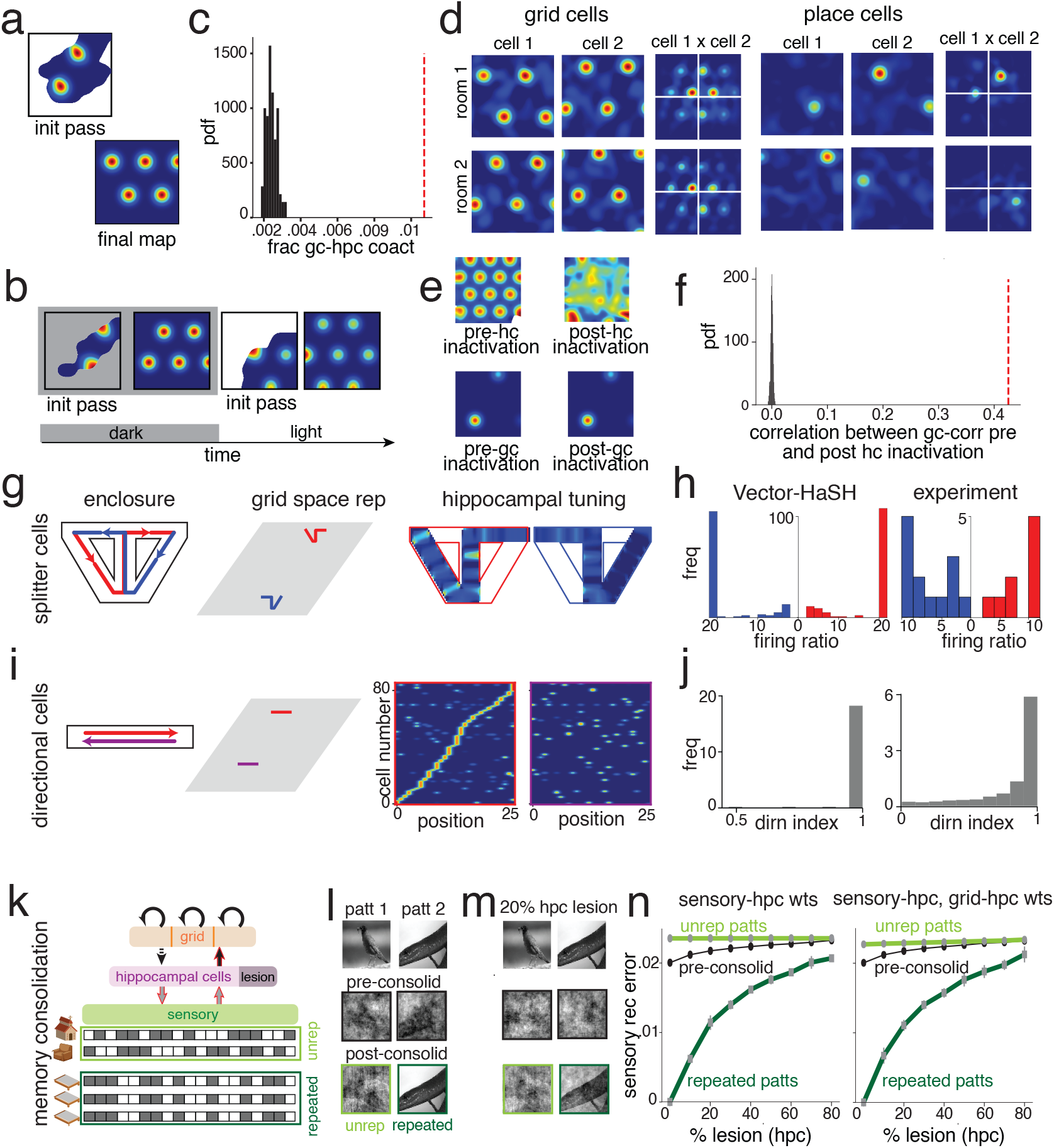
Vector-HaSH reproduces multiple aspects of entorhinal and hippocampal phenomenology. (a) Grid fields are present in first steps in an environment (*top*), consistent with the map formed after fuller exploration (*bottom*). (b) Dark to light (cues provided): grid responses rapidly remap (shift phase). (c) Grid field locations correlated with place field locations across environments (red), relative to shuffled responses. (d) The tuning curves of grid cells shift across environments (*left*), but cell-cell relationships (phase offsets) of co-modular neurons are preserved (closest peak to white cross-hairs). Not so for hippocampal cells. See also SI Fig. S21 for preservation of grid cell-cell relationships across environments of different dimensionalities. (e) Spatial tuning of grid cells (*top-left*) is lost after hippocampal inactivation (*top-right*) when velocity estimation is noisy, however (*bottom*) co-modular grid cell-cell relationships (red line) are preserved (black: shuffled cell indices). (f) Spatial tuning of hippocampal cells (*bottom-left*) is preserved after grid inactivation (*bottom-right*). (g) *Left*: Alternating T-maze task, with two trajectory types: from-left-to-right (red) and from-right-to-left (blue). Contextual remapping to distinct regions of grid space (*middle*) yields hippocampal remapping: tuning curve of hippocampal cell in the two contexts (*right*). (h) Hippocampal firing selectivity ratios for central stem on left vs. right trajectories: Vector-HaSH (*left*) and experiments (*right*)^119^. (i) *Left*: Directional trajectories on a 1D track. *Middle* panel as in (g). *Right*: Hippocampal cells ordered by location on track for rightward(red box) and leftward (second box) runs. (j) Directionality index (normalized metric for neural activity difference on opposite direction runs): Vector-HaSH (*left*) and experiments (*right*)^120,121^: a majority of cells are directional. (k) Memory consolidation in Vector-HaSH: repetition of some sensory patterns drives increments in the strengths of the learned weights (gray arrows outlined in red). (l) In the continuum regime, memorized patterns (*top*) are recalled only approximately (*middle*); consolidation of pattern 2 but not 1 leads to much richer recall of pattern 2 (*bottom*). (m) A 20% hippocampal lesion leads to deterioration of all patterns without consolidation (*middle*), but consolidated patterns are protected (*bottom*). (n) Recovery error versus lesion percentage without consolidation (black) and after consolidation for unrepeated (light green) and repeated (dark green) patterns.

The grid-hippocamapal-sensory interactions in Vector-HaSH permit rapid cue-based resetting of the state of the circuit: After measuring grid responses briefly in a dark environment, if the lights are turned on to reveal a familiar environment with visible orienting cues, the sensory-to-hippocampal weights rapidly reset the hippocampal (and hence grid) states within one cycle of information flow through the loop, Fig. 7b (right), similar to experimental work that reported rapid resetting upon addition of sensory input^110^.

Vector-HaSH recapitulates statistical properties of correlations between grid and hippocampal fields between environments^51^, by virtue of the contribution of grid inputs to hippocampal cells. We simulated Vector-HaSH in a pair of environments and measured the fraction of grid cells with fields that overlapped with place fields across both environments (dashed line, Fig. 7c); this fraction was significantly larger (p-value of 0.0) from the fraction obtained if the place fields were randomly shuffled, maintaining their statistics (black distribution, Fig. 7c). This phenomenon, that a place field is more likely to recur at a location overlapping a grid cell’s field if they had overlapping fields elsewhere, is consistent with experimental data^51,111–113^and with models in which grid and place cells are synaptically coupled with grid cells contributing to the formation of place fields^50,51,114,115^.

A fundamental prediction of the continuous attractor model of grid cells, repeatedly verified through analysis of experimental data^34,36,63,64^, is that cell-cell relationships between co-modular grid cells should be preserved across states and environments, and the population states of grid cells should lie on a torus. Grid cells in Vector-HaSH possess this property: two grid cells each change their tuning curves across environments, however their relationship (relative phase) is preserved across environments, Fig. 7d (left), SI Fig. S21. By contrast, hippocampal cells have globally remapped and their cell-cell relationships are not preserved, Fig. 7d (right). Like continuous attractor models of grid cells, from which Vector-HaSH inherits several of its invariance properties, the internal grid states and thus grid cell phase relationships are predicted to be invariant to varying environment dimension (1D, 2D, 3D), geometry, and topology, even when the tuning curves of individual cells become deformed or otherwise changed^34^. We demonstrate this for a 1D linear track and a 2D square room in SI Fig. S21.

Though grid cell projections determine hippocampal states in Vector-HaSH, there is also a strong reverse influence of place cells on grid cells. When place cells are lesioned in the presence of noisy velocity inputs to grid cells, grid cell spatial tuning is destroyed (Fig. 7e,*top*), a prediction that is consitent with experimental results^22,116^. At the same time, they maintain their cell-cell relationships (Fig. 7e *bottom*; details in *Methods*), again consistent with experiments^116^. Thus, our model predicts that place cells are critical for reliable spatial tuning of grid cells. In contrast, place cells can maintain their fields without disruption when grid cells are lesioned, assuming the presence of sensory inputs throughout an environment (Fig. 7f)^117^. In short, the circuit uses all means at its disposal to estimate position: based on velocity (via grid cell velocity integration), based on external spatial cues (via hippocampus), both, or either; it reconciles questions of whether place cells emerge from grid cells or vice versa^118^.

Vector-HaSH can reproduce the the phenomenology of splitter cells^119–122^, hippocampal neurons whose firing in an unchanging environment varies based on context, e.g. varying with start and target locations, or for random foraging versus directed search. To model splitter cells, we assumed that context changes trigger re-initialization (remapping) across grid modules. At reinitialization, a randomly selected set of initial grid phases is assigned to each context, and sensory-hippocampal-grid associations are built while traversing the environment in this context. When the agent returns to this context, the stored associations are recalled, similar to^123^.

Under these assumptions, we simulate experiments in which splitter cells have been observed. Simulated runs on the central stem of a T-maze for future left- or right-turns are distinct based on the context of arriving from the right- or left-side of the environment. These therefore map to distinct regions of grid space Fig. 7g, predicting grid cell remapping (differential shifts of grid phase across modules, SI Figs. S26, S27) and total hippocampal remapping, Fig. 7g. The resulting distinct place cell representations in the central stem of the T-maze are consistent with splitter cells; the grid cell results can be tested in future experiments.

The same is true for left-versus right-runs on a one-dimensional track (Fig. 7i), tree-shaped mazes, clockwise versus counterclockwise runs on a closed path, and radial mazes, SI Figs. S22, S23,S24,S25.

In other words, contextual grid cell remapping is predicted to yield a variety of cell behaviors, from splitter cells to directionally selective place cells. The ratios of splitter to non-splitter cells and of direction-dependent to non-directionally tuned cells were similar to the ratios seen in experiments, Figs. 7h,j^121,122^, Fig. 7h-j. Vector-HaSH predicts similar splitter-like and directionality-dependence in the spatial tuning curves of grid cells, SI Fig. S27.

Finally, we consider the dynamics of memory consolidation within the hippocampus. Hippocampal memories generally get transferred to cortex over time, but at least some episodic memories, for some duration, remain hippocampally-dependent. Hippocampal damage results in recall degradation, but items seen or recalled repeatedly are more resistant to damage^124,125^. To model these effects, we exposed Vector-HaSH to a set of inputs. A fraction were presented or recalled multiple times. Each presentation or recall led to a further increment of the hippocampal-sensory weights, or the hippocampal-sensory and hippocampal-to-grid weights, with the same learning rules as before. We found that memories reinforced in this way are remembered with richer detail relative to the rest, Fig.7l. After removal of a fraction of the hippocampal cells, recall of the consolidated items remains relatively robust, Fig. 7m, SI Fig. S19. There was no major difference when hippocampal-to-grid weights were additionally strengthened relative to the consolidative effect of sensory-hippocampal weight strengthening. Within the sensory-hippocampal pathway, the primary effect on consolidation was from strengthening of the hippocampal-to-sensory weights rather than the sensory-to-hippocampal weights, Fig.7n. These results are consistent with and provide a more specific prediction about mechanisms of memory consolidation, which Vector-HaSH suggests might be localized to directed synapses from hippocampus to cortex^126^.

An alternative hypothetical mechanism of consolidation is that a repeated sensory state activates new scaffold states, creating multiple associations between the sensory state and the scaffold states. Our numerical results in Vector-HaSH (1000 runs; data not shown) did not support this mechanism: associating an input with two different scaffold states regularly (100% of runs) resulted in the activation of a third, unrelated scaffold state when the partial sensory input was presented for recall.

### Mechanism for the method of loci (memory palace) technique

An intriguing memory technique known for millennia, the memory palace or method of loci^127–130^, is widely exploited by memory athletes in mnemonic competitions^131,132^. Given a list of typically non-spatial items to memorize, such as a sequence of playing cards, memory athletes imagine walking along a specific path through a familiar and richly remembered space, such as one’s childhood home or school, Fig. 8a. As they walk, they “place” or attach items from the sequence to be memorized at different locations along the walk. At recall time, they repeat the walk and “collect” the items they had placed. Counterintuitively, by adding to their memory task the additional demand of forming and recollecting new item-spatial associations, they are able to perform highly accurate one-shot memorization and recall^133,134^.

**Figure 8.**
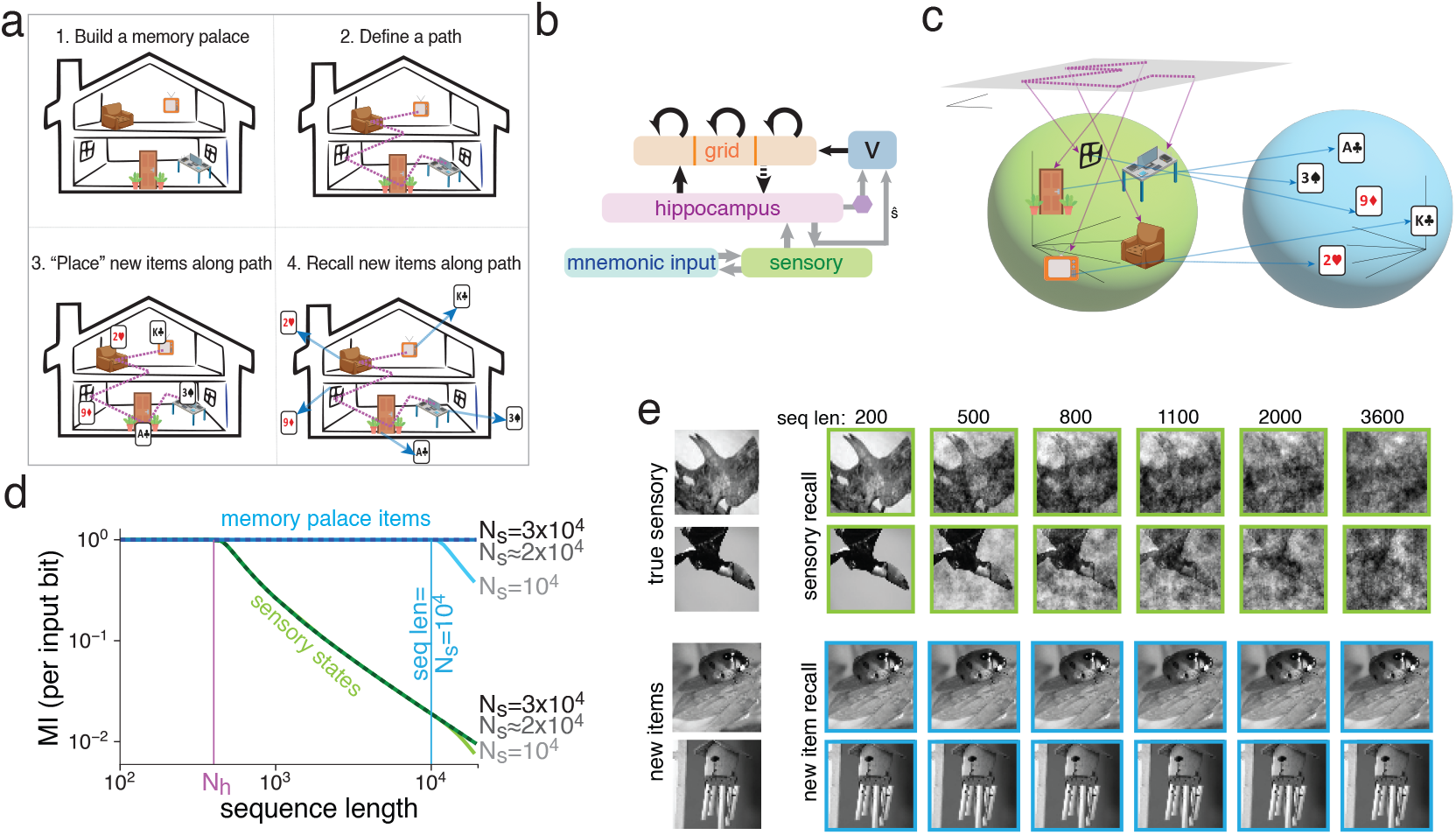
Method of loci: Accurate recall of arbitrary inputs by heteroassociation onto landmarks in a “memory palace” formed by Vector-HaSH. (a) The memory palace technique (figure adapted from^135^). (b) Model of the Method of loci: the full sequential Vector-HaSH circuit serves as a memory scaffold for new inputs, via heteroassociation with the sensory network. The path through the memory palace is learned via sequence learning, as in Fig. 6. The recollected sensory landmarks along the path are now (hetero)associated with the new items to be stored. (c) The large sensory area encoding spatial landmarks now forms the scaffold basis for heteroassociation of arbitrary sensory inputs in neocortex (cf. the smaller hippocampal basis for heteroassociation of sensory inputs within Vector-HaSH). (d-e) Scaffold states for neocortical input in the memory palace model are the (potentially imprecisely) recalled spatial landmarks. Because landmark recall is reliable even when imprecise (Fig. 3g, new items represented in neocortex can be recalled perfectly deep into the memory continuum even where landmark recall is substantially degraded relative to the true landmark, (d). Error bars (too small to be visible): standard deviation over 10 runs. (e) Two examples of the actual spatial landmarks, the recalled sensory states, and the recalled new neocortical inputs in the memory palace model, at different depths into the memory continuum. In (d-e), *λ* = {3, 4, 5} , *N*_*h*_ = 400.

A simple extension of Vector-HaSH provides the first model for how memory palaces might work, helping to explain their power. Deep in the memory continuum of Vector-HaSH, memorized items are only approximately reconstructed, thus the items to be memorized cannot be recalled well using the native memory mechanism. Instead, suppose Vector-HaSH is initialized to a starting location in a highly familiar space, and items at locations along the path taken by the memory athlete through this space are recalled. We assume that the neocortical representations of the sequence of new items to be remembered are now associated with the *recalled sensory states* along the spatial trajectory. Crucially, even though the recalled sensory states are mere approximations of the actual sensory inputs, the approximate states are reliably the same because sensory reconstruction in Vector-HaSH is reliable even when approximate, Fig. 3b. In other words, the recalled sensory states even if deep in the memory continuum can effectively be used as an extension of the fixed scaffold, now playing the scaffold role for new neocortical inputs, Fig. 8b-c. Association with this reconstructed state allows the new patterns to themselves be highly accurately reconstructed even in the memory continuum of Vector-HaSH.

Two advantages of using this extended scaffold are that the effective information capacity bottleneck, which was previously the size of the hippocampus, is now the size of the sensory input area and could be much larger, meaning that perfect memorization of a much larger number of inputs is possible, Fig. 8d; and that even it were the same size, if sensory reconstruction is deep in the continuum regime, the extended scaffold can be newly utilized and thus would be far from its continuum.

The result of heteroassociating new neocortical sequences with recalled sensory states from a previous familiar memory is that the recall of the new sequence is precise and detailed, Fig. 8e (bottom two rows), even when the hippocampal-entorhinal circuit is operating in the memory continuum and the sensory recalled states are only approximate, Fig. 8e (top two rows).

A prediction of Vector-HaSH during memory palace encoding and recall is the activation of the core grid cell and hippocampal representations underlying locations in the palace, not merely (and more faithfully than) the sensory states corresponding to those locations.

## Discussion

### Related models

Vector-HaSH adds to and helps define a new class of memory models^86,136^that we will name *robust hash-based memory*: All these models share the idea of a large pre-structured set of robust (large-basin) fixed point states that serve as abstract error-correcting indices for memory content. However, Ref. 136 does not model how to store externally specified inputs. Ref. 86 shares the tripartite architecture of Vector-HaSH, with heteroassociation of external inputs to a scaffold. However, Ref. 86 does not include grid cell-like representations; as a consequence, it lacks the properties of strong generalization and high-capacity sequence memory.

Dense associative memories are capable of storing large numbers of patterns, however their implementation is abstract in the form an gradient descent on energy landscapes; when implemented in a neural circuit with pairwise synapses, they require exponentially many neurons (rather than the linearly many of Vector-HaSH) to store exponentially many patterns^76,137,138^. An orthogonal class of memory models considers the problem that synaptic strength bounds lead to the collapse of performance in Hopfield-like networks. They show that adding multiple timescales per synapse can rescue performance towards unbounded Hopfield levels^139,140^. Our model is focused on improving over Hopfield-level performance, and the solution is architectural. Because the architectural level is complementary to the single-synapse level, it can incorporate such dynamics in the future.

Vector-HaSH is similar to the models of Refs. 141, 142, which also involve fixed grid cell representations interacting with hippocampal cells. In Ref. 141, hippocampal cells drive a low-dimensional update mechanism via grid cells, much like our vector updating model. However, in both these models the grid-hippocampal weights are learned and thus the scaffold is not invariant even though the grid representations are fixed. Therefore, they lack the properties of high capacity, large robust basins, avoidance of catastrophic forgetting and the memory cliff and strong generalization. Further, because the model in Ref. 142 does not store external inputs, it is not a memory model. Another related model of the hippocampal complex is that of Ref. 72. Like Refs. 141,142, it lacks a fixed scaffold, in two senses: there is no fixed grid representation (recurrent interactions within the proto-grid network are (re)learned for each environment), and the grid-hippocampal weights are learned. Thus it too lacks the high capacity, large basins, and avoidance of catastrophic forgetting properties made possible by a fixed scaffold. Memory formation occurs via gradual error backpropagration rather than rapid associative learning. It also lacks the low-dimensional shift mechanism learned from hippocampal or sensory states, and therefore the ability to store sequential memory at any level approaching that needed for episodic memory. Ref. 72 involves recurrent learning within hippocampus, whereas the current iteration of Vector-HaSH does not yet include recurrent excitation within hippocampus.

Overall, Vector-HaSH contrasts with those involving learning of the geometric and topological structure of the environment^143^, such as successor representation^144^, PCA^42^ and TEM^72^. Because all the internal representations in these models is based on environment dimension, topology, and geometry, they do not predict invariant internal grid representations or preserve cell-cell relationships across environments of distinct geometries. These models also require sensory inputs to form representations in new environments, in contrast with Vector-HaSH and animals, which form grid-like representations from the first pass in novel environments, even in the dark. Vector-HaSH does not contain mechanisms for grid cell deviations from hexagonal spatial tuning in deformed environments; exploring the addition of slow time-scale plasticity in the grid-hippocampal scaffold synapses, which should generate such effects while maintaining cell-cell relationships, is an interesting future research direction.

Given that the hippocampus is orders of magnitude smaller than the cortical states that represent events memorized by hippocampus and that memories involve combinations of different sensory inputs, it is clear that any model of hippocampally-based memory must involve state compression. In Vector-HaSH, the hippocampal representation is compact because the grid-hippocampal circuit functions as a content-independent *pointer* or hash mechanism for content in the cortex: the grid-driven hippocampal states are hashes of the sensory inputs. In its conceptualization as a pointer system, Vector-HaSH resembles the models of^145,146^. For episodic memory, the pointers are locality-sensitive hashes (LSH) (with locality determined in the temporal rather than content deomains) because contiguous grid states determine the scaffold states to which near-in-time inputs are attached, meaning that near-in-time inputs will have more-similar scaffold representations.

A contrasting way to compress information is via content-based compression, as done by the bottleneck layer of an autoencoder (Fig. 3 and Refs. 99, 147). As we have seen here, direct content compression through learning is not highly performant: these models lack the capacity, resistance to catastrophic forgetting, and sequence memory properties of Vector-HaSH.

Nevertheless, the commonalities among many of these models point toward a converging view of the hippocampal complex. The highly performant features of Vector-HaSH suggest a first-draft computational understanding of the circuit mechanisms of hippocampus as a highly performant memory system. Vector-HaSH is the first model we know of, besides MESH^148^, that is capable of storing an exponentially large set of input patterns in an associative content-addressable memory.

### Future extensions and directions

Vector-HaSH accounts for a range of phenomena in entorhinal cortex and hippocampus^101,119–122,124,125^. At the same time, there are numerous avenues for future research and extensions. These include incorporating different subregions of the hippocampus; mechanistically relating the ordered and phasic updates that are assumed in Vector-HaSH (from sensory to hippocampal to grid, back to hippocampus, then out to sensory) to hypothesized gating roles for the hippocampal LFP oscillations^23,149–154^; investigating how the hippocampus and Vector-HaSH deal with conflicts between internal states and external cues, and with changes in primary versus contextual inputs; modeling and experimentally determining how current and target states might be simultaneously represented within the same circuit; examining how hierarchical and similarity-respecting representations for distinct but similar memories are encoded; exploring the dynamics and circuit mechanisms of fragmentation of space and events into submaps and discrete episodes (e.g. via surprisal^103^); exploring the roles of within-hippocampus recurrence and different hippocampal subregions; showing how slower time-scale plascticity in the sensory-hippocampal weights could result in map distortions and representational drift^155^; characterizing the contribution of different specific cell types and their roles in episodic memory within this circuit; and modeling the process of memory transfer from hippocampus to cortex.

### Relationship to anatomy

Vector-HaSH is based on the structure of hippocampal-entorhinal circuitry. However, in some respects it varies from the classic view: While hippocampal outputs to and inputs from entorhinal cortex are believed to be separated between deep versus superficial layers of entorhinal cortex, the scaffold involves a tight loop in which grid cells drive hippocampus and receive direct mirrored input back in a way that reinforces the input grid patterns. This structure is a prediction that the deep-to-superficial entorhinal projection closes a fully self-consistent loop, which can potentially be tested connectomically. Although much is known about entorhinal-hippocampal circuitry, new discoveries can still surprise: recent reports show that deep layers of EC, which receive hippocampal inputs and were believed to primarily send outputs to neocortex in fact send a copy of their outputs back to the hippocampus^156^.

Random and fixed grid-to-hippocampal weights were key for several properties of Vector-HaSH. Intriguingly, developmental work from^157^ shows that the stellate layer II MEC neuron circuit (which form the majority of the grid cell population) was the first to mature, and activity in this network then drove maturation of the hippocampal circuit, followed by layer 5 of entorhinal cortex, and finally, layer II of LEC. This maturation order is very closely tied to structure formation in our model: we posit that the scaffold is formed first, and within it, the grid cell circuit is formed first so it can build the fixed connections to hippocampus.

Are random connections a unique solution for Vector-HaSH? Our theory and simulations show the sufficiency of random weights and the insufficiency of several types of non-random or learned weights. However, they do not eliminate the possibility of non-random solutions. The situation is similar to the construction of expander graphs for high-capacity error-correcting codes^158^: though non-random solutions can exist in principle, it has been hard to find them, while random connections are sufficient. Because Vector-HaSH is a full dynamical neural circuit, it can be easily and directly queried for experimental predictions about representation, dynamics, and learning under a large variety of conditions and perturbations. There might be connectomic or ultrastructural signatures of fixed versus plastic synapses and their development order.

### Summary

In sum, we have proposed a model that unifies the spatial and episodic memory roles of the hippocampal complex by showing that nominally spatial representations and architectures are critical for a well-behaved episodic memory, even if the memories are devoid of spatial content. Unlike unstructured recurrent memory models^76,84,93–99,137,138,159^, Vector-HaSH involves two key novel features. The first is the *factorization* of the associative memory problem into the creation of an abstract rigid fixed point scaffold (for robust denoising and autoassociative recall of abstract states and state sequences), and separately a feedforward heteroassociation (“pointer” creation) step in which content is attached to these abstract states. The second is the use of low-dimensional structured states (via grid cells) with random projections to a larger area to generate the scaffold and ensure large basins. For sequence memory, the Euclidean metric nature of the grid states became critical, so that mere two-dimensional vectors rather than full information about the next-state pattern were required to recurrently resurrect the next state.

The prestructured scaffold concept agrees with results and hypotheses about the existence of internally generated underlying states and sequences that become associated with new experiences and inputs^66,113,160–162^, and with the observed re-use of the same circuit responses across spatial and non-spatial domains^67,163–165^including most recently even in mental (imagined) traversals in non-spatial domains^73^; recent findings in cortex, in which sensory responses are shaped by pre-existing internal states^166^, suggest that the brain might more broadly exploit prestructured scaffolds in its representations.

Vector-HaSH provides a computational hypothesis for the mechanisms of the memory palace technique, based on understanding the advantages of co-localizing spatial and non-spatial memory. The model explains why impressive memory performance does not require exceptional intellectual ability or structural brain differences, but can be leveraged by anybody trained to appropriately engage the hippocampus^131,133,167^. From a neuro-AI perspective, the specific biological architectures, representations, and learning rules of Vector-HaSH led it to significantly outperform fully end-to-end supervised trainable memory models with similar architectures, comparable or more parameters, and fewer constraints – a clear realization of the hypothesis that biological structures (inductive biases) can produce better performance than fully end-to-end trainable models as commonly used in machine learning.

## Methods

### Method Details

In Ref. 86, the MESH associative memory architecture was introduced, leveraging a three-layer network to store numerous independent memory states. This architecture allowed for a high-capacity memory with a trade-off between the number of stored patterns and the fidelity of their recall. However, MESH did not require specifically grid cell encodings, did not exhibit strong generalization in scaffold learning, and did not exhibit a high sequence capacity.

In Vector-HaSH, our memory scaffold consists of a recurrent circuit incorporating MEC grid cells and a hippocampal layer that may be interpreted as the proximal CA1 and distal CA3 regions of the hippocampal complex.

Specifically, we represent the MEC grid cells as outlined in Ref.^115^, where each grid module’s state is expressed using a one-hot encoded vector that represents the module’s phase (and thus the active grid cell group within the module). The states are on a two-dimensional discretized hexagonal lattice with period *λ* . Thus, the state of each grid module is represented by a vector with a dimensionality of *λ* ^2^.

*M* such grid modules are concatenated together to form a collective grid state 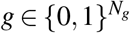, where the 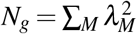. The continuous attractor recurrence in the grid layer^33^ is represented by a module-wise winner-take-all dynamics, which ensures that the equilibrium states of *g* always correspond to a valid grid-coding state.

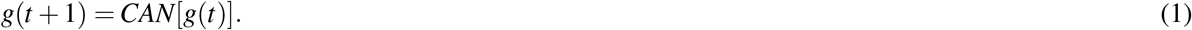

We represent these equilibrium states by 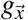, where we index the coding states by the two-dimensional location 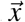. For coprime periods *λ*_*M*_, the grid states can encode a spatial extent of 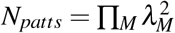 spatial locations.

This layer of grid cells projects randomly onto the hippocampal layer, through a *N*_*h*_ *× N*_*g*_ random matrix *W*_*hg*_, with each element drawn independently from a Gaussian distribution with a mean of zero and standard-deviation one *N*(0, 1). This matrix is sparsified such that only a *γ* fraction of connections is retained, leading to a sparse random projection. This projection constructs an *N*_*h*_ dimensional set of hippocampal sparse states, 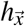 defined as

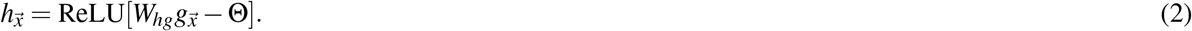

The return weights from the hippocampal layer back to the grid cell layer is set up through Hebbian learning between the predetermined set of grid and hippocampal states, _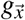_ and _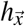_.

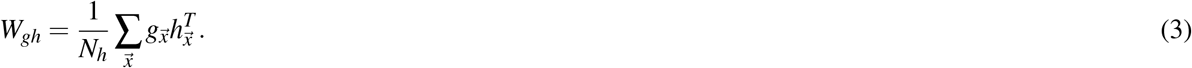

The dynamics of the hippocampal scaffold is then set up as

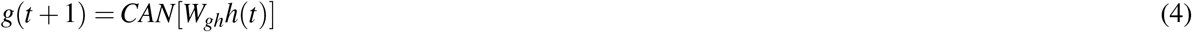

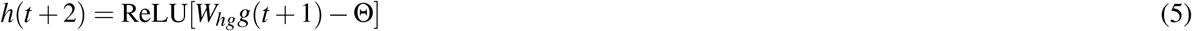

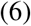

These equations maintain each 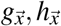 state as a fixed point of the recurrent dynamics, as we prove in SI Sec. C.1.

This constructed hippocampal memory scaffold is then used to generate independent memory locations to store information presented through a sensory encoding layer, representing the non-grid cell component of the Entorhinal cortex. Information to be stored is presented as a binary encoding of states in the sensory layer, and is ‘tagged’ onto a memory location 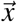 of the scaffold through pseudo-inverse learned heteroassociative weights.

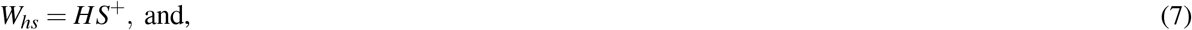

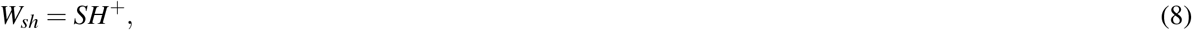

where *H* is a *N*_*h*_ *× N*_*patts*_ dimensional matrix with columns as the predetermined hippocampal states _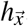_, and *S* is a *N*_*s*_ *N*_*patts*_ dimensional matrix with columns as the encoded sensory inputs to be stored at location 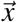. To reduce computational time-complexity, we use an exact pseudoinverse rather than an iterative pseudoinverse for calculation of these inter-layer weights, unless otherwise specified.

Given the above equations, we can now perform bi-directional inference of sensory inputs from grid states and vice versa:

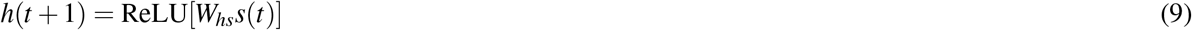

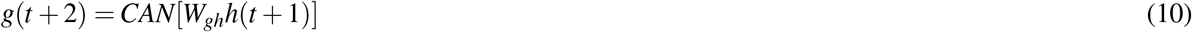

and

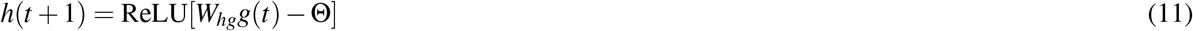

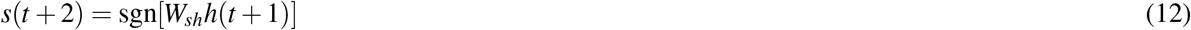

The above two sets of equations can then be combined to use Vector-HaSH as a content-addressable memory to recover stored sensory inputs from corrupted inputs — first the grid states are inferred from the corrupted sensory input, and then the true sensory input is recalled from the inferred grid state.

The above equations have been written considering sensory inputs to be random binary states. In cases where sensory states are continuous valued (as in Fig. 3b, for example) the *s* reconstruction equation, Eq. (12) is replaced with simply *s*(*t* + 2) = *W*_*sh*_*h*(*t* + 2).

Equations (1-12) describe the core working of Vector-HaSH— this core version and its variants can then be used to generate item memory, spatial memory, episodic memory, as well as a wide range of experimental observations, such as those discussed in Fig. 7. In general, across all models, we assume that the relevant synapses are plastic during input presentation for memory storage, and are frozen during testing of memory retrieval.

#### 1 High-capacity pattern reconstruction

For the basic task of pattern storage and reconstruction, we utilize the simplest form of Vector-HaSH without any additional components. To examine reconstruction capacity, *N*_*patts*_ sensory cues are stored in the network via training the *W*_*hs*_ and *W*_*sh*_ weights as described in Eqs. (7-8).

The *N*_*patts*_ sensory cues need to be stored corresponding to distinct scaffold states. In our implementation, for simplicity, we selected scaffold states in a “hairpin” like traversal, similar to that shown in Fig. 5a *top middle* to achieve this.

Then, a clean or corrupted version of a previously stored pattern is presented to the network in the sensory encoding layer, which then propagates through the network via Eqs. (9-12), finally generating the recalled pattern *s*. In all numerical examples we consider in the main text we either construct random binary {− 1, 1} patterns, or consider images from `mini-imagenet`^168^. In particular, we took 3600 images from the first 6 classes {‘house-finch’, ‘robin’, ‘triceratops’, ‘green-mamba’, ‘harvestman’, ‘toucan’} and center-cropped them to consider the middle 60 × 60 image and converted them to grayscale. We refer to this set of grayscale images as `bw-mini-imagenet`. In all models, the memorized patterns are a noise-free set, then we test memory recall with noise-free, partial, or noisy cues.

In Figs. 2, 3, the recall performance and quality was examined in networks with three grid modules, *γ* = 0.6, and *θ* = 0.5.

The capacity in Fig. 2c(*right*) was evaluated by injecting a noise into the hippocampal layer of magnitude 20% of the magnitude of the hippocampal state vector, and requiring the iterated dynamics to return the hippocampal state to within 0.6% of the original hippocampal state (Here magnitudes and distances were calculated via an *L*^2^ metric).

In Fig. 2d(*left*) and SI Fig. S2, the critical 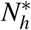 is estimated as the smallest value of *N*_*h*_ such that all scaffold states have been stabilized as fixed points. The corresponding module periods for data points plotted in SI Fig. S2, for two modules are listed in Table. 1 left, and for three modules are listed in Table. 1 right. Similarly, the grid module periods for the data in Fig. 2c(*left*) are listed in Table 2.

**Table 1.** Grid module periods, number of grid cells and total number of patterns for data in Fig. 2e.

| $\lambda$ | $N_g$ | $N_{patts}$ |
| --- | --- | --- |
| 2,3 | 13 | 36 |
| 3,4 | 25 | 144 |
| 4,5 | 41 | 400 |
| 5,6 | 61 | 900 |
| 6,7 | 85 | 1764 |
| 7,8 | 113 | 3136 |
| 1,2,3 | 14 | 36 |
| 2,3,5 | 38 | 900 |
| 3,4,5 | 50 | 3600 |
| 4,5,7 | 90 | 19,600 |
| 5,6,7 | 110 | 44,100 |

**Table 2.** Grid module periods, number of grid cells and total number of patterns for data in Fig. 2f.

| $\lambda$ | $N_g$ | $N_{patts}$ |
| --- | --- | --- |
| 7,8 | 113 | 3136 |
| 3,5,8 | 98 | 14400 |
| 3,4,5,7 | 99 | 176400 |
| 1,3,4,5,7 | 100 | 176400 |

To estimate the basin sizes of the patterns stored in the scaffold, as shown in Fig. 2e, we compute the probability that a given pattern is perfectly recovered (i.e., remains within its correct basin) as we perturb the hippocampal states with a vector of increasing magnitude. We assume that the size of any given basin can be estimated as the typical magnitude of perturbation that keeps the system within the same basin of attraction — this is not generally true for non-convex basins, particularly in high-dimensional spaces. However, this estimate is relevant in the context of testing robustness under corruption with uncorrelated noise. Further, we later demonstrate in SI Sec. C.3 that the basins are indeed convex. Here grid module periods *λ* = {3, 4, 5} , number of grid cells *N*_*g*_ = 50, and *N*_*h*_ = 400 hippocampal cells were used. Probability that a given pattern remains within its correct basin was estimated by computing the fraction of runs where a given pattern was correctly recovered for a 100 different random realization of the injected noise.

Figure 2f examines the learning generalization in Vector-HaSH, i.e., the capability of Vector-HaSH to self-generate fixed points corresponding to scaffold grid-hippocampal states despite training on a smaller number of fixed points. For a given number of training patterns, we calculate the number of generated fixed points by counting the number of states that when initialized at a scaffold state remain fixed upon iteration through Eqs. 4,5. As discussed in the main text, when training on a given number of training patterns (that is less than the complete set of all patterns), the ordering of the patterns is crucial in controlling the generalization properties of the model. For Vector-HaSH, we order patterns such that a two-dimensional contiguous region of space is covered (see Sec. C.4 for additional details of the ordering and the freedom of possibilities in this ordering), resulting in the strongest generalization (Sec. C.4). For comparison, in Fig. 2f we also consider “shuffled hippocampal states”, wherein scaffold states are randomized in order before subsets are selected for training. We also consider “random hippocampal states”: here we consider each hippocampal state vector and randomize its indices, in effect constructing a new state vector with exactly the same sparsity and statistics, but now uncorrelated to the grid state corresponding to that hippocampal state. Then, we use bi-directional pseudoinverse learning between grid and hippocampal states and construct this as a scaffold. This lack of structured correlations between grid and hippocampal population vectors results in catastrophic forgetting, with no observed fixed points remaining once all scaffold states have been used for training.

All curves shown in Fig. 3c-f are averaged over 5 runs with different random initialization of the predefined sparse connectivity matrix *W*_*hg*_, error bars shown as shaded regions represent standard deviation across runs. In Figs. 3b,e,h, grid module periods *λ* = {3, 4, 5} , *N*_*g*_ = 50, *N*_*s*_ = 3600 was used. The total capacity of the network in this case is capped by 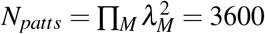. In Fig. 3d, all shown networks have ≈5 × 10^5^ synapses, with Vector-HaSH module periods *λ* = {2, 3, 5} , and layer sizes: *N*_*g*_ = 38, *N*_*h*_ = 275, *N*_*s*_ = 900. Number of nodes in other networks are as follows: (i) Hopfield network of size *N* = 708, synapses = *N*^2^. (ii) Pseudoinverse Hopfield network of size *N* = 708, synapses = *N*^2^. (iii) Hopfield network with bounded synapses was trained with Hebbian learning on sequentially seen patterns. Size of the network *N* = 708, synapses = *N*^2^. (iv) Sparse Hopfield network (with sparse inputs) with a network size of *N* = 708, synapses = *N*^2^, sparsity = 100(1 − *p*). (v) Sparse Hopfield network. Size of the network *N*, synapse dilution *κ*, synapses = *κ × N*^2^ = 10^5^. (vi) Tailbiting Overparameterized Autoencoder^99^ with network layer sizes 900, 275, 38, 275, 900.

For stored patterns of size *N*, recall of an independent random vector of size *N* would appear to have a mutual information of 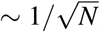, which when evaluating the total mutual information across all 𝒪 (*N*) patterns or more would appear to scale as 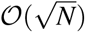, despite no actual information being recalled. To prevent this apparent information recall, in Fig. 3f if the information recall is smaller than 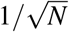 we then set it explicitly to zero.

To examine Vector-HaSH’s performance on patterns with correlations, in Fig. 3e we trained it on `bw-mini-imagenet` images using grid module sizes *λ* = {3, 4, 5} , and layer sizes: *N*_*g*_ = 50, *N*_*h*_ = 400, *N*_*s*_ = 3600. The plotted curve shows the mean-subtracted cosine similarity between recovered and stored patterns illustrating that Vector-HaSH shows gradual degradation as the number of stored patterns is increased. The resultant curve is an average over 5 runs with different sparse random projections *W*_*hg*_.

##### 1.1 Multiple Traces Theory

In Figs. 7l-n, we consider Vector-HaSH with *λ* = {3, 4, 5} , *N*_*g*_ = 50, *N*_*h*_ = 400, *N*_*s*_ = 3600, *γ* = 0.6, and *θ* = 0.5. We use random binary patterns in n, and `bw-mini-imagenet` patterns in l-m. The results are averaged over 20 runs. For sensory inputs presented multiple times, the sensory hippocampal weights are reinforced multiple times using online pseudoinverse learning rule^90^, and the grid hippocampal weights are reinforced multiple times using Hebbian learning (Fig 7b, right). The *W*_*hs*_ weights are invariant to reinforcement due to the iterative pseudoinverse causing perfect hippocampal reconstruction from sensory inputs. Given a particular lesion size, the cells to be lesioned are randomly chosen from the set of all hippocampal cells, and their activation is set to zero. Sensory recovery error is defined as the mean L2-norm between the ground truth image and the image reconstructed by the model. During testing, the model receives the ground truth sensory image as input, and the reconstruction dynamics follow Eqs. (9-12). Additional results from each layer of Vector-HaSH while testing the Multiple-Trace Theory are shown in Fig. S19, right. Further, Fig. S19, left shows the results when only *W*_*sh*_ weights are reinforced, assuming pre-trained scaffold weights *W*_*gh*_. In both case, same parameter settings were used as in Figs. 7l-n.

#### 2 Mapping, recall, and zero-shot inference in multiple spatial environments without catastrophic interference

Here we add a path-integration component to Vector-HaSH, that utilizes a velocity input to change the grid cell population activity akin to Ref.^33^, such that the phase represented by each module changes in correspondence to the velocity input. Corresponding to the discrete hexagonal lattice space used to represent each grid module, for simplicity the velocity is assumed to have one of six directions, and magnitude is assumed to be fixed at a constant such that the phase of each grid module updates by a single lattice point in a single time-step. This input velocity vector, that we call a velocity shift operator,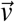, is thus represented by a six-dimensional one-hot encoded vector determining the direction of the shift.

In order to capture the inherent randomness and uncertainty present in real-world scenarios, a small amount of neuronal noise was introduced by adding random perturbations to the activation values of hippocampal cells in Vector-HaSH. This noise, generated from a uniform distribution between 0 and 0.1, mimics the fluctuations and disturbances observed in individual neurons, and corresponds to a noise magnitude of roughly 25% the magnitude of the hippocampal state vectors.

In Fig. 4a,c we first demonstrate bi-directional recall of grid states from sensory inputs and vice versa. Here we consider Vector-HaSH with *λ* = {3, 4, 5} , *N*_*g*_ = 50, *N*_*h*_ = 400, *N*_*s*_ = 3600. We train the model on a total of 600 sensory inputs taken from `bw-mini-imagenet` (including the 4 landmarks placed in the room shown in Fig. 4c). To demonstrate zero-shot recall in panel c, the model dynamics are simulated on a novel trajectory (right) through the same room with some locations overlapping with the previous trajectory. Note that the reconstructed landmarks do not have perfect recall. Instead, the reconstructions are degraded relative to the originally stored landmarks since the total number of stored landmarks in the model exceeds *N*_*h*_ = 400 (Fig. 2f).

For all other panels of Figure 4, we use Vector-HaSH with grid module periods *λ* = {3, 4, 5, 7} , *N*_*g*_ = 99, *N*_*h*_ = 342, *γ* = 0.1, and *θ* = 2.5. The total capacity of this grid coding space is 176400 ≈2 × 10^5^. Each room is stored by allocating a random 10 × 10 patch of the grid coding space to it (This is constructed by first choosing any random point in the room to map to a randomly chosen area of the grid coding space. Then as the model moves in the room, path integration correspondingly updates the grid phases in each grid module. The region of grid coding space explored as the model physically explores a room is then the patch of grid coding space storing the particular room).

To each of the 100 locations comprising a room, we simulate an independent sensory landmark as a binary {−1, 1} vectors. At initialization, before observing any room, we begin with a pretrained memory scaffold, wherein the *W*_*hg*_ and *W*_*gh*_ matrices have already been constructed and trained corresponding to Eqs. 2, 3.

When first brought to a room, the grid state is initialized to the grid state vector corresponding to the random region of grid coding space allocated to the room. Then, as path integration updates the grid state upon moving around the room, the observed sensory landmark states are associated with the corresponding grid-hippocampal scaffold states through learning the *W*_*hs*_ and *W*_*sh*_ matrices following Eqs. 7, 8.

In the first two tests of each room (first tested right after each room has been learned, and then tested after all rooms have been learned; shown in Fig. 4d) sensory landmark cues can be observed by Vector-HaSH. Using Eq. 9, the observed sensory landmarks can be used to reconstruct the hippocampal state, resulting in the reliably reconstructed hippocampal tuning curves as seen in Fig. 4e. For testing stable recall in dark (Fig. 4d,e), Vector-HaSH is provided a random single sensory landmark cue from any given room. This landmark is used to ascertain the grid state corresponding to that landmark through Eq. 9. Thereafter, path integration is used to construct the grid-hippocampal scaffold state as room is explored in the absence of any further sensory cues. As seen in Fig. 4e this also reliably reconstructs the hippocampal state at each location in every room.

In Fig. 4f, we examine the dark recall of 3600-dimensional sensory landmarks in each room in a continual learning setting. Here we begin again with simply the pretrained grid-hippocampal scaffold. As the *i*^th^ room is explored, the sensory-hippocampal weight matrices are updated to store the thus far observed landmarks and their locations. At each step of exploration within the *i*^th^ room, Vector-HaSH is queried on the current and all previous rooms in the following fashion: for any completed room *j* (i.e., 0 ≤ *j* < *i*), Vector-HaSH is dropped randomly anywhere in the room and allowed to observe the sensory landmark solely at that start location and no further sensory landmarks. Then the model moves around the room through path integration, and attempts to predict the sensory landmarks that would be observed at each location. We then compute the average mutual information recovered for each landmark at each position in the room, which is shown in Fig. 4f. For the partially completed room *i*, Vector-HaSH is similarly dropped randomly in the room, restricted to the set of previously observed locations within the room. The mutual information recovered during sensory prediction is similarly only evaluated over the previously observed portion of the room.

For the baseline model shown in Fig. 4f, we first construct the grid-hippocampal network through random hippocampal states with the same sparsity as those in Vector-HaSH, and bi-directional pseudoinverse learning between grid and hippocampal layers. Thereafter, the sensory landmarks are associated with the hippocampal layer as in Vector-HaSH described above, and this baseline model is subjected to an identical test protocol to examine continual learning. The number of nodes in the baseline model is kept identical to Vector-HaSH.

For Fig. 4h, we follow the same analysis as in the experiment^101^. Dot product between population vectors (PVs) across all combinations of the eleven test rooms were computed. To construct the population vectors, we record the activations of hippocampal cells for each of the 10 × 10 positions in the simulated room. We stack these into 100 composite population vectors (PVs), one for each position in the room. To compute overlaps between representations, the activation of each hippocampal cell in any particular room was expressed as a ratio of its activation to the maximal activation of that cell across all rooms. The overlap was then calculated as the normalized dot product between the hippocampal cell activation vectors in two rooms i.e., the sum of the products of corresponding components divided by the total number of hippocampal cells (*N*_*h*_ = 342) for a given position/pixel, averaged over 100 positions. The color-coded matrix in Fig. 4h shows the average dot product values for PVs across rooms ( ^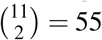^ room pairs). Repeated exposures to three familiar rooms were also added to this analysis leading to a total of 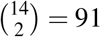 room pairs.

For Fig. 4j, we plot the distribution of PV normalized dot products computed above (for multiple visits to all the rooms) and use this PDF to compute the corresponding CDF. Similarly, the CDFs for shuffled data are computed through the same procedure, but using shuffled data to compute the PV normalized dot products. Shuffled data is obtained either by random assignment of rate maps across rooms (shuffle room) or by shuffling of cell identities within rooms (shuffle cells) or by a combination of the two procedures (shuffle room and cells). The number of different shuffles generated in each case was 1000.

#### 3 Extension of Vector-HaSH to continuous space

So far we have considered the grid states to be {0,1}-valued discretely varying modular one-hot states. This leads to a finite number of grid phases per module, and hence a finite number of grid population vectors that can be exactly enumerated, leading to the wealth of theoretical advancements and results described above. To bring Vector-HaSH closer to biological realism, we constructed continuous-valued grid states, Fig. 4l, as a Gaussian bump of activity on a two-dimensional lattice of neurons with periodic boundary conditions, similar to the one-hot states on a periodic lattice considered earlier (cf. Fig. 2b). We set up the recurrent dynamics on this lattice as a simplified abstraction of a continuous attractor model (see Methods for details) to stabilize these grid states and enable path integration on the grid manifold. Since the number of phases in each module is now infinite (the Gaussian bump need not be centered on a neuron in the lattice) it is computationally challenging to demonstrate memory capacity results similar to our analysis for the discrete model above. As a proof of concept, we demonstrated landmark reconstruction from grid phases and vice-versa in Fig. 4m.

For Fig. 4m, we used 3 grid modules, consisting of 81, 144 and 225 cells each. The gaussian bump of activity in each module was constructed to have a standard deviation of 0.5. Continuous attractor dynamics we approximated through a circular mean to determine activity mean location, and re-initialization of a gaussian bump centered at the calculated mean location.

We also used this continuous extension of Vector-HaSH in Fig. 7e-f. The results presented in Fig. 7f are computed similarly to^116^. We first compute the temporal correlation between every pair of grid cells before and after hippocampal activation. We compute the correlation between these temporal correlations (shown by the vertical red dashed line), and compare this to 1000 random shuffles of the grid correlations (shown as the null distribution in black).

#### 4 Path learning in the hippocampal scaffold

Here again, we add a path-integration component to Vector-HaSH as described in the section above, such that a velocity shift operator,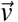, can be used to path integrate and update the grid cell population activity akin to Ref.^33^, such that the phase represented by each module changes in correspondence to the input shift.

For learning of trajectories in space, this vector 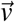 is either associated with spatial locations and corresponding hippocampal state vectors (as in path learning) or with sensory landmark inputs (as in route learning).

The results in Fig. 5 and Fig. 6 were generated using *N*_*h*_ = 500, *γ* = 0.6, *θ* = 0.5, *M* = 3 and 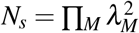 with *λ* = {5, 9, 13} in Fig. 5b-e, Fig. 6b and *λ* = {3, 4, 5} in Fig. 5f,g, Fig. 6c.

All networks in Fig. 6e were constructed to have approximately 5 × 10^5^ synapses, with network parameters identical to those in Fig. 3d. All panels in Fig. 6 considers random binary patterns, apart from Fig. 6c,d which considers `bw-mini-imagenet` images.

##### 4.1 Path learning

Learning associations from the hippocampal layer directly to the velocity inputs through pseudoinverse learning would result in perfect recall for only *N*_*seq*_ ≤ *N*_*h*_, which may be much smaller than the grid coding space, and would hence result in an incapability to recall very long sequences. To obtain higher capacity, we learn a map from the hippocampal cell state to the corresponding velocity inputs at that spatial location through a multi-layer perceptron, MLP. For all the results shown in Fig. 5 and Fig. 6c,h, we use a single hidden layer in the MLP with 250 nodes. The dynamics of the network are as follows:

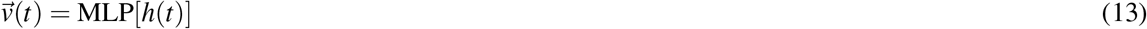

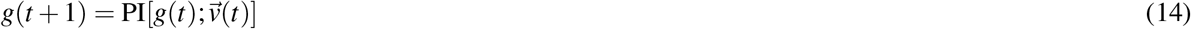

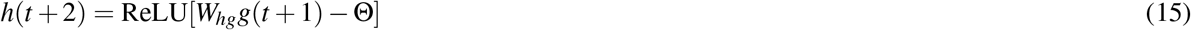

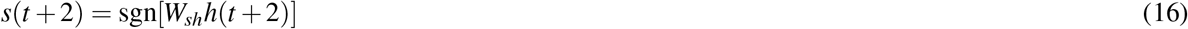

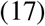

Thus, when cued with a sensory state at the start of an episode, the sensory inputs to hippocampus reconstruct the corresponding hippocampal and grid states. Then, through the MLP, the hippocampal state projects to a low-dimensional velocity vector that is used to update the grid cells via path integration. From this updated grid state, the corresponding hippocampal state is constructed, which then reconstructs the next sensory pattern of the episode. The new hippocampal state also maps to the next velocity vector, that continues the iteration by updating the grid state. In this fashion, the memory scaffold along with the MLP successively construct grid and hippocampal states, and the heteroassociative weights to the sensory layer successively construct the memorized patterns of the episode.

##### 4.2 Route learning

Since detailed sensory information cannot be recalled at very high capacities, route learning is performed by learning associations between the *recollection* of the sensory inputs at a location 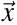, and the velocity shift vector 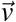 determining the direction of motion of the trajectory being learned at that location. This association can be learned directly through pseudoinverse learning as

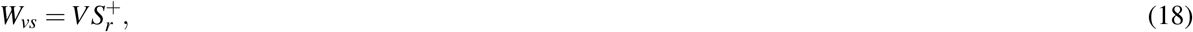

where, *S*_*r*_ is a *N*_*s*_ *× N*_*seq*_ dimensional matrix with columns as the recalled sensory inputs _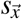_, and *V* is a 6 × *N*_*seq*_ dimensional matrix with columns as the corresponding velocities. These associations can then be used to recall long trajectories through

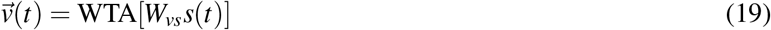

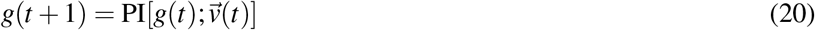

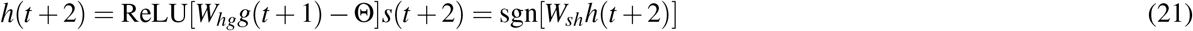

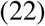

As argued in Sec. D.7, this results in perfect sequence recall for *N*_*seq*_ *≤ N*_*s*_, which can scale as the exponentially large capacity of the grid coding space. Note that the results in Sec. D.7 rely on *S*_*r*_ being a rank-ordered matrix. While this holds for random binary patterns through Eq. 21 applying a sign nonlinearity, this does not directly hold for continuous valued sensory states, where no nonlinearity is necessary. In this case, we take the input sensory patterns, and apply an inverse sigmoid function to them before storage in the *W*_*sh*_ matrix. Then, we use Eq. 21 with the sign nonlinearity replaced with the sigmoid nonlinearity. This application of the inverse sigmoid and then the sigmoid ensures that the final recovered states correspond to the inital patterns, but the sensory states are recovered through a nonlinear readout.

##### 4.3 Goal and context-based remapping

When initialized in a new environment, we model the grid state population activity to be randomly initialized in the grid-coding space (a mechanistic model for such random initialization will be discussed in future work), i.e., the grid state undergoes *remapping*. This grid coding state, along with the corresponding hippocampal coding state and sensory observations at that location are then stored in the corresponding weight matrices, i.e., *W*_*hs*_ and *W*_*sh*_, via Eqs. (7-8). When brought back to a previously seen environment, these weight matrices in Vector-HaSH use the observed sensory observations to drive the hippocampal cell (and hence grid cell) population activity to the state initialized at the first traversal of that environment.

Similar to new environments, we also model contextual information (such as goals, rewards, start-end location pairs) to be appended to the sensory inputs. We allow new contextual information to also trigger reinitialization of grid state, which then permits storage of multiple paths that involve the same spatial location, provided that they are distinguished by a contextual signal.

We use this set up of manual reinitialization of the grid state to reproduce the experimental observations of splitter cells^119^, route dependent place cells^120^, directional place fields in one-dimensional environments^121^ and on directed routes in two-dimensional environments^122^ in Figs. 7g-j and S22; and of directional place fields in a radial eight-arm maze^122^ in Fig. S24. In all of these cases, we first generate trajectories corresponding to the paths that the animals are constrained to traverse in the given experiment. These trajectories, are then stored in Vector-HaSH at a random location in the grid coding space through a path learning mechanism. At new contextual cues, the grid state in the model is reinitialized and the agent then continues at a new location in the grid coding space. This results in different spatial firing fields, irrespective of whether the agent is at the same spatial location as in a different previous context.

For all the simulations in Figs. 7g-j, S22 and Fig. S24, Vector-HaSH with *λ* = {3, 4, 5, 7} , *N*_*h*_ = 500, *N*_*g*_ = 99, *θ* = 2.0 and *γ* = 0.10 was used. The total size of the grid coding space is 420 × 420 ≈10^5^. In order to capture the inherent randomness and uncertainty present in real-world scenarios, a small amount of neuronal noise was introduced by adding random perturbations to the activation values of hippocampal cells in Vector-HaSH. This noise, generated from a uniform distribution between 0 and 0.1, mimics the fluctuations and disturbances observed in individual neurons.

###### Splitter cells

For Fig. 7h, we follow an analysis method similar to the analysis done on the experimental data^119^. The central stem is divided into 4 equal regions (Fig. S25b), and the mean activation of every hippocampal cell is computed in each of the four regions. Figure S25c plots mean activations in each of the four regions, of cells that show different activity patterns as Vector-HaSH traverses the central stem on Left-Turn and Right-Turn trials. The “activation ratio” on Right-Turn trials versus Left-Turn trials is then calculated for each cell in the region for which the given cell has maximum difference in activations. The distribution of these activation ratios is plotted in Fig. 7h, that shows the frequency distribution of cells with preferential firing associated with Left-Turn or Right-Turn trials. Note that the distribution of cells preferring left-turn and right-turn trials is approximately even. The percentage of hippocampal cells with non-differential firing was found to be ≈ 3.896%, and the percentage of hippocampal cells with differential firing was found to be ≈ 96.103% in Vector-HaSH (using a threshold of 2 on the activation ratio).

###### Route encoding

In Fig. S22a,c we employed an ensemble analysis approach mirroring that used in^120^ to validate if hippocampal cells demonstrate route-dependent activity. Our simulated session comprised four blocks, each representing one of four routes (0-3), with 11 trials per block. We performed ensemble analysis on the maze region common to all routes.

We compared the population vector (PV)—activations of all hippocampal cells on an individual trajectory—to the average activation of these cells across all trajectories on each route (route-PV). Specifically, we compared the PVs for each trajectory to the average activation population vectors (route-PVs) of all four routes, excluding the trajectory in consideration from its route-PV calculation to avoid bias.

Using cosine similarity, we assessed the likeness between each trajectory PV and each of the four route PVs. We then calculated the fraction of correct matches (the highest similarity score was with its corresponding route-PV) and incorrect matches (a higher similarity score was with a different route-PV). The comparison results are shown in Fig. S23a, left.

We repeated the process 10,000 times with randomized data to estimate the chance probability of correct matches. We randomized the session data by shuffling trials across blocks, randomly assigning each trajectory to one of the four routes, thereby disrupting any correlation between the hippocampal cell activations and a specific route. Fig. S23a,right depicts a typical result from one such shuffle.

For each matrix element (*i, j*), we plotted the distribution of data from these 10,000 matrices in Fig. S23b. We then estimated the Probability Density Function (PDF) from this distribution using a Gaussian kernel (Python’s `scipy.stats.gaussian_kde` method). To gauge the chance probability of correct matches in our original, unshuffled analysis, we calculated the percentile position of our observed match proportion, referencing the same matrix element (*i, j*) from the unshuffled matrix in Fig. S23a.

Fig. S22c presents the probability of correct matches in the unshuffled analysis based on these distributions from 10,000 shuffles. Low diagonal values indicate that trajectories significantly match only their corresponding route-PVs.

###### Directional cells

For Figs. 7j, S22d and Fig. S24, the directionality index is defined similar to that defined for the experimental data analysis^121^,^120^. Given the activation (*A*) of a hippocampal cell in positive and negative running directions (*A*_+_ and *A*_−_), we define the directionality index as |*A*_+_ ™ *A*_−_| / |*A*_+_ + *A*_−_| . By this definition, a directionality index of one indicates activity in one direction only, and a directionality index of zero indicates identical activity in both directions.

We use the same definition of directionality index to compute the directionality of the grid cells in Vector-HaSH, shown in Fig. S27.

## A Supplementary Information

This SI is structured as follows: First, we present the quantification metrics and tools used to generate the numerical results presented in this paper in SI Sec. B. Then, in SI Sec. C-D, we provide theoretical guarantees of the results about Vector-HaSH, first in SI Sec. C focusing on the grid-hippocampal memory scaffold and in SI Sec. D focusing on heteroassociative learning with the sensory cells. In particular, in SI Sec. C we prove that the setup of the memory scaffold described in the main text results in a network with an exponentially large number of robust fixed points with large basins of attraction. Then, in SI Sec. D, we first demonstrate that heteroassociative pseudoinverse learning will result in a memory continuum with the desired properties, and then show that one may feasibly replace the pseudoinverse learning with simpler Hebbian learning and continue to obtain qualitatively similar results.

## B Quantification Metrics

### B.1 Software and Data

The source code for the models presented in this paper will be made available at the following GitHub repository upon acceptance: https://github.com/FieteLab/

### B.2 Mutual Information

In this Appendix, unless otherwise specified, we use 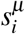 to represent the *i*^th^ bit of the *µ*^th^ pattern stored in the network, and *σ*_*i*_ to represent the *i*^th^ bit of the pattern recovered by the network. Here we primarily consider the case of random patterns such that bits of *s*^*µ*^ are independently sampled from i.i.d. random variables. This allows us to calculate information theoretic quantities for a single bit, and then scale the calculation by the pattern length to obtain the corresponding quantities for entire patterns.

Further, for simplicity of notation in this section, we overload *σ* and *s* to also represent the random variables from which the stored patterns and recovered patterns are being sampled.

We characterize the quality of pattern recovery by a network through the *mutual information* between stored patterns *s* and recovered patterns *σ* . For discrete random variables, the mutual information can be quantified as:

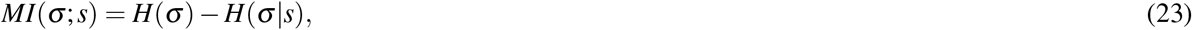

where *H*(*σ* ) is the information entropy of the recovered pattern *σ*,

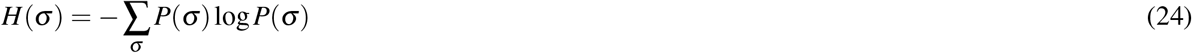

and *H*(*σ*|*s*) is the conditional entropy of the recovered pattern given the stored pattern *s*,

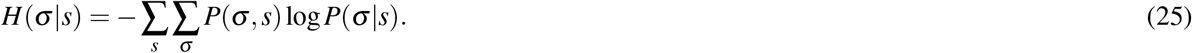

As we now show in the following sections, the mutual information can be explicitly computed for dense and sparse random binary patterns.

#### B.2.1 Dense binary patterns

For unbiased random binary {-1,1} patterns,

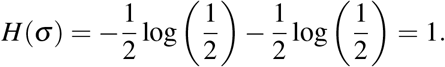

Further, since we assumed that each bit is independent, we obtain

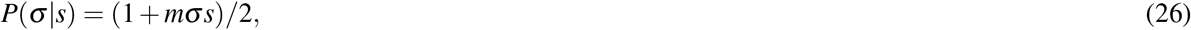

where *m* is the overlap between the stored and recovered pattern, 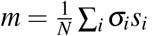 ^98^. Using Eq. (25), this can be used to obtain

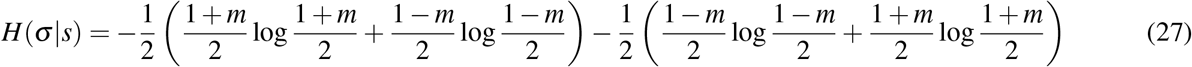

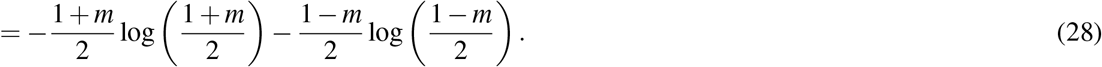

Following Eq. (23) we thus obtain

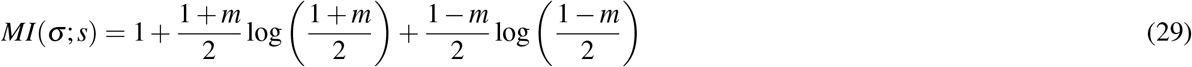

#### B.2.2 Sparse binary patterns

For sparse binary {0,1} patterns, let *p* denote the fraction of “1” bits in the stored pattern (i.e., the average activity of the stored pattern). Let the average activity of the recovered pattern be denoted as *q* = ∑_*i*_ *σ*_*i*_/*N*.

Let *P*_1*e*_ be the probability of error in a bit of *σ* if the corresponding bit of *s* is 1, and *P*_0*e*_ be the error probability in a bit of *σ* if the corresponding bit of *s* is 0. Then,

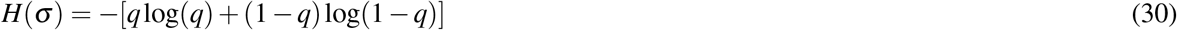

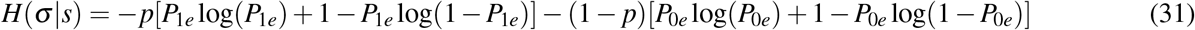

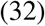

To obtain the probabilities *P*_1*e*_ and *P*_0*e*_, we compute the overlap *m* and the average activity of the recovered pattern *q* in terms of these probabilities as

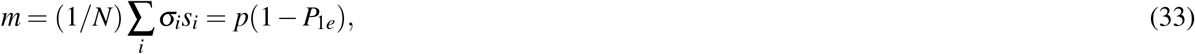

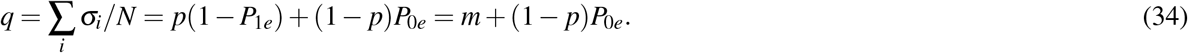

These equations can then be solved to obtain

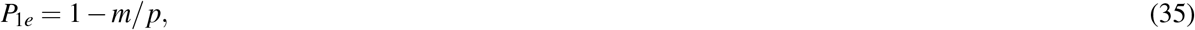

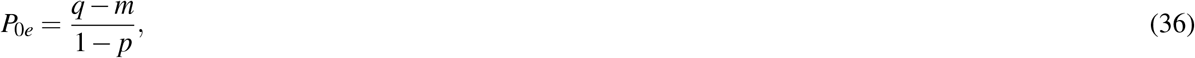

which can then be used to compute *MI*(*σ* ; *s*) using Eq. (23).

#### B.2.3 Continuous random normal patterns

The calculation of mutual information so far has been restricted to the case of discrete binarized patterns. For continuous valued patterns (as in Fig. 3), entropy is ill-defined via Eq. (24). Instead, in this case we can defined the differential entropy as

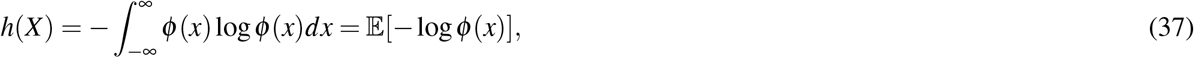

where *φ* (*x*) is the probability density function of the random variable *X*.

For random continuous patterns with patterns are sampled from a normal distribution with zero mean and unit variance, this gives

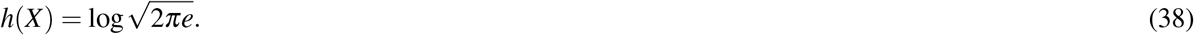

The conditional entropy can similarly be calculated as

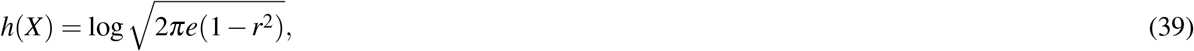

where *r* is the correlation coefficient between *X* and *Y* . This can be used to obtain the mutual information

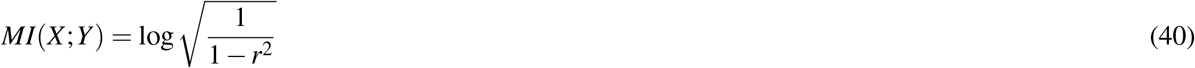

Thus in the case of random normal patterns the mutual information between the stored pattern *s* and the recovered pattern *σ* can be computed directly through the correlation between them using Equation 40 above.

### B.3 Metrics

We quantify the recovery error, i.e., the error between the stored pattern and the recovered pattern in the network by computing the *L*^2^ norm of the difference between stored and recovered patterns. This recovery error is then used to quantitatively apply a recovery threshold to ascertain the capacity of the memory scaffold.

After choosing a recovery threshold (see *Methods*), the capacity of the network is defined as the largest number of stored patterns for which the average recovery error across patterns is below threshold.

## C Theoretical Results on the Memory Scaffold

First, we prove that the memory scaffold network has 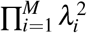 fixed points, while having only 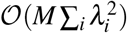 synapses, establishing an exponentially large number of fixed points. Then, we demonstrate that each of these basins are maximally large, and finally demonstrate that these basins are convex, ensuring robustness of basins and protection against adversarial input.

### C.1 Justification for the exponentially large capacity of the memory scaffold

We first provide broad qualitative justification for why the memory scaffold as constructed in Vector-HaSH is capable of storing such a large number of fixed points, then present a mathematical proof in a simplified setting. In this subsection, for ease of notation, we denote the number of phases in the *i*^th^ grid module, 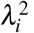, as *l*_*i*_.

Unlike associative memory in the usual context of random patterns (as in the random shuffled hippocampal states considered in Figs. 2d,f and 3d), note that the hippocampal states are determined by a random projection of the structured grid states. As a result, the predefined hippocampal states inherit similar pattern-pattern correlations as the predefined grid cell states. This allows for Hebbian learning to act more efficiently in learning pairwise correlation resulting in a high capacity. Indeed, while in Hopfield networks any given fixed point is destabilized due to interference from other fixed points (resulting in catastrophic forgetting when a large number of fixed points have been memorized), the shared pattern-pattern correlations in the memory scaffold result in the interference terms being positively correlated with each fixed point (which also leads to the scaffold generalization properties Fig. 2f, Sec. C.4).

To show this result more quantitatively, recall that

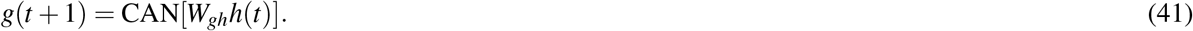

where CAN[*x*] is a nonlinear function that acts independently on each module of grid cells, such that CAN[*x*] will be a vector with exactly one element set to “1” in each of the *l*_*i*_ indices corresponding to each module. and all other elements set to “0”. Further, the element set to 1 in a given module corresponds to the same index as the largest element *x* within that module.

Corresponding to the state *h*^*µ*^ , consider the pattern *h*(*t*) = *h*^*µ*^ + *ζ* , where *ζ* represents a random noise vector. For simplicity, we assume that *ζ* is a continuous-valued vector whose each component is drawn independently from a normal distribution with zero mean and variance *ε*^2^.

From *h*(*t*), we aim to recover *g*(*t* + 1) = *g*^*µ*^ via the mapping *W*_*gh*_. For ease of notation, we denote the prespecified random projection *W*_*hg*_ as *W* .

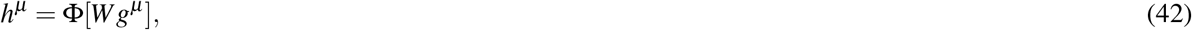

where Φ represents the neural transfer function for the grid to hippocampal synapses, which we implement as a thresholded rectifying function (see *Methods*). We implement *W* such that each element is independently sampled from a prespecified distribution (see *Methods*). Without loss of generality, we can assume that this distribution has zero mean and unit variance, since any transformations of the mean and variance can be absorbed into the nonlinearity Φ.

Now, from the definition of *W*_*gh*_ and *h*^*µ*^ ,

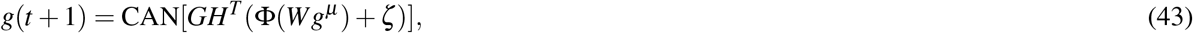

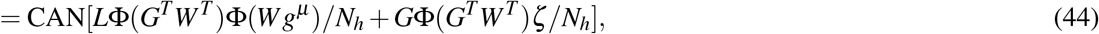

where we have added a scaling factor 1/*N*_*h*_ that leaves the CAN continuous attractor dynamics unchanged, but will be useful for normalization of random variables later in our calculation.

For analytic simplicity, we make the assumption that the nonlinearity Φ in the above equation can be ignored. While this is a gross simplification, the obtained results are broadly consistent with the numerical observations in Fig. 2. This approximates the above equation to

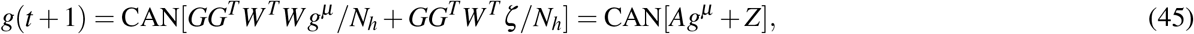

where *A* = (*GG*^*T*^ )(*W*^*T*^*W* /*N*_*h*_), and *Z* = *GG*^*T*^*W*^*T*^ *ζ* /*N*_*h*_.

Since each element of the *N*_*h*_ *× N*_*g*_ matrix *W* was drawn independently from a normal distribution with unit variance, *W*^*T*^*W* (and hence *A*) can be treated as a matrix random variable. Under the distribution of the matrix random variable *A* and the vector random variable *Z* we will compute the probability of *g*(*t* + 1) = *g*^*µ*^ . Note that this simplification of the problem into Eq. (45) has fundamentally relied on Eq. (42), which establishes hippocampal states as being derived from random projections of grid states. Qualitatively, hippocampal states being projections of grid states results in a similarity of state-state relationships between grid states and hippocampal states. As a result, overloading the scaffold network weights with a large number of patterns will not result in loss of previously stored information through interference; instead, pattern interference will re-inforce previously stored patterns (which also results in the strong generalization property, Sec. C.4). In contrast, if the hippocampal states were arbitrarily determined (such as through consideration of random sparse vectors, or sensory-input-dependent vectors), then interference due to storage of additional patterns would result in catastrophic forgetting, as in classic Hopfield memory.

We first focus on the structure of the matrix *GG*^*T*^. This matrix will have a block structure, with the sizes of the blocks determined by the number of phases in each grid module, *l*_*i*_. In particular we write *GG*^*T*^ as

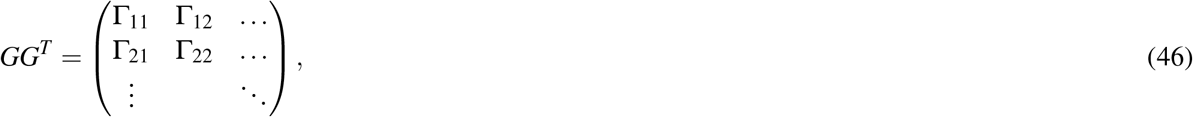

with each Γ_*ij*_ being a submatrix of size *l*_*i*_ × *l* _*j*_. From the structure of the grid code matrix *G*, it follows that

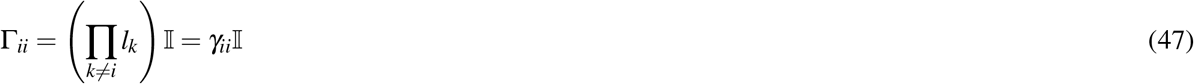

and

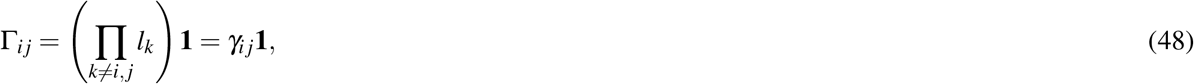

where 𝕀 is an appropriately sized identity matrix, and **1** is an appropriately sized matrix with each element equal to 1. This can be shown by noting that *GG*^*T*^ = ∑_*µ*_ *g*^*µ*^ (*g*^*µ*^ )^*T*^ , and that each *g*^*µ*^ (*g*^*µ*^ )^*T*^ will be a matrix with a single nonzero element equal to 1 in each Γ block of *GG*^*T*^ .

We now compute the distribution of the matrix random variable *W*^*T*^*W* . As argued above, each element of the *N*_*h*_ × *N*_*g*_ matrix *W* can be assumed to be drawn independently from a distribution with zero mean and unit variance. As is justified later, we can assume that these elements are drawn from a normal distribution in particular, since we shall be applying central limit theorem which will wash away particulars of the shape of the distribution.

Thus *W*^*T*^*W* can thus be approximated to have each diagonal element distributed as the sum of the squares of *N*_*h*_ standard normal variables, and each off-diagonal element distributed as the sum of the products of *N*_*h*_ pairs of uncorrelated standard normal variables. Thus

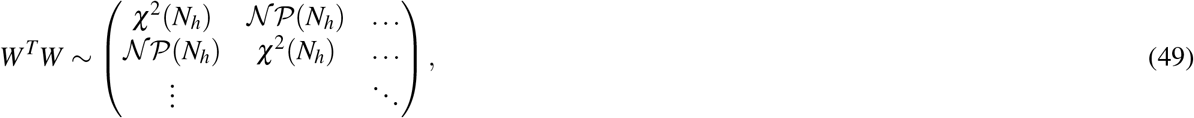

where χ^2^(*N*) is the sum of *N* i.i.d. χ^2^ distributions, and 𝒩 𝒫 (*N*) is the sum of *N* i.i.d. normal product distributions (i.e., the distribution of the product of two i.i.d. standard normal variables). Note that we have suppressed the indices on each matrix element, however it should be noted that each element is an independent sample from the distribution and are identical in distribution but not in value.

In the large *N*_*h*_ limit, each of these matrix elements is the sum of a large number of random variables and can hence be approximated as a normal distribution due to central limit theorem. Thus, χ^2^(*N*_*h*_) ∼ 𝒩 (*N*_*h*_, 2*N*_*h*_), and 𝒩 𝒫 (*N*_*h*_) ∼ 𝒩 (0, *N*_*h*_), where 𝒩 (*µ, σ* ^2^) is a normal distribution with mean *µ* and variance *σ* ^2^.^3^

We thus treat *W*^*T*^*W* /*N*_*h*_ as a matrix random variable with elements on the diagonal being drawn from a distribution𝔻, having unit mean and a variance of 2/*N*_*h*_; and elements on the off-diagonal being drawn from a distribution 𝒟, having zero mean and 1/*N*_*h*_ variance. For ease of calculation, we write this matrix as having a block structure similar to *GG*^*T*^ , given by

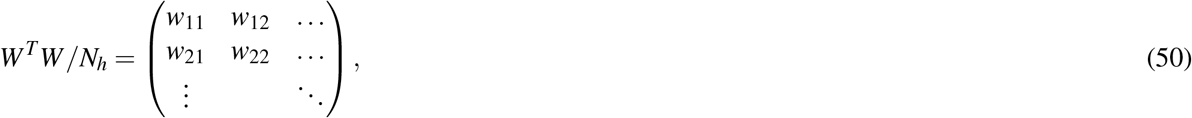

with *w*_*ij*_ being an *l*_*i*_ × *l* _*j*_ matrix such that *w*_*ii*_ has diagonal entries drawn from 𝒟 and off diagonal entries drawn from 𝒪, and *w*_*ij*_ for *ij* being a matrix with all entries drawn from 𝒪

We can now compute the distribution of the elements of *A*. The matrix *A* will have a similar block structure to *GG*^*T*^,

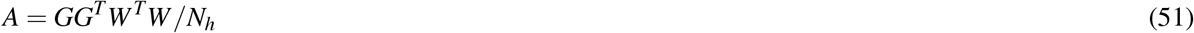

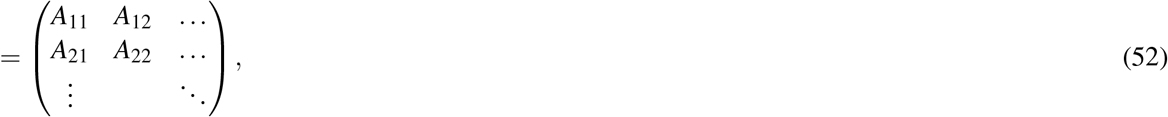

with

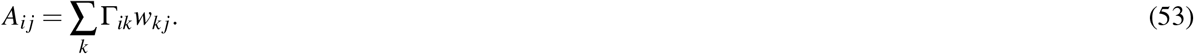

Since each *w*_*ij*_ consists of elements drawn from random normal distributions, the distribution of the matrix variables *A*_*ij*_ can be exactly computed through sums of random normal variables. Even without explicit computation, we can ascertain certain properties of *A* given the symmetry of grid states across module-preserving permutations. In particular, *A*_*ii*_ will be a matrix random variable with diagonal element drawn from an i.i.d. distribution 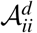, and each off-diagonal element drawn from a different i.i.d. distribution 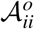. In contrast, *A*_*ij*_ for *ij* will have all elements drawn from an i.i.d. distribution 𝒜_*ij*_.

We first consider *A*_*ii*_.

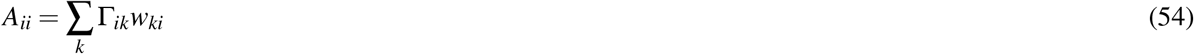

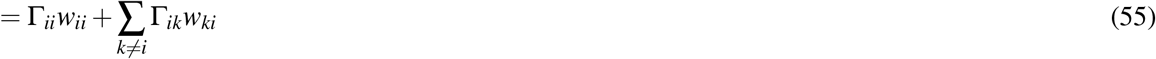

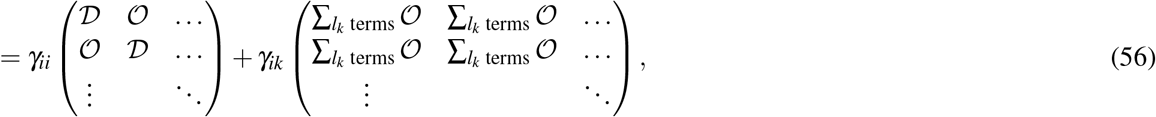

where we have omitted subscripts on individual random variables for simplicity, but it should be noted that each random variable is i.i.d., including the summands in the above expressions. Thus,

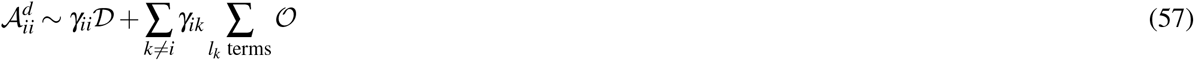

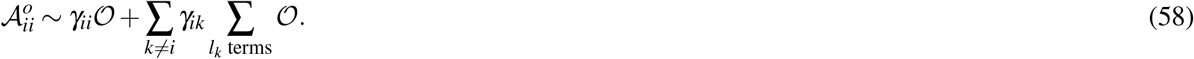

A similar calculation can be done to obtain

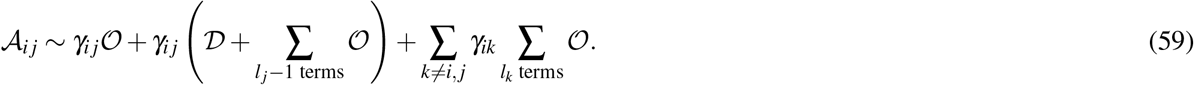

From the same symmetry as in *A*, we can also argue that elements of *Z* can also be split into a similar Ƶ block structure, 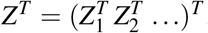, with all 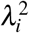 elements of *Z*_*i*_ drawn from an i.i.d distribution *Ƶ*_*i*_. More specifically, *Z* = *GG*^*T*^*W*^*T*^ *ζ* /*N*_*h*_. First note that *W*^*T*^ *ζ* will be a random vector with each element constructed from the sum of *N*_*h*_ i.i.d. normal product distributions multiplied by the scale of *ζ* , i.e., *ε*. Thus *W*^*T*^ *ζ* /*N*_*h*_ is identically distributed to *ε*𝒪. Left multiplying this vector with *GG*^*T*^ we obtain in the *i*^th^ subvector

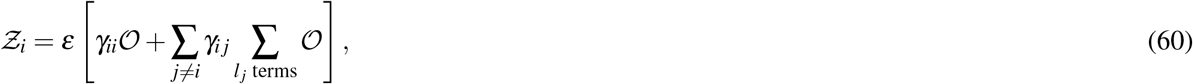

where again we have omitted subscripts on individual random variables for simplicity.

Let the mean and standard deviation of 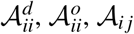 and *Ƶ*_*i*_ be denoted as 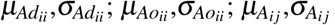; and 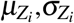 respectively. Since 𝒟 ∼ 𝒩 (1, 2/*N*_*h*_) and 𝒪 ∼ 𝒩 (0, 1/*N*_*h*_), we obtain

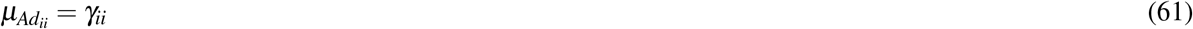

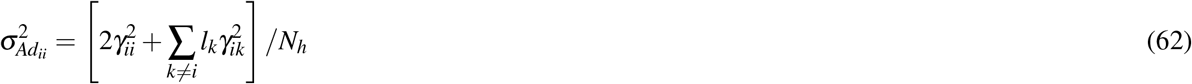

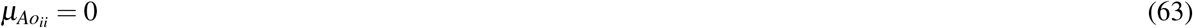

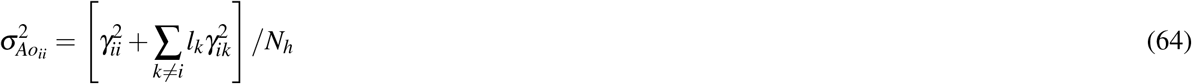

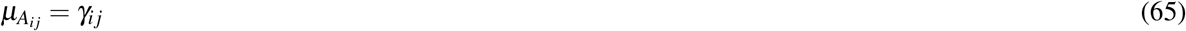

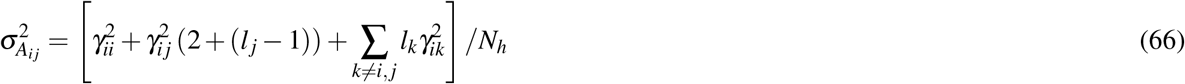

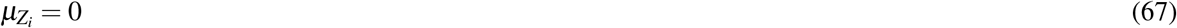

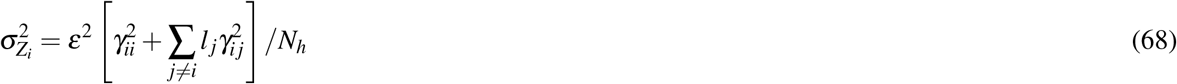

Next, we note the conditions on these blocks to make *g*(*t* + 1) = *g*^*µ*^ , the condition necessary for *g*^*µ*^ to be a scaffold fixed point. Without loss of generality, we assume that *g*^*µ*^ corresponds to the grid state such that each *l*_*i*_ length subvector of *g*^*µ*^ has the first element set to 1 and all others set to zero. The *i*^th^ subvector of *Ag*^*µ*^ + *Z* will then have the first element given by

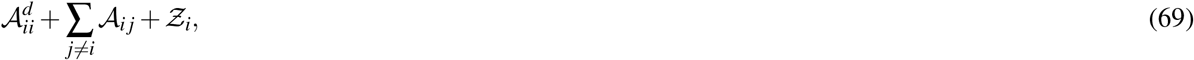

and all other elements given by

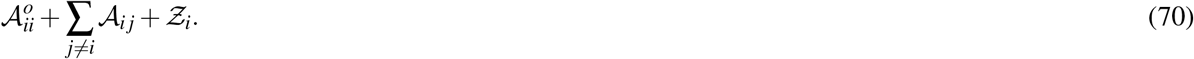

For this module to be correctly reconstructed through the continuous attractor network dynamics, we require that the first element of the subvector to be larger than the others. Thus, the probability of the correct reconstruction is given by

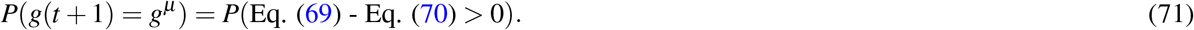

As seen earlier, each of these random variables are being drawn from a normal distribution (due to central limit theorem in the limit of large *N*_*h*_). In terms of the parameters of these normal distributions, Eq. (71) can be written as

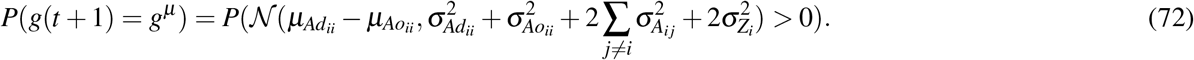

For ease of notation, we define

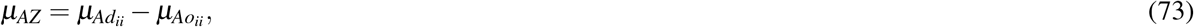

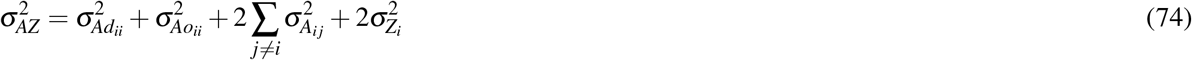

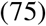

such that the right-hand side of Eq. (72) is equal to 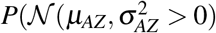. This can then be computed as

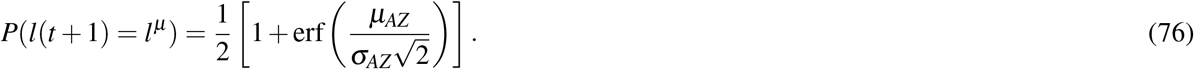

The above-derived expressions for the terms in *µ*_*AZ*_ and *σ*_*AZ*_ can be simplified to

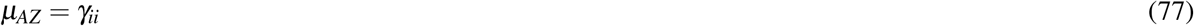

and

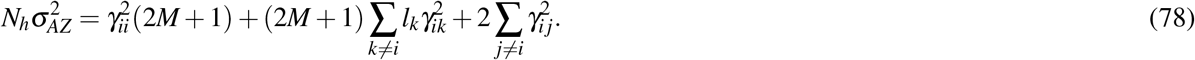

Recall that *γ*_*ii*_ = *P*/*l*_*i*_ and *γ*_*ij*_ = *P*/(*l*_*i*_*l* _*j*_), for *P* = Π_*i*_ *l*_*i*_. Thus, the ratio 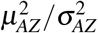 simplifies to

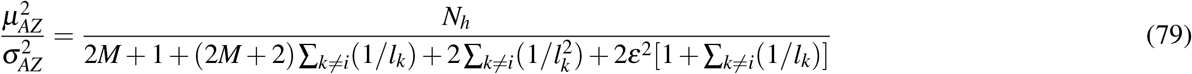

Inverting the obtained expression allows for computation of 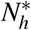,

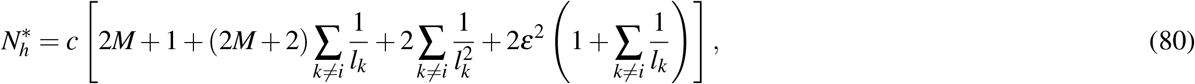

where *c* = 2 [erf^−1^(1−2*P*)]2 and *P* is the threshold selected for accuracy of the recovered pattern. This allows us to estimate the critical number of hippocampal cells (as a function of the number of grid cell modules, *M*, the period of the modules 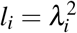,and the input noise *ε*) beyond which the hippocampal memory scaffold stores all grid-hippocampal states as fixed points.

If the number of grid cells far exceeds the number of modules (as would be expected^81^), then *λ*_*k*_ ≫ *M* and thus *l*_*k*_ ≫ *M*_2_ and the summands in Eq. (80) can all be ignored. This makes 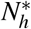 asymptotically independent of the grid periods, and is given by 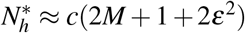. This can be seen qualitatively in Fig. 2, where for a fixed *M*, the critical number of hippocampal cells 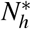 approaches a constant with increasing *N*_*g*_ (and hence increasing *l*_*k*_). Moreover, if *ε* ≪ 1 and *M* ≫ 1, then 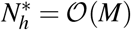. This has been verified qualitatively in Fig. 2, where 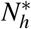 increases linearly with increasing number of modules *M*. Note that due to the simplifications necessary for this analytic result, 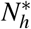 obtained from Eq. (80) are not directly comparable to numerics in Fig. 2, however the above-mentioned qualitative trends all seem to hold.

These results thus demonstrate a crucial property of the hippocampal memory scaffold network — it has 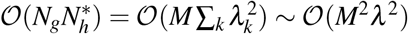 synapses while having 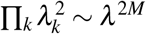 fixed points. Thus, the number of fixed points grows exponentially faster than the number of synapses in the network, resulting in the network being useful as a memory scaffold as in MESH^86^.

### C.2 Memory Scaffold has Maximally Sized Basins of Attraction

Due to the symmetries in grid code, we argue here that the memory scaffold in Vector-HaSH has no spurious fixed points, and has convex, maximally sized basins of attraction that are equal in volume.

First, we note that as a result of the CAN dynamics in the grid layer (cf. Eq. (4)), the only possible grid states are the Π_*i*_ *K*_*i*_ modular one-hot states. Correspondingly, the hippocampal states (determined by random projections of the grid states) must then also be one of the Π_*i*_ *K*_*i*_ states, establishing that no spurious fixed points can arise.

Thus, the union of the basins about each of the predefined fixed points of the grid-hippocampal scaffold cover the entire space 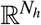. Note next that each *h*^*µ*^ are equivalent, i.e., there is no special *µ* since each *g*^*µ*^ is equivalent up to a module-preserving permutation of bits and *h*^*µ*^ are determined by a random projection of *g*^*µ*^ . Thus, 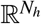 must be partitioned into basins with equal volume that are maximally large (and hence are of the same volume as the Voronoi cell about these fixed points).

### C.3 Convexity of Scaffold Basins

The existence of maximally sized equi-volumed basins around each predefined scaffold fixed point, as we have shown, is not sufficient to guarantee robustness to noise. A large basin could in principle have some boundaries that come arbitrarily close to the fixed points – such a situation holds for instance when a system is susceptible to adversarial inputs, where a very small perturbation of the input leads to a very different classification as an output. Noise robustness requires a second condition, that of basin convexity. Here we demonstrate that the obtained basins are convex, and thus the large basins must result in basin boundaries that are well separated from the fixed points themselves.

We are interested in the basins in the space 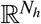: the hippocampus receives sensory input from the sensory layer, mediating the recall of scaffold states. Thus, noise robustness will hence be required there. The broad idea of the proof is as follows: first, we demonstrate that perturbations in the hippocampal latent space are equivalent to considering real-valued perturbations with small magnitudes in the grid-cell layer latent space. Then we show that the continuous attractor dynamics on grid cells result in convex basins in the grid-cell space, which directly translates to convex basins in the hippocampal space.

Consider a hippocampal population vector given by a small perturbation to a predefined hippocampal state fixed point *h*^*µ*^ , which we denote as *h* = *h*^*µ*^ + *ε*. Let *δ* denote the magnitude of the perturbation *ε*. This hippocampal state is projected onto the grid cells through *W*_*gh*_ to obtain 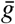 before the continuous attractor dynamics, where

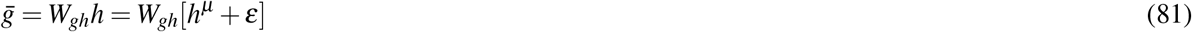

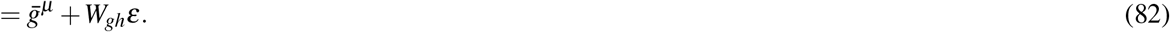

Note that *W*_*gh*_*ε* will have a magnitude of approximately *δ* times the magnitude of 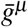, and further, the nonzero elements of *ε* are uncorrelated with *h*^*µ*^ , and hence *W*_*gh*_*ε* can be treated as an independent small real-valued perturbation to 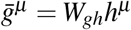.

If we can now show that the continuous attractor dynamics on grid cells has a convex basin, that would indicate that all points near *g*^*µ*^ map to *g*^*µ*^ , and since points near *h*^*µ*^ map to points near *g*^*µ*^ , this would imply convexity of basins in *p*-space.

The symmetry of the grid code implies that it will suffice to show that the basin about any one fixed point is convex. Without loss of generality, we choose the fixed point *g*^*µ*^ as the grid population vector with the first bit in each module set to 1 and all other 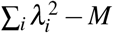 bits set to 0. Let *x* and *y* be two vectors within the continuous attractor dynamics of *g*^*µ*^ , i.e., *CAN*[*x*] = *CAN*[*y*] = *g*^*µ*^ . Thus, for the *k*^th^ module, *x*_*k*;1_ > *xk*,*I* and *yk*;1 > *yk*,*I* for *i* > 1. Adding the two inequalities with coefficients *a* and (1 ™ *a*), we obtain *ax*_*k*;1_ + (1 ™ *a*)*y*_*k*,1_ > *ax*_*k*;*i*_ + (1 ™ *a*)*y*_*k*,*i*_ for all *i* > 1 for 0 ≤ *a ≤* 1. Thus, continuous attractor dynamics (which enforce modular winner-take-all dynamics) map the *k*^th^ module of *ax* + (1 ™ *a*)*y* to the *k*^th^ module of *g*^*µ*^ . Since this holds for all *k*, thus CAN[*ax* + (1 ™ *a*)*y*] = *g*^*µ*^ . Hence, for any two vectors *x* and *y* in the basin of *g*^*µ*^ , all vectors on the line from *x* to *y* also lie in this basin. By definition, this makes the basin of *g*^*µ*^ , and as argued earlier this imposes convexity of basins in the hippocampal cell space.

### C.4 Scaffold weights can be learned on a vanishing fraction of all states

As shown in Fig. 2, when the hippocampus-to-grid cell synaptic weights (*W*_*gh*_) are learned on a small number of grid states, the scaffold is able to generalize: all grid states become stable fixed points of the scaffold dynamics. Thus an animal only needs to traverse small regions in space after which the grid-hippocampal scaffold is recurrently stabilized for all other states. Here we show that in the large *N*_*h*_ limit, it will suffice to train on only 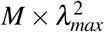 patterns for stabilization of the complete scaffold, where *λ*_*max*_ is the largest period of any grid module.

Similar to Sec. C.1, we make the grossly simplified assumption that the nonlinearities in the hippocampus can be ignored. The grid cell state would then evolve as

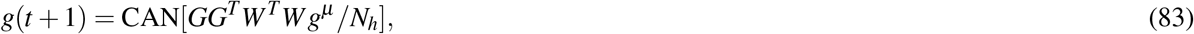

where again we add an 1/*N*_*h*_ scaling factor that renders the continuous attractor network dynamics unchanged. For *g*^*µ*^ to be a fixed point of the recurrent scaffold dynamics, we thus require that *g*(*t* + 1) be equal to *g*^*µ*^ . Unlike Sec. C.1, we assume here that *G* is the matrix constructed by appending grid cell population vectors over only the first *N*_*patts*_ number of states (rather than over the entirety of scaffold states).

As argued earlier in Sec. C.1, for large *N*_*h*_, the matrix *W*^*T*^*W* /*N*_*h*_ can be considered to be a random variable with i.i.d 𝒩 (1, 2/*N*_*h*_) random variables on the diagonal, and i.i.d 𝒩 (0, 1/*N*_*h*_) entries on the off-diagonal. In the limit of *N*_*h*_ → ∞, these distributions tend to Dirac delta distributions we can thus treat *W*^*T*^*W* /*N*_*h*_ as simply being an identity matrix. Thus, in this limit, it suffices to examine the scaffold fixed points under the dynamics

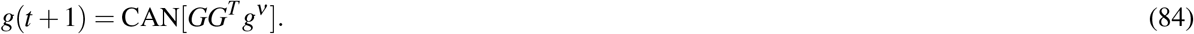

As earlier, we write *GG*^*T*^ as a block matrix

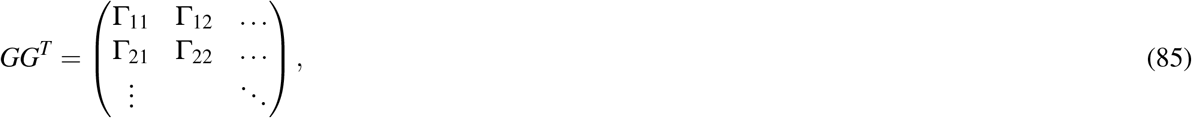

with each Γ_*ij*_ being a submatrix of size 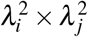. We define a construct a sequence of patterns *g*^*µ*^ as follows: let the first pattern *g*^1^ be such that the first element in each 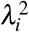 subvector is set to one, and all other elements set to zero. Then, each successive pattern shifts the active element by one, modulo the total number of elements in the subvector 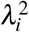. Mapped to real space, this corresponds to the sequence of locations shown in Fig. S14a (*top left*). As we will now show, setting *W*_*gh*_ based on only 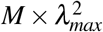 grid patterns will suffice to stabilize all patterns.

Note that

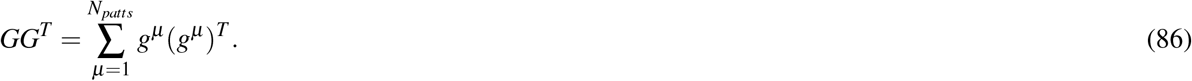

Each *µ* term of this summation, *g*^*µ*^ (*g*^*µ*^ )^*T*^ will be a matrix with exactly one ‘1’ in each block Γ_*ij*_, at the location (*µ* mod 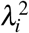 mod 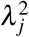), and will be zero everywhere else.

For *i*≠ *j*, the periods *λ*_*i*_ and *λ*_*j*_ are coprime. In this case, we can see that if *g* and *g* contribute a ‘1’ at the same location (*m, n*) in Γ_*ij*_ then *µ* − *ν* must be a multiple of 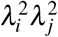. This can be seen since *m* = *µ* mod 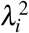, thus *µ* = *m* mod 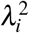. Similarly *ν* = *m* mod 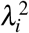, implying that *µ* − *ν* = 0 mod 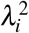. Similar reasoning leads to *µ* − *ν* = 0 mod 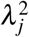 and thus *µ* − *ν* = 0 mod 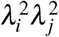. Crucially, this means that if *µ*≠ *ν*, then *µ* − *ν* must be at least 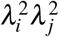, which is equal to the number of elements in Γ_*ij*_.

Thus, if both *µ* and *ν* contribute a ‘1’ to (*m, n*) in Γ_*ij*_, it must be that all other elements in Γ_*ij*_ have been increased by 1 due to patterns between *µ* and *ν*. In essence, elements of Γ_*ij*_ increase sequentially through increasing terms in the summation Eq. (86). Starting from all elements at zero before any learning, all patterns increase to 1 one-at-a-time up to 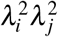, all patterns increase up to 2 through the next 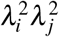 patterns and so on. Crucially, at any point during learning, the largest element of Γ_*ij*_, which we denote max Γ_*ij*_, can differ from the smallest element of Γ_*ij*_, which we denote min Γ_*ij*_, by at most 1.

Next, we observe that for Γ_*ii*_, the pattern *µ* contributes a ‘1’ at the location (*µ* mod 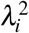, *µ* mod 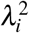). This leads to two observations: first, that Γ_*ii*_ will have nonzero entries only on its diagonal; second, the smallest element on the diagonal will be 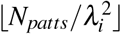 and the largest element on the diagonal will be 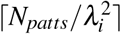.

Now, consider the matrix *GG*^*T*^ constructed using Eq. (86), trained through the first *N*_*patts*_ patterns. We apply Eq. (84) for a given *g*^*µ*^ , for *µ* that need not be within {1… , *N*_*patts*_} . Note *g*^*µ*^ has a 1 at only one location per module. Thus the *i*^th^ subvector of *GG*^*T*^ *g*^*µ*^ can have values as small as 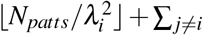 min Γ_*ij*_ at the index where *g*^*µ*^ equals 1 in the *i*^th^ module; and, it can have values as large as ∑ _*j*≠*i*_ max Γ_*ij*_ at the other entries. For this subvector to map to *g*^*µ*^ under the continuous attractor network dynamics, we thus require

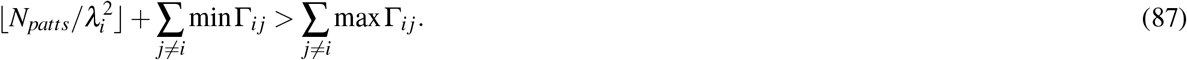

Thus,

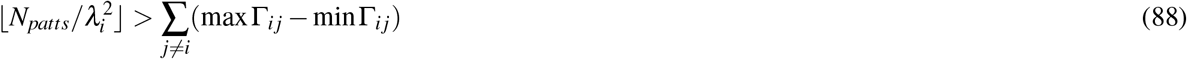

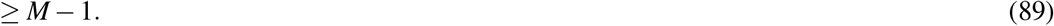

Thus 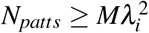. Since the correct subvector needs to be recovered for all modules, thus 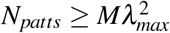 for stabilization of *all* grid states *g*^*µ*^ in the large *N*_*h*_ limit.

Note that the proof above relies on the particular ordering of grid and hippocampal states described above. As we demonstrate, this ordering is optimal, and no other ordering of grid states can result in ‘faster’ generalization to all scaffold fixed points. To see this, note that we showed above that the particular ordering choice made ensures that the largest and smallest elements of Γ_*ij*_ can differ by at most 1. Moreover, this difference of at most one resulted in the generalization result proved above. Correspondingly, any other ordering that maintains this difference between the largest and smallest elements of Γ_*ij*_ will also demonstrate generalization to all scaffold states at 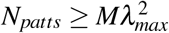. Faster generalization could only be possible if the elements of Γ_*ij*_ we all identical, leading to a difference of zero. This is however impossible, since the sum of elements in Γ_*ij*_ is equal to *N*_*patts*_, which is increasing in steps of 1 and is thus not always divisible by the number of entries in Γ_*ij*_, i.e., 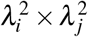. Conversely, any other ordering, will result in a potentially larger difference between the smallest and largest elements of Γ_*ij*_, which (following Eq. 88) will thus require a larger number of patterns to generalize to all scaffold states.

However, as noted in Fig. 2g, other contiguous orderings of grid states result in generalization upon learning an approximately similar number 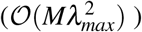 of patterns.

## D Theoretical Results on Heteroassociative Learning

Here we demonstrate that pseudoinverse learning first perfectly recovers the hippocampal states provided that *N*_*s*_ > *N*_*patts*_ (in the noise-free case). Following the memory scaffold results proven earlier, reconstruction of the correct hippocampal states then results in correct retrieval of the corresponding label layer states. Next, we prove that for *N*_*patts*_ < *N*_*h*_, the reconstructed feature layer states are also perfectly reconstructed, and for larger *N*_*patts*_ the overlap of the stored and recovered patterns decays gracefully as described in the main text. We then prove that given an ideal memory scaffold, heteroassociative *Hebbian* learning is also sufficient to obtain a memory scaffold with the same qualitative properties, with only a smaller prefactor on the memory capacity.

### D.1 Perfect Reconstruction of Hippocampal States Through Heteroassociative Pseudoinverse Learning

The projection of the learned sensory inputs onto the hippocampus is given by *W*_*hs*_*S* = *HS*^+^*S* = *H*Λ_*S*_, where Λ_*S*_ = *S*^+^*S* is an orthogonal projection operator onto the range of *S*^*T*^ . If *N*_*s*_ ≥ *N*_*patts*_, *S* has linearly independent columns, and Λ_*S*_ = 𝕀, the identity matrix. Thus, *W*_*hs*_*S* = *H*, i.e., cuing any memorized patterns results in accurate reconstruction of the corresponding hippocampal scaffold state

However, to examine Vector-HaSH as an associative memory, it is necessary to examine to reconstruction of the correct hippocampal scaffold state when cued with noisy or corrupted versions of the memorized patterns as well. Following the results of SI Sec. C.3 and C.2, we see that the memory scaffold has maximally large convex basins of attraction. Note that once *W*_*hs*_ has been trained with pseuodo-inverse learning, the mapping from the sensory layer to the hippocampal layer is simply a linear transformation, that maps stored sensory patterns to their corresponding hippocampal scaffold states. Thus, the regions in sensory space that map to a given scaffold state must simply be a lower-to-higher-dimensional linear transformation of convex basins in the scaffold space about the chosen state.

Hence, the basins of attraction for a given scaffold state must be convex regions in the sensory input state that include the sensory pattern that has been associated with that scaffold state.

### D.2 Perfect Reconstruction of *N*_*h*_ Sensory States Through Heteroassociative Pseudoinverse Learning

To show that up to *N*_*patts*_ ≤ *N*_*h*_ sensory inputs can be perfectly reconstructed through Vector-HaSH, we require that the matrix of fixed point hippocampal states *H* be strongly full rank, i.e., all submatrices formed by considred a subset of columns of the matrix *H* are full rank. While we do not rigorously prove that *H* is strongly full rank, we hypothesize that this is true if the hippocampal states: 1) are determined by random projection from grid cells and 2) involve some nonlinear transformation of the grid inputs (we posit that almost every nonlinear transformation in the space of all functions is sufficient, without fine-tuning the functional form). For instance, simple rectification with threshold is sufficient, for a wide range of activation thresholds (SI Figs. S12,S13; in contrast, a linear hippocampal layer does *not* result in strongly full rank hippocampal states, SI Fig. S14), since rank(*W*_*hg*_*G*) = rank(*G*) = *N*_*G*_ ™ *M* + 1^115^, which does not need to be as large as *N*_*h*_.

The intuition for this claim is as follows: applying a thresholded rectifying function, *H* = ReLU[*W*_*hg*_*G* ™ Θ], effectively acts as an independent random perturbation to the elements of *H*. Assuming that these perturbations are truly random, *H* (and submatrices of *H* formed by selecting varied numbers of fixed points) will become full rank. This is numerically verified in Fig. S12, where the rank can be seen to be min(*N*_*patts*_, *N*_*h*_).

We can now show that the “knee” of the Vector-HaSH memory continuum must be at *N*_*h*_, with *N*_*patts*_ ≤ *N*_*h*_ sensory states being perfectly reconstructed.

The projection of the hippocampus states onto the sensory layer is given by *W*_*sh*_*H* = *SH*^+^*H* = *S*Λ_*H*_, where Λ_*H*_ = *H*^+^*H* is an orthogonal projection operator onto the range of *H*^*T*^ . Since *H* is strongly full rank (as justified above), thus for up to *N*_*patts*_ ≤ *N*_*h*_, the projection operator Λ_*H*_ will equal 𝕀, the identity matrix. Thus *W*_*sh*_*P* = *S*.

### D.3 Mutual information recalled in Vector-HaSH scales as 1/*N*_*patts*_

Let 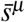 be the reconstruction of pattern *s*^*µ*^ in the feature layer before the application of the sign nonlinearity in Eq. (12), i.e., 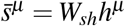. Correspondingly, let 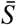 be the matrix constructed with 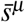 as its columns, i.e., 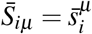 . In this notation, we wish to prove that 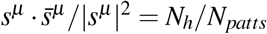.

As earlier, 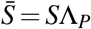. Since *N*_*patts*_ > *N*_*h*_, rank(*H*) = *N*_*h*_, and the projection operator Λ_*H*_ is thus no longer an identity operator. Instead, Λ_*H*_ projects on to the *N*_*h*_-dimensional hyperplane 𝒫_*H*_ spanned by the rows of *H*. Notationally, let 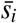 be the vector corresponding to the *i*^th^ row of 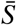, and similarly, let *s*_*i*_ be the vector corresponding to the *i*^th^ row of *S*. In this notation, the vectors 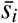 (i.e., the rows of 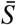) are the vectors obtained by projecting *s*_*i*_ (i.e., the rows of *S*) onto 𝒫_*H*_.

By construction *s*_*i*_ are *N*_*patts*_-dimensional random vectors with no privileged direction. Thus, |*s*_*i*_|^2^, the squared magnitude along each dimension, will on average be equally divided across all dimensions. Hence, on average, the component of *s*_*i*_ projected onto 𝒫_*H*_ (i.e., 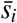) will have a squared magnitude of *N*_*h*_|*s*_*i*_|^2^/*N*_*patts*_ and thus 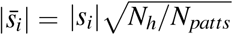 . However, 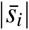 is also the cosine of the angle between *s*_*i*_ and the hyperplane 𝒫_*H*_, and hence averaged over *i*,

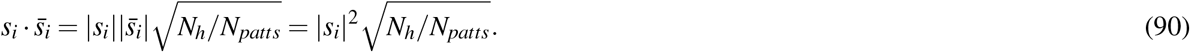

Note that 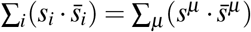, and ∑_*i*_ |*s*_*i*_|^2^ = ∑_*µ*_ |*s*^*µ*^ |^2^. Thus the above equation can be rewritten as

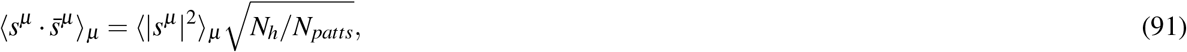

where ⟨⟩_*µ*_ denotes an average over all patterns *µ*.

In the notation of Eq. 29, this gives 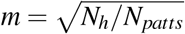. In the limit of small *m*, note that log(1 + *m*) ≈ *m*, and the right-hand side of Eq. 29 can simply be approximated in the asymptotic limit as

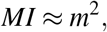

and thus mutual information scales as

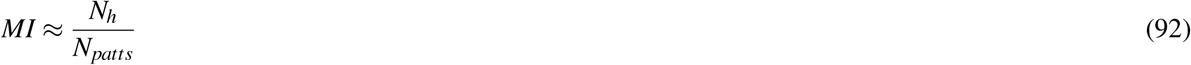

As a consequence of this result, note that since the mutual information is always positive and only smoothly degrades 1 with increasing *N*_*patts*_, thus the recovered state only gradually moves away from the true pattern in sensory space (cf. Fig. 3g). For random uncorrelated sensory patterns, the boundary of the Voronoi cell about a memorized pattern corresponds to the boundary at which no information is being recovered specific to one particular pattern. Thus, an always positive mutual information indicates that the recovered pattern always remains with the Voronoi cell corresponding to the true pattern, i.e., the recovered pattern is always closer to the correct patterns as compared to any other pattern.

### D.4 Space- and time-complexity of memory in Vector-HaSH

We show in SI Sec. D.1 that the number of sensory cells can scale as fast or faster than the maximal scaffold capacity 𝒪 (*K*^*M*^). Thus, *N*_*s*_ ≫ *N*_*h*_, *N*_*g*_, and thus the number of synapses in the model, # synapses = 2*N*_*h*_(*N*_*s*_ + *N*_*g*_) + *M* ∗ *K*^2^ is dominated by 2*N*_*s*_ ∗ *N*_*h*_. Moreover, the number of hippocampal cells is asymptotically constant for large *K* at a fixed *M* (Fig. 2d *right*, SI Fig. S2), and thus the number of synapses scales as 𝒪 (*N*_*s*_). Further, the number of nodes, *N*_*s*_ + *N*_*h*_ + *N*_*g*_ also scales as 𝒪 (*N*_*s*_). Since the number of patterns perfectly reconstructed *N*_*h*_ is constant (with respect to the asymptotic scaling of the number of synapses at a fixed *M*), the continuum in Vector-HaSH ranges from storing𝒪)(1) patterns with 𝒪 (*N*_*s*_) =𝒪 (# synapses) each, up to storing 𝒪 (*N*_*s*_) =𝒪 (scaffold size) patterns with positive recovered information.

The memory storage requirement for Vector-HaSH is equal to the number of synapses in the model, which as noted above scales as 𝒪 (*N*_*s*_*N*_*h*_). This permits storage of *N*_*h*_ patterns perfectly, consisting of a total of *N*_*s*_*N*_*h*_ bits of information. Thus, as in Hopfield and Hopfield-like networks, the total information stored and recalled in Vector-HaSH scales as the number of synapses. The time complexity for perfect recovery of all 𝒪 (*N*_*s*_*N*_*h*_) = 𝒪 (# synapses) bits of information scales as 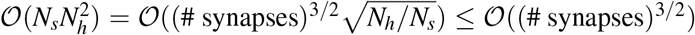. In contrast, for Hopfield and Hopfield-like networks, the time complexity for recovery of 𝒪 (# synapses) scales as 𝒪((# synapses)^3/2^). Vector-HaSH thus has an asympototically faster time complexity than Hopfield-like networks for recovery for the same number of total bits of information (when normalized by the number of synapses in the model).

When the number of patterns stored is larger and scales with the number of sensory cells, *cN*_*s*_ for 0 < *c* ≤ 1, Vector-HaSH partially recovers the stored information (Fig. 3). In this regime, Vector-HaSH has additionally improved time and space complexity as compared with the number of synapses: a space requirement of only 𝒪 (# synapses) and a time complexity of only 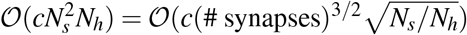 is needed to store an input of 𝒪 (*c*(# synapses) × (*N*_*s*_/*N*_*h*_)) bits of information.

The above scalings can also be reinterpreted in terms of the total information, *I*, stored in the networks (where for perfect recovery *I* = *N*_*h*_*N*_*s*_ = # synapses, and for partial recovery 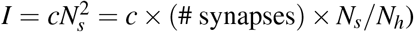. For perfect recovery the space complexity requirements scale as *I*, and time complexity scales as 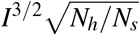. For partial recovery the space complexity scales as *IN*_*h*_/(*cN*_*s*_), and time complexity scales as 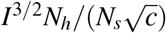.

### D.5 One-Step Heteroassociation Leads to a Memory Continuum even with Hebbian Learning

The memory continuum in Vector-HaSH is a result of the one-step heteroassociation from the hidden to the feature layer, given a memory scaffold that perfectly recovers the hidden states. This holds irrespective of the nature of heteroassociation (pseudoinverse learning or Hebbian learning).

Here we consider a simpler scenario where *W*_*sh*_ is trained through Hebbian learning and the hippocampal states are assumed to be correctly reconstructed (corresponding to pseudoinverse learning from *S* to *H*).

We assume here that the sensory patterns being stored are random unbiased binary vectors drawn uniformly from 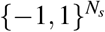. If the weights from the hippocampal cells to the sensory inputs, *W*_*sh*_ are trained using Hebbian learning:

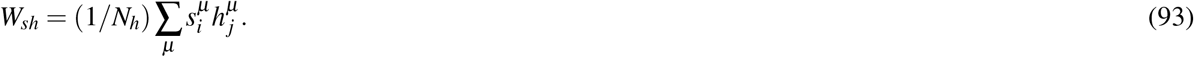

To evaluate the accuracy of a recovered sensory state through this weight matrix, we estimate the probability that the *i*^th^ bit of *s*^*ν*^ is recovered correctly. Since we assume in this simplified scenario that the hippocampal state *p*^*ν*^ has been correctly recovered:

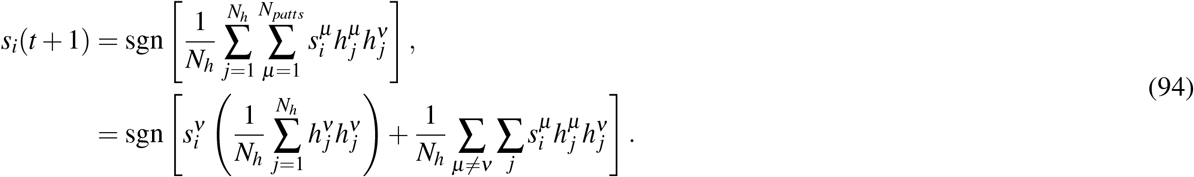

Here we have separated the pattern *ν* from all the other patterns. Next, we multiply the second term on the right-hand side by a factor 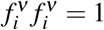, and pull 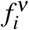 out of the argument of the sign-function since 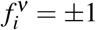.

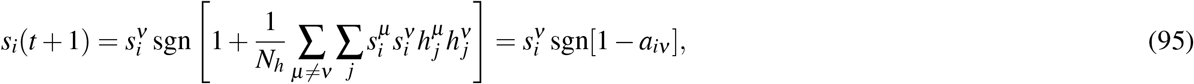

where

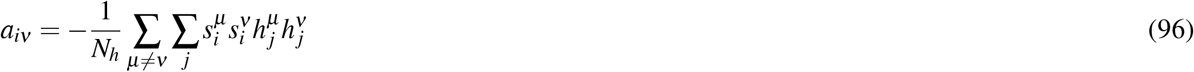

Successful recovery of the *i*^th^ bit of *s*_*ν*_ will occur if 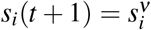, which holds provided that *a*_*iν*_ > 1. The probability of an error in a given randomly chosen bit can thus be calculated as the probability that *a*_*iν*_ > 1.

Since the sensory states were assumed to have been drawn uniformly from 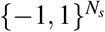, we can treat the product 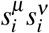 as being +1 or −1 with equal probability. We assume that the distribution of hippocampal cell activity has mean *µ*_*p*_ and variance 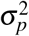. Further, assuming that 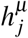 and 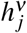 can be treated as independent random variables, the product 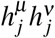 would then have mean 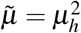 and variance 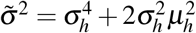. Accounting for the random sign introduced by 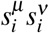, the summand in Eq. (96) can then be treated as a random variable *X* with mean 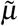 and variance 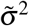 with probability 0.5, and mean 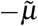 and variance 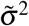 with probability 0.5. For large *N*_*h*_ and *N*_*patts*_, we are summing over a large number of random variables — thus due to the Central Limit theorem, the precise details of the distribution will not matter, apart from an estimate of its mean and variance. By symmetry, the mean of *X* will be zero. This variance can be calculated to be 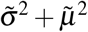.

Summing over (*N*_*patts*_−1)*N*_*h*_ ≈ *N*_*patts*_*N*_*h*_ terms in Eq. (96), evaluated through central limit theorem, thus gives a normal distribution, with zero mean, and variance 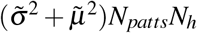. This gives

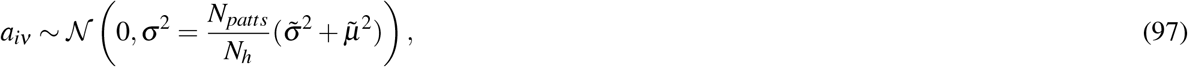

with

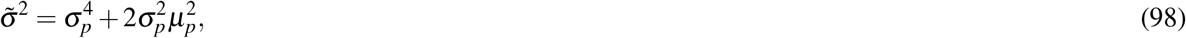

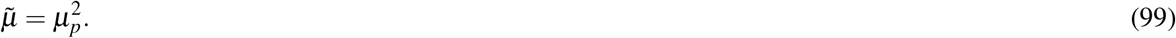

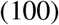

The probability of error in the activity state of neuron *i* is therefore given by:

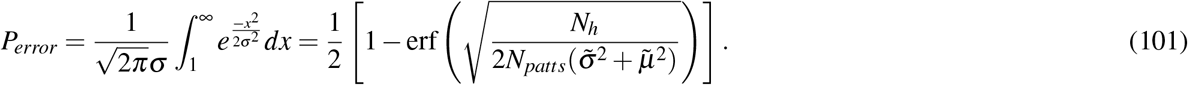

Thus the probability of error increases with the ratio *N*_*patts*_/*N*_*h*_. The mutual information between the stored and recovered Sensory States is then:

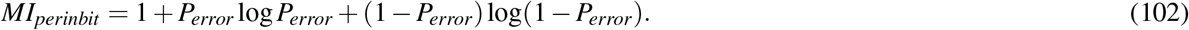

Since erf(*x*) ≈ *x* and *log*(1 + *x*) ≈ *x* for small *x*, in the limit of a large *N*_*patts*_ the above expression can be approximated to

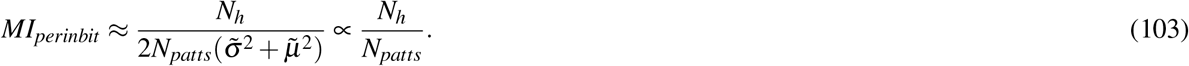

This asymptotic scaling is verified in Fig. S8

### D.6 Two-sided threshold for familiarity detection involves a small feed-forward network

Calculation of the mean of the hippocampal population vector and comparison with the thresholds *θ*_1_ and *θ*_2_ can be performed by a simple, single-hidden-layer feed-forward network with two hidden units. The network receives inputs from the hippocampus *h*:

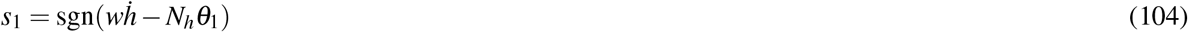

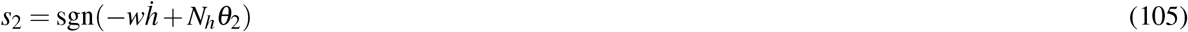

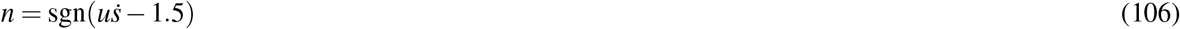

where *w* and *u* are both ones vectors, and *s* = (*s*_1_, *s*_2_) are the outputs of the two hidden layer units. *n* = −1 implies a novel pattern, and *n* = − 1 implies a familiar pattern. This simple circuit permits novelty detection within a biologically plausible setup.

### D.7 Pseudoinverse learning of mappings from sensory states to velocities and memory palace items

Similar to pseudoinverse learning done from hippocampal cells to sensory cells, an exactly equivalent mathematical theory applies for pseudoinverse learning from sensory cells to the one-hot representation of velocities associated with each sensory state in 6 (see *Methods* for details of the learned velocity representations). In particular, as seen in Sec. D.2, pseudoinverse learning of a matrix *W*_*yx*_ that maps from a layer *X* of dimensionality *N*_*x*_ to a layer *Y* of dimensionality *N*_*y*_ is successful in exact recovery of all patterns in layer *Y* when learning up to rank(*X* ) patterns. In the case that rank(*X* ) = *N*_*x*_, as is the case when learning mappings from *S* to *P* and vice-versa (Sec. D.1, D.2), the number of learned patterns that can be perfectly reconstucted is simply *N*_*x*_.

Thus, learning mappings from sensory cells to either velocity representations or memory palace task items will be exactly successful for up to *N*_*s*_ velocities of memory palace items provided that the mappings are being learned from patterns that form a full rank matrix. However, as seen in Fig. 6, mappings must be learned from the reconstructed sensory states rater than the ground truth sensory states, since with increasing number of stored patterns the reconstructed states deviate from the ground truth states.

Thus, even if the sensory states form a full rank matrix, for successful mappings, it will be necessary that the rank of the recovered sensory states must be *N*_*s*_. Following the results presented in Sec. D.3, it would appear that the recovered sensory patterns would form a matrix of rank *N*_*h*_, the dimensionality of the hyperplane 𝒫_*H*_. However, this would only be the case if the recovered sensory states were obtained directly from *W*_*sh*_*H* without any additional nonlinearity. For the case of binary sensory states, the recovered sensory patterns are given by sgn[*W*_*sh*_*H*]. This sign nonlinearity in effect behaves like a small random perturbation to each of the *N*_*s*_ bits of *W*_*sh*_*H*, rendering the reconstructed sensory states to be full rank (assuming that the ground-truth sensory states matrix is full rank). Thus, reconstruction of velocity mappings and memory palace task items (or indeed any other readout from sensory the sensory cells) will be successful for up to *N*_*s*_ patterns, provided that the mappings are being learned from the reconstructed sensory states (rather than the ground truth sensory states).

**Figure S1.**
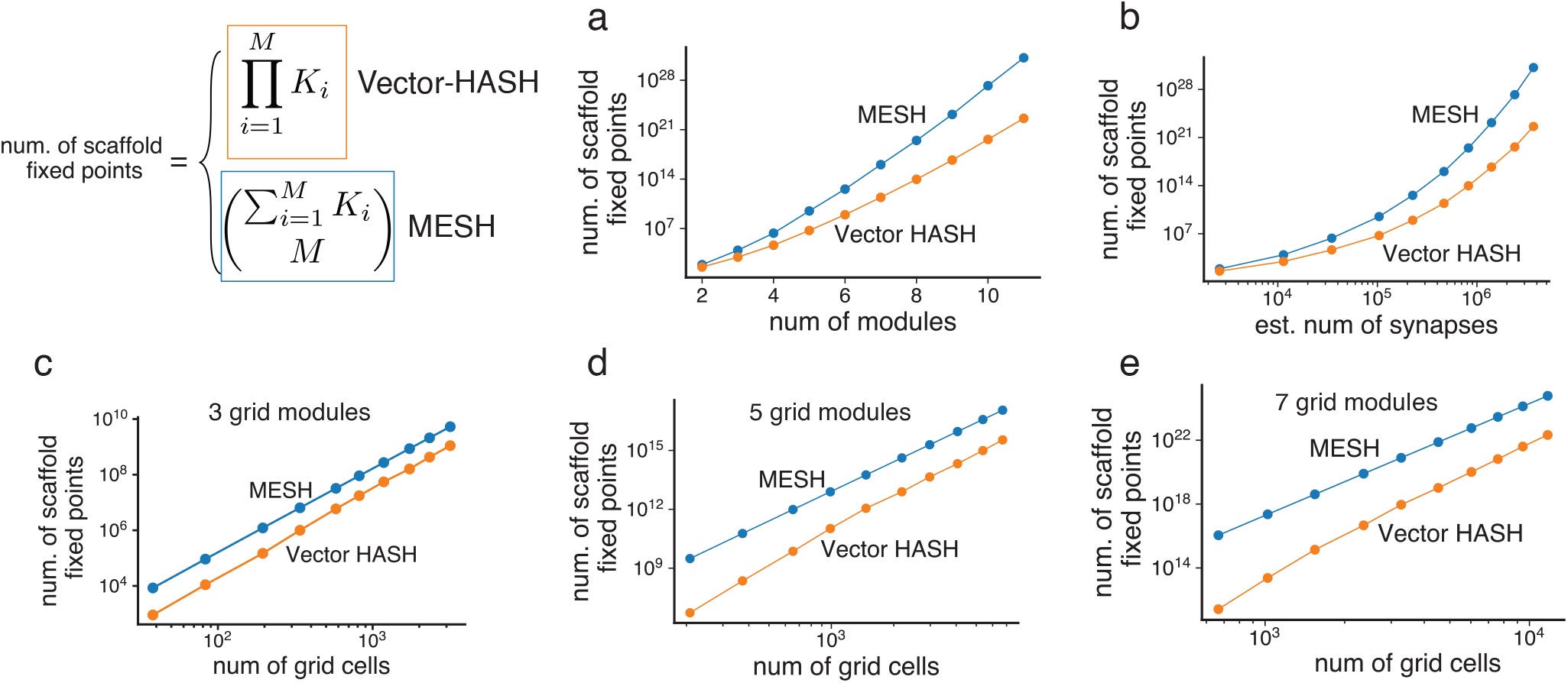
Theoretical capacity results in Vector-HaSH relative to MESH. The number of scaffold fixed points increases exponentially in the number of modules (a), and faster than a power law, but slower than exponentially in the number of synapses (b). The number of synapses with increasing number of modules were estimated based on a number of hippocampal cells extrapolated from Fig. 2f. The number of fixed points increases as a power law with the number of grid cells at a fixed number of modules, with the power law exponent increasing with the number of modules.

**Figure S2.**
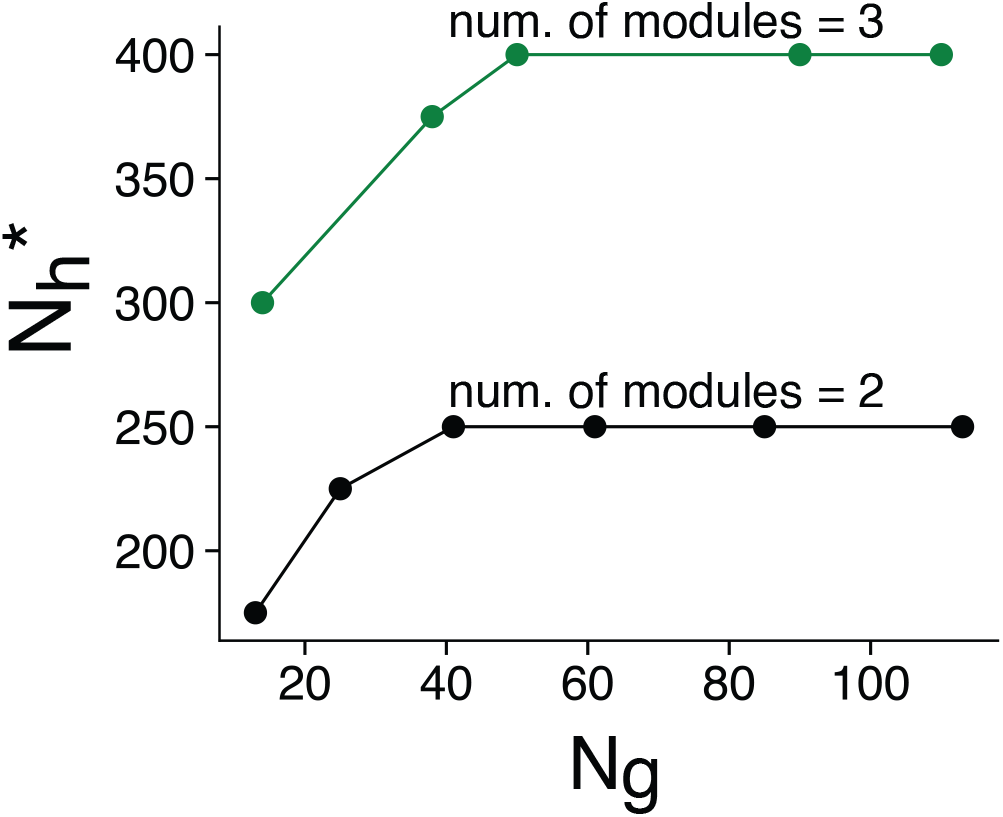
Critical number of hippocampal cells necessary to support all scaffold fixed points is asymptotically independent of the number of grid cells. For a given number of modules, the critical number of hippocampal cells, 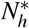 increases slowly with the number of grid cells, but then asymptotically approaches a constant, as expected from the theoretical results in Sec. C.1.

**Figure S3.**
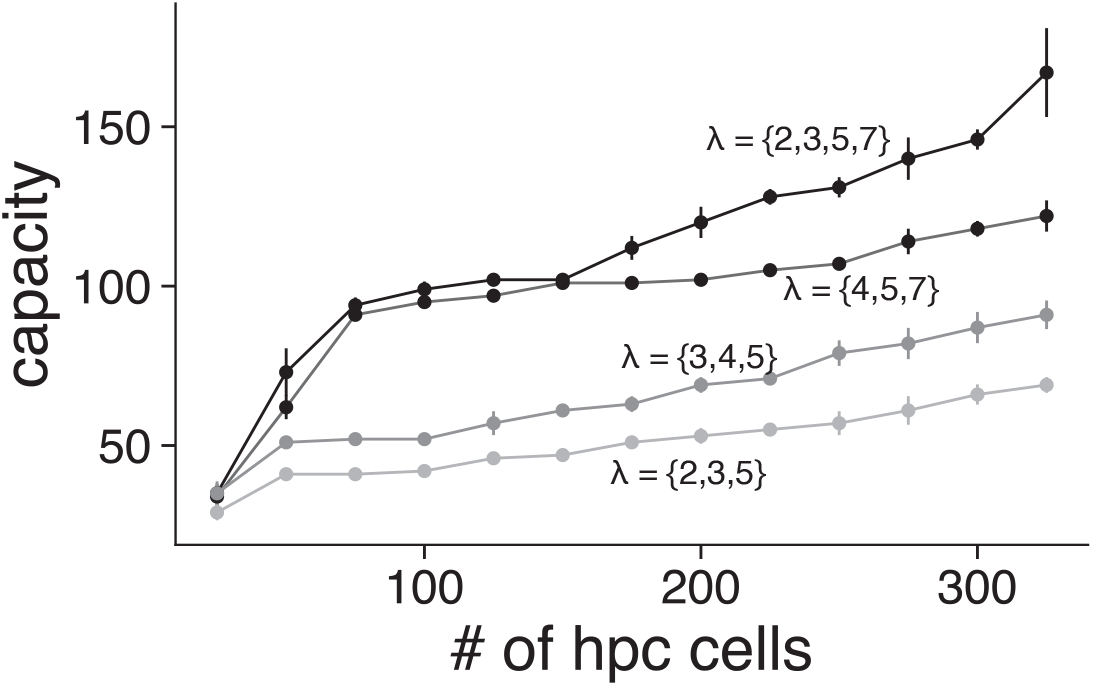
Scaffold constructed with bi-directional learning between grid states and sparse hippocampal states has low capacity. As seen in Fig. 2f *inset*, construction of random sparse hippocampal states with bidirectional learning between grid and hippocampal states results in a scaffold that exhibits catastrophic forgetting. We calculate the capacity of the network as the largest number of trained patterns such that all trained patterns are stored as fixed points. Note that this capacity is limited by the number of hippocampal cells, as might be expected from Hopfield like capacity bounds.

**Figure S4.**
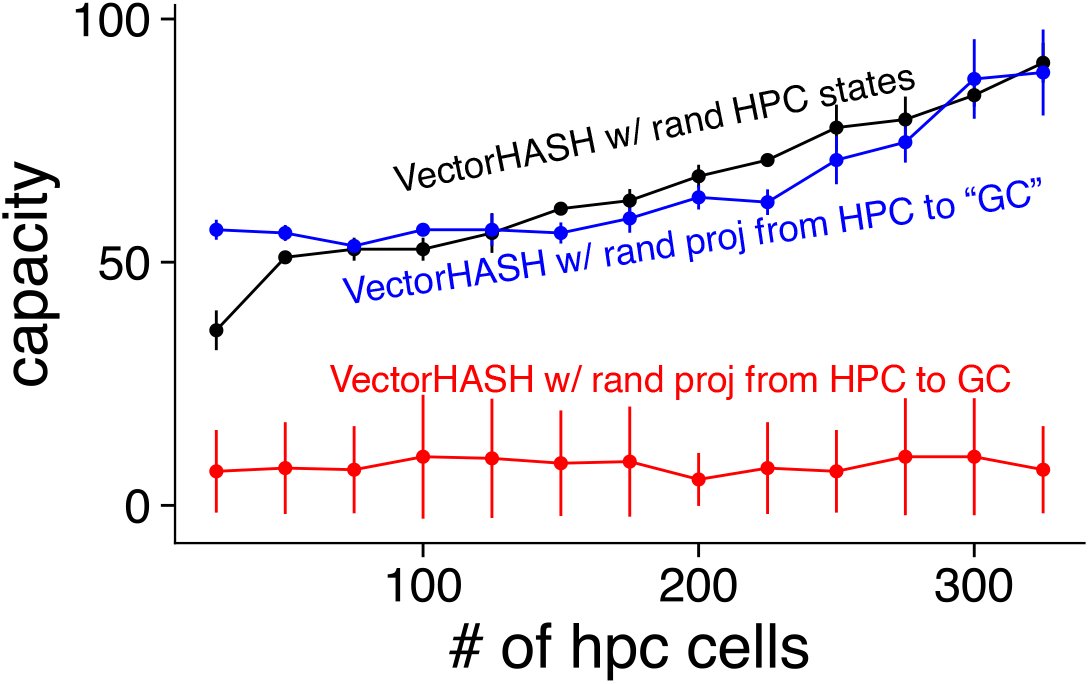
Scaffold constructed with random projections from hippocampal cells to grid cells has low capacity. Instead of the random grid-to-hippocampal projections in Vector-HaSH, we consider instead random projections from random sparse hippocampal states to the grid cell layer, with pseudoinverse learning for the return projections. In the blue curve, the hippocampal inputs to grid cells result in states that are not grid-like. In red, the random sparse drive from hippocampus to grid cells is then mapped to/selects the nearest grid state. Black curve corresponds to bi-directional learning between grid states and sparse hippocampal states, Fig. S3. For both red and blue curves, sparse hippocampal states constructed to have the same sparsity as Vector-HaSH with grid periods *λ* = {3, 4, 5}.

**Figure S5.**
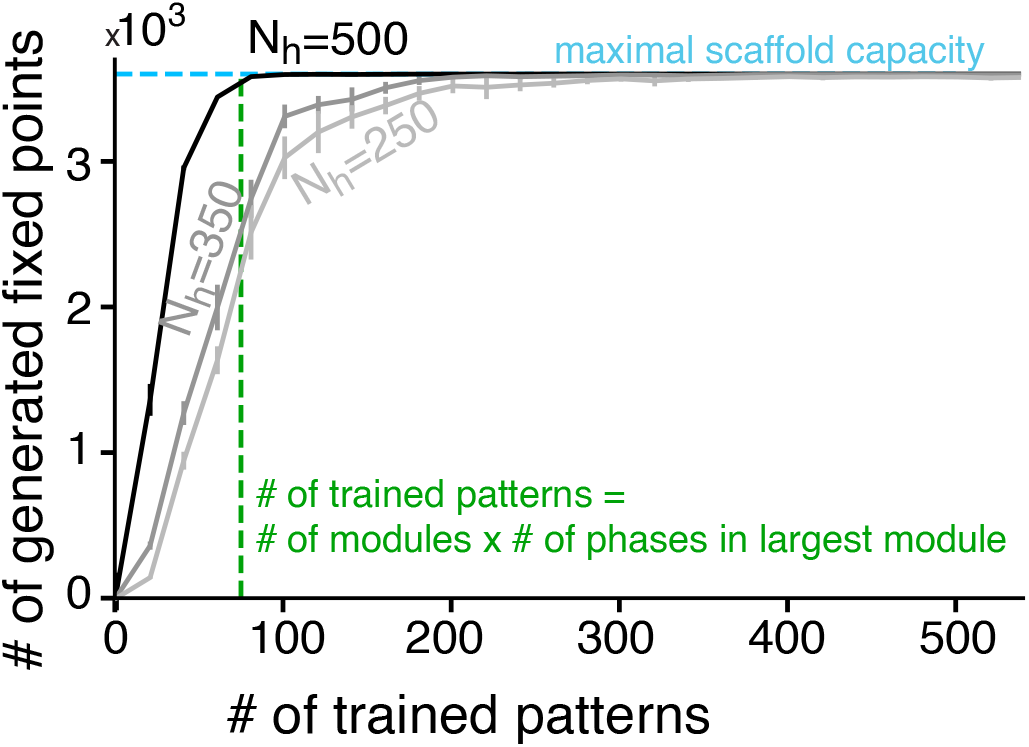
Learning generalization approaches theoretical expectations with increasing. *N*_*h*_ The number of generated fixed points approaches the maximal scaffold capacity for a very small number of learned patterns (see also Fig. 2f). As the number of hippocampal cells increases, the number of learning patterns necessary for complete generalization approaches the theoretical expectation of *M* × *K*_*max*_, as proved in SI Sec. C.4.

**Figure S6.**
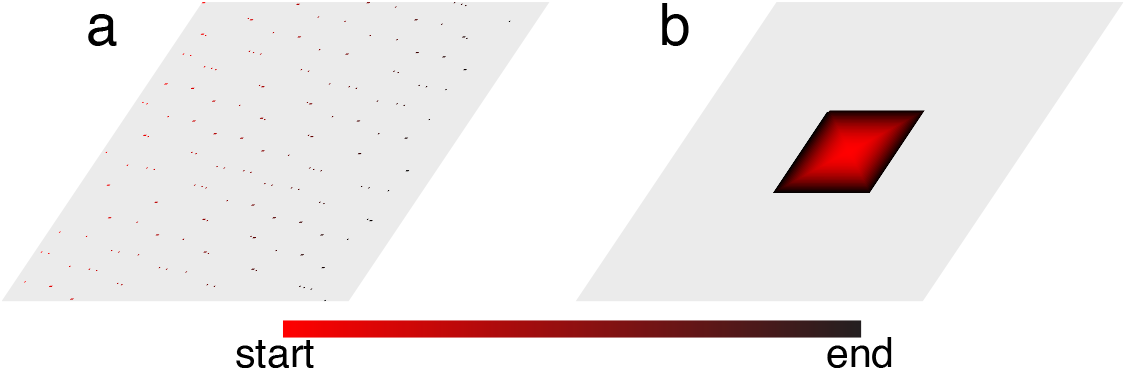
Minimum learning region for generalization of fixed point dynamics at all grid patterns. As seen in Figs. 2f, S5 all the exponentially many scaffold states are stabilized after learning from only a small number of grid patterns. Here we show visually the minimum learning region that results in complete generalization to all scaffold fixed points. (a) shows the minimal learning region for the fastest possible generalization to all scaffold states (see SI Sec. C.4 for an analytic proof), (b) shows the smallest region needed for a path that spans a two-dimensional contiguous region, generated by a spiraling outward path. Both (a) and (b) are shown corresponding to a scaffold size of 44100, generated with *λ* = {2, 3, 5, 7}. As argued in SI Sec. C.4, the minimum learning area as a fraction for complete generalization approaches zero with increasing scaffold sizes.

**Figure S7.**
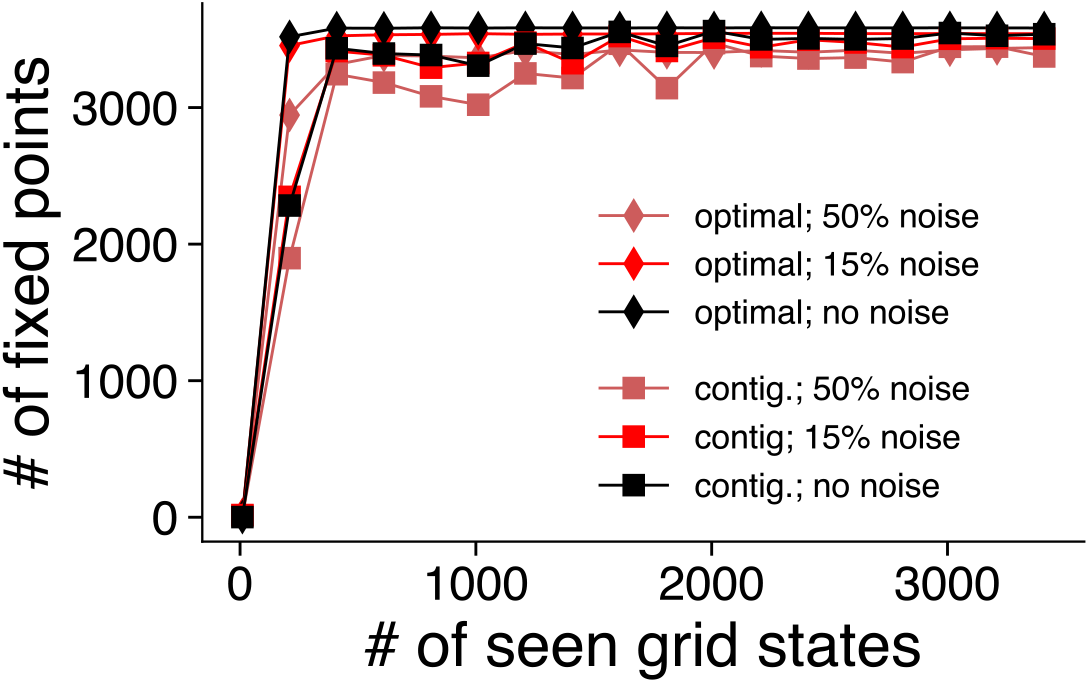
Learning generalization of scaffold is unaffected by noise during training. The number of generated fixed points approaches the maximal scaffold capacity for approximately similar number of learned patterns despite varied levels of hippocampal noise during training (Percentage strength of noise calculated as ratio of magnitude of noise vector to average magnitude of hippocampal state vector). This result holds for the optimal order for scaffold generalization, denoted by the diamond markers (cf. SI Sec. C.4, SI Fig. S6a) as well as a two-dimensional contiguous region (cf SI Fig. S6b).

**Figure S8.**
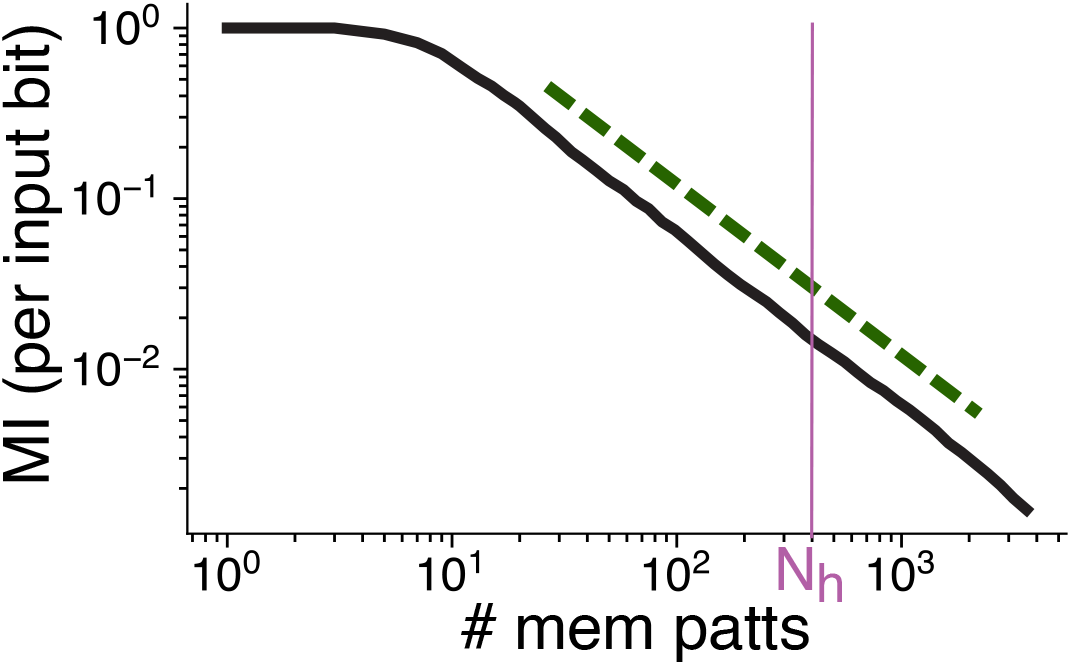
Hebbian learning between sensory layer and scaffold also produces memory continuum. A memory continuum is obtained in Vector-HaSH even if the weights between the sensory and hippocampal layers are bi-directionally trained using Hebbian learning (instead of pseudoinverse learning, as in Fig. 3. This continuum is also asymptotically proportional to the theoretical bound on memory capacity (forest green dashed line indicative of slope of theoretical upper bound, vertical and horizontal position of dashed line is arbitrary). However, the proportionality constant is lower, with the gradual degradation of information recall occurring well before *N*_*h*_. Vector-HaSH parameters identical to Fig. 3c with *λ* = {3, 4, 5}.

**Figure S9.**
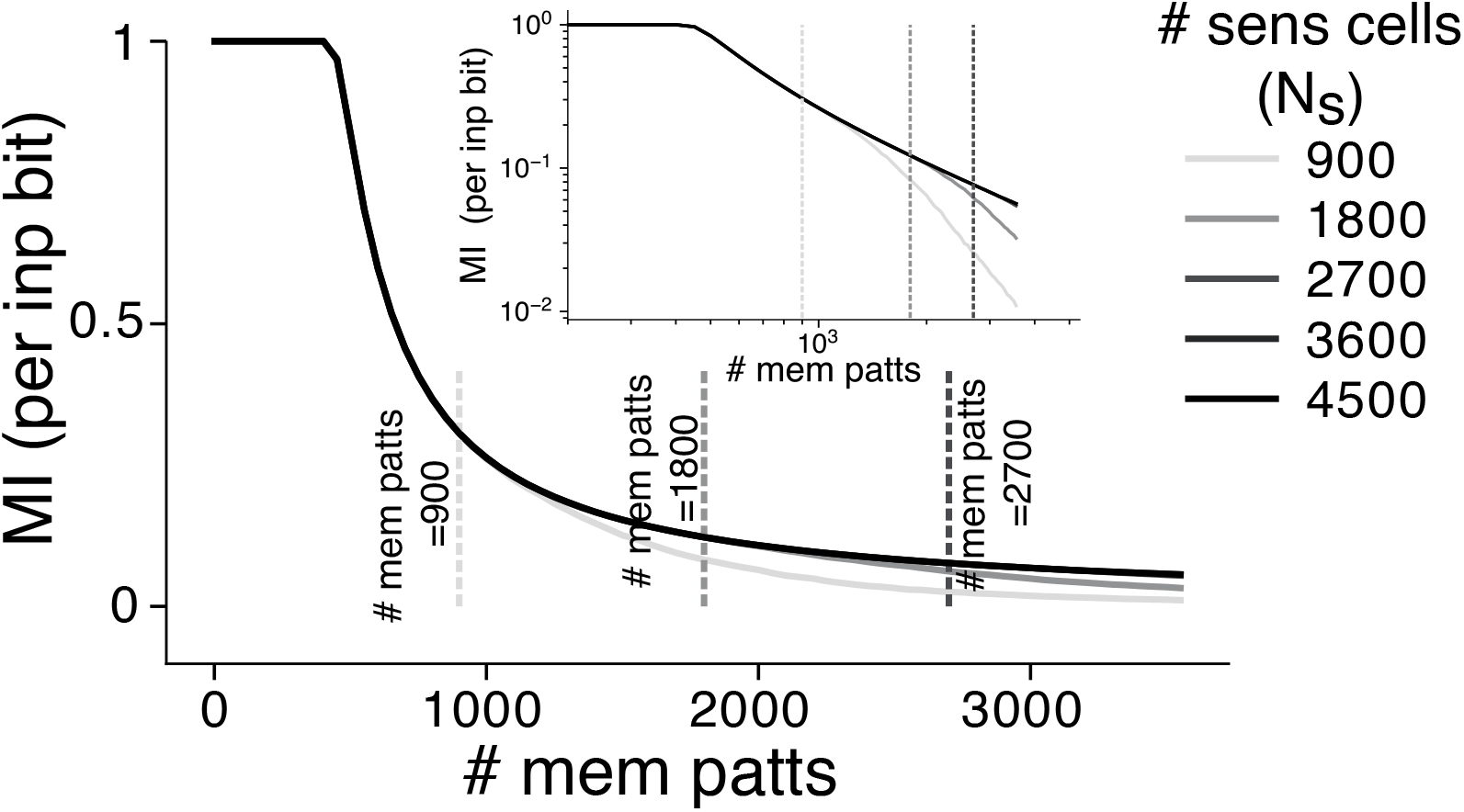
Effect of varying *N*_*s*_ on memory continuum. As shown in SI Sec. D.1, the number of sensory cells determines the number of scaffold states that can be exactly recovered through the sensory-to-hippocampal weights. For *N*_*s*_ less than the total number of scaffold states, the obtained memory continuum is distorted towards the tail for larger than *N*_*s*_ patterns stored. For all *N*_*s*_ larger than or equal to the number of scaffold states, the memory continuum is identical, corresponding to the results shown in Fig. 3.

**Figure S10.**
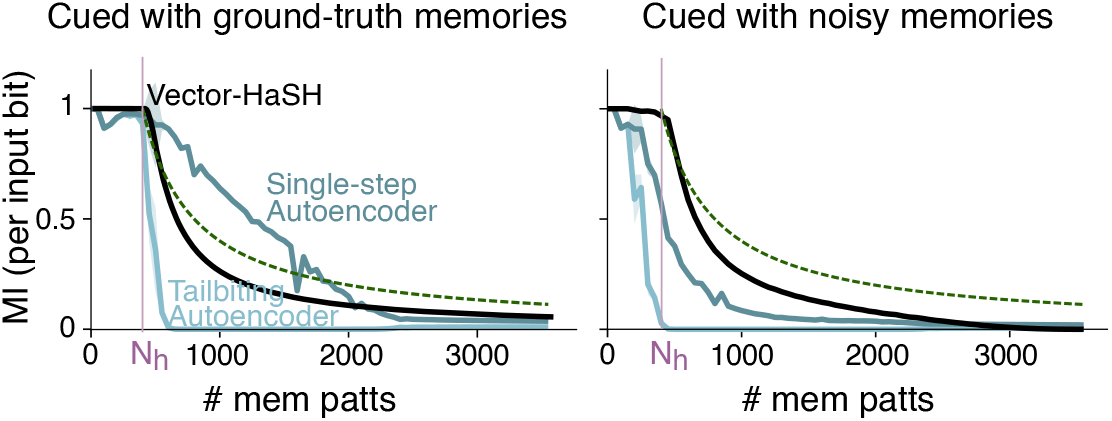
Vector-HaSH outperforms Autoencoders, particularly when recovering from noisy cues. *Left* When cued with ground-truth memorized sensory patterns, Vector-HaSH recovers a gradually degraded amount of information per pattern (cf. Fig. 3), unlike the memory cliff shown by tailbiting Autoencoders trained as associative memories^99^. Naively however, it appears that this memory cliff is absent in a single step (i.e., non-tailbiting) of the Autoencoder. However, we see in *Right* that single-step Autoencoders are not associative memories, since they are unable to reconstruct memories from corrupted cues. Here grid periods were set to *λ* ={3, 4, 5}, with *N*_*h*_ = 400. Stored sensory cues were random binary {−1, 1} patterns, and noisy cues were generated by flipping 10% of bits from a given memorized sensory pattern.

**Figure S11.**
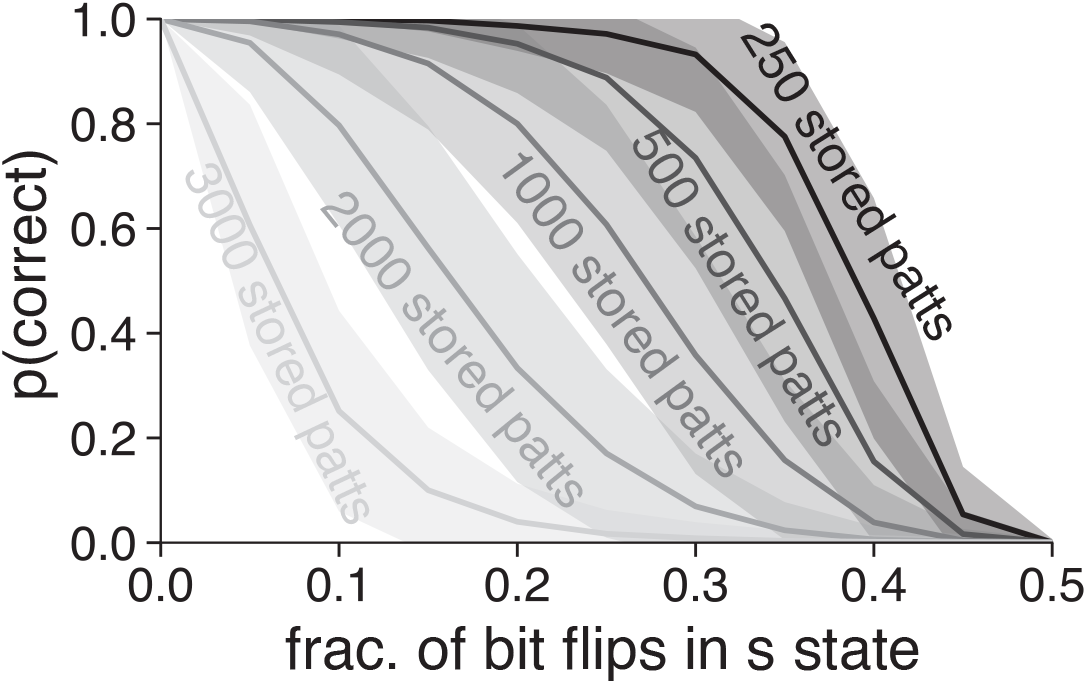
Basin structure for recovery of sensory hippocampal and grid states varies with number of stored patterns. While the scaffold has a large number of well-structured basins (cf. Fig. 2, SI Sec. C.1), the basins for sensory recovery are additionally governed by the heteroassociative learning between the sensory states and the scaffold. As a result, the basin sizes reduce with increasingly large number of stored patterns, due to overcrowding of the number of stored states within the sensory-to-hippocampal weights. The grid periods were set to *λ* = {3, 4, 5}, with *N*_*h*_ = 400, resulting in a maximal scaffold capacity of 3600 patterns, with perfect sensory recovery up to 400 patterns. For more than 400 stored patterns, *p*(*correct*) refers to the probability of exact recovery of grid and hippocampal states, and probability of reliable recovery of the sensory state (which is not exact due to being in the memory continuum regime, Fig. 3.

**Figure S12.**
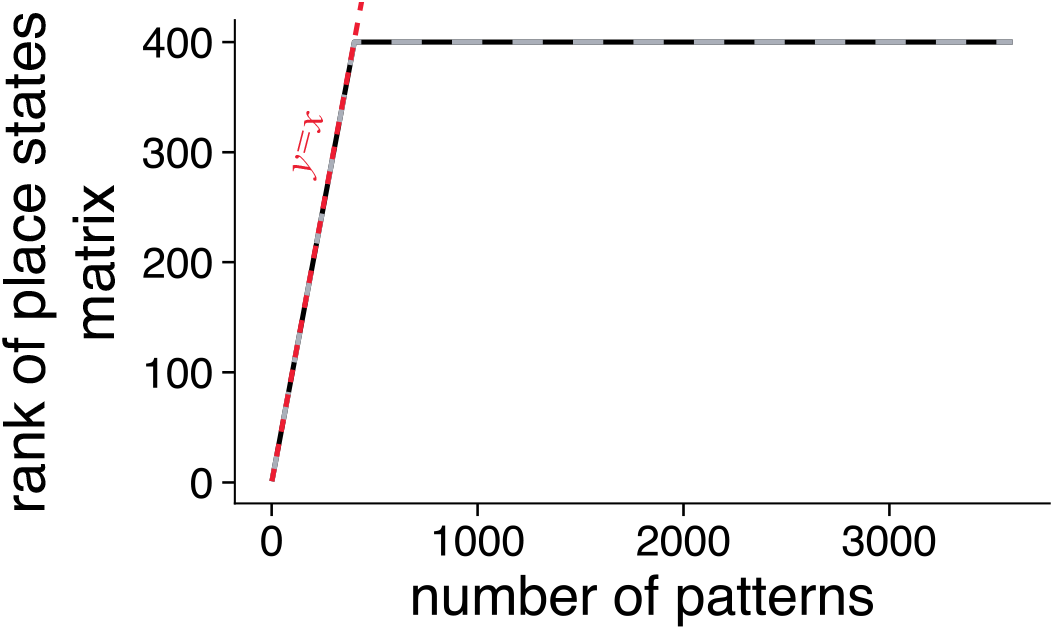
Hippocampal states form a strongly full rank matrix. Rank of the *N*_*h*_ × *N*_*patts*_ hippocampal states matrix for varying number of patterns *N*_*patts*_ for two different random permutations of the ordering of hippocampal states shown in black and gray. For up to *N*_*patts*_ ≤ *N*_*h*_ the rank of the matrix is *N*_*patts*_ (as indicated by the red *y* = *x* line), and is there after *N*_*h*_ for larger numbers of patterns.

**Figure S13.**
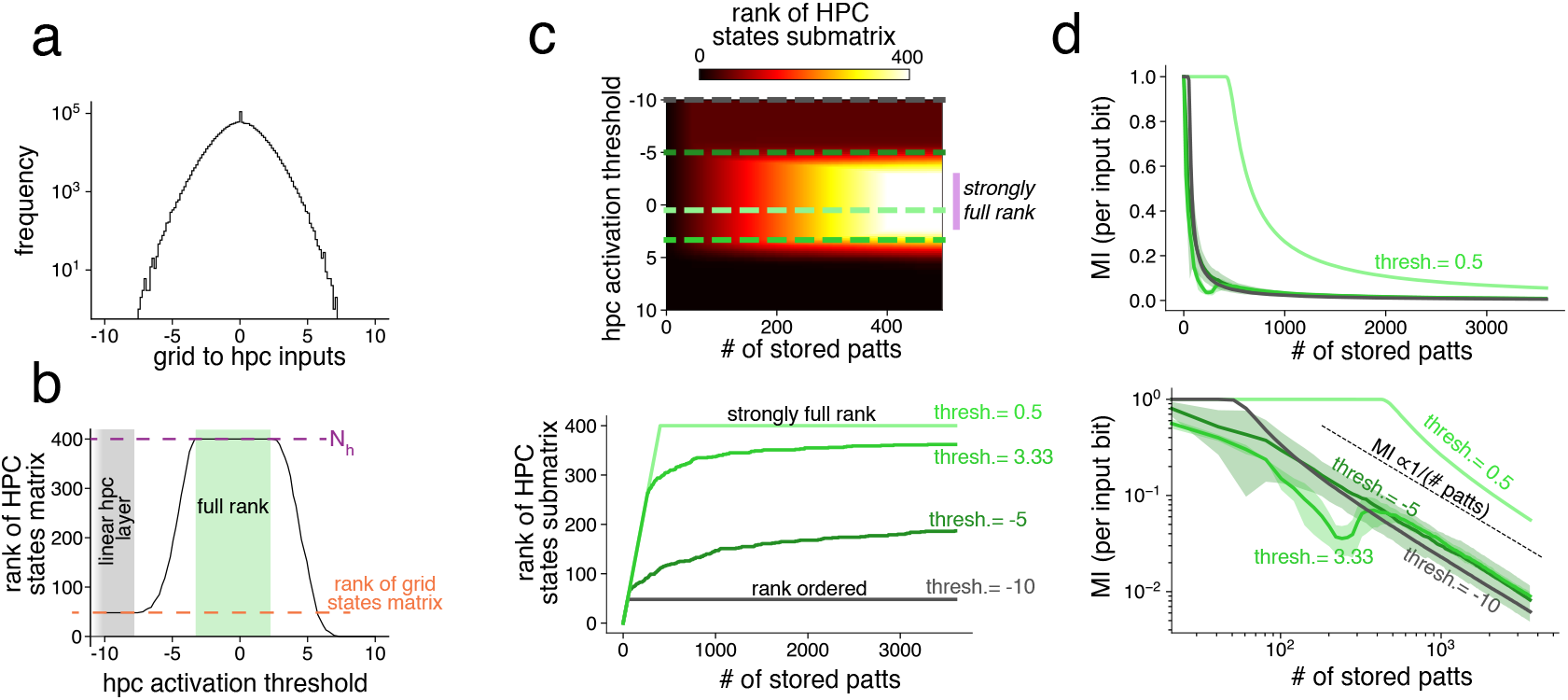
Activation threshold applied in the hippocampal layer dictates nature of memory continuum. (a) distribution of pre-nonlinearity inputs to the hippocampal layer from grid cells. Any activation threshold above the largest value (∼ 7.5) results in zero hippocampal activity, and any threshold below the smallest value (∼−7.5) results in a purely linear hippocampal layer. (b) A linear hippocampal layer (corresponding to thresholds in the gray region) results in a HPC states matrix of rank equal to the rank of the grid cell states matrix (which equals *N*_*g*_ − *M* + 1 as shown in Ref.^115^), whereas a range of thresholds (shown in green) result in a full rank HPC states matrix. (c)*Top*: The rank of the *N*_*h*_ × *N*_*patts*_ submatrix of hippocampal states constructed over the first *N*_*patts*_. Here the hippocampal states have been ordered according to the optimal order that leads to fastest scaffold learning generalization (Sec. C.4). *Bottom*: Rank versus *N*_*patts*_ for the particular values of thresholds considered in panel (d). At a threshold of 0.5 (the value used in almost all simulations in the main text, see *Methods* for more details) we see that the hpc states matrix is strongly full rank. Moreover, as seen in Fig. S12, this matrix is strongly full rank independent of the ordering of the scaffold states. At the lowest threshold value, corresponding to a linear hippocampal layer, the matrix appears to be rank ordered (i.e., submatrices formed by considering the first *k* column vectors of *H* are full rank for all *k*). However, for a linear hippocampal layer the rank ordering of the matrix is dependent on the ordering of the scaffold states, as examined in Fig. S14. (d) Information recovered per input bit as a function of the number of patterns stored in the network (similar to Fig. 3d) for varying threshold values on a linear scale (*top*) and a logarithmic scale (*bottom*). The strongly full rank matrix (identical the the *λ* = {3, 4, 5} curve in Fig. 3d) and the rank ordered matrix both demonstrate perfect recovery up to a knee; all values of thresholds result in a smooth decay of recovered information that is asymptotically proportional to a theoretically expected bound that scales inversely with the number of stored patterns

**Figure S14.**
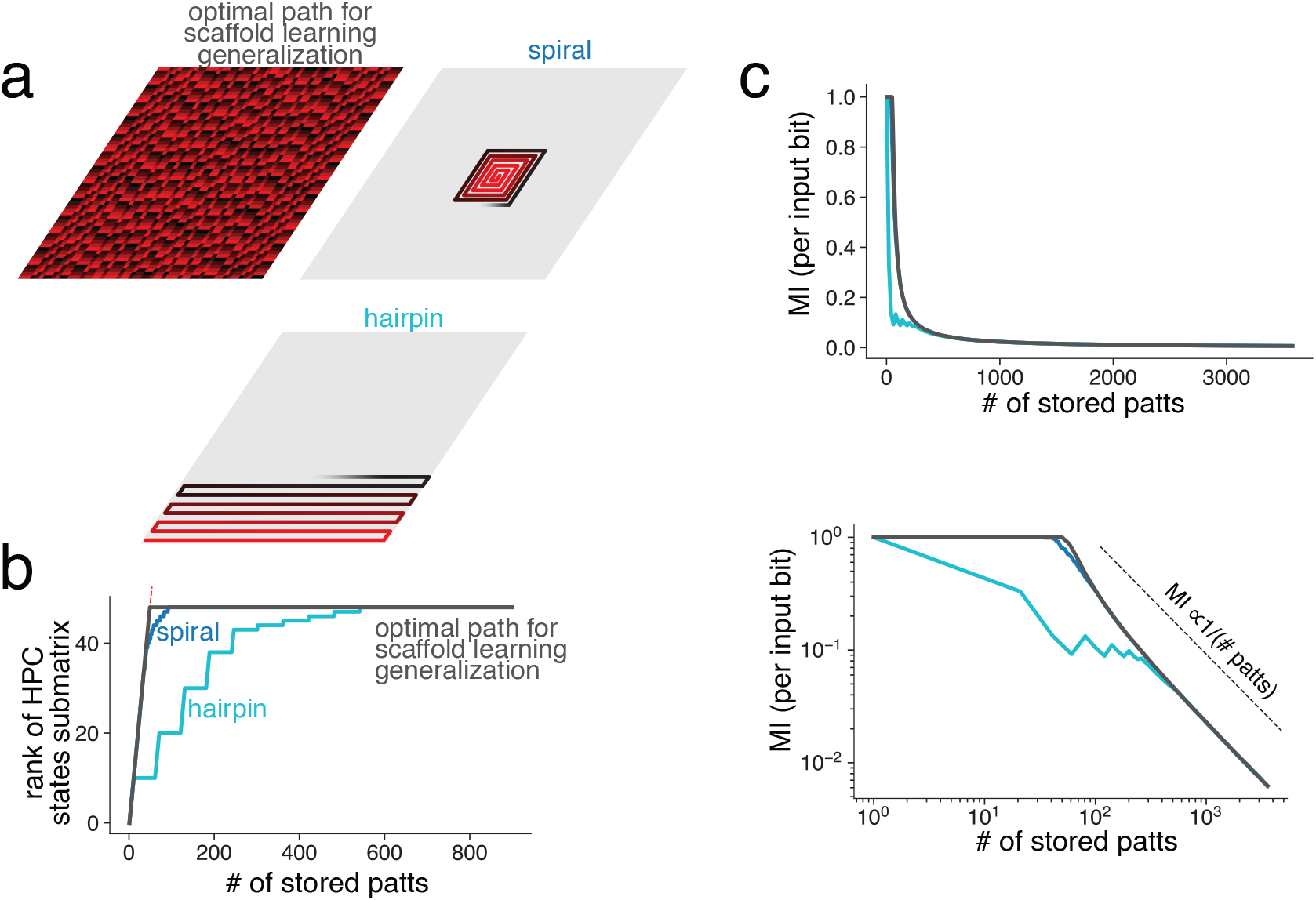
Linear hippocampal layer generates memory continuum only for specific ordering of scaffold states during learning. (a) Three examples of potential ordering of scaffold states that could be considered: *top left* the discontinuous path that leads to the fastest scaffold learning generalization (Sec. C.4) shown for *λ* = {3, 4, 5} ; *top right* a continuous spiral path; *bottom* a continuous ‘hairpin’ path. (b) The hpc states submatrix is rank ordered along the optimal path, and approximately rank ordered for the continuous path. The hairpin path however is significantly deviated from a rank ordered matrix. (c) Information recovered per input bit as a function of the number of patterns stored in the network (similar to Fig. 3d). The rank ordered matrix demonstrates perfect recovery up to a knee at the rank of the grid states matrix; this is also closely approximated by the spiral ordered matrix. A hairpin ordering however results in poor information recovery even at a small number of patterns. In all cases, the asymptotic decay of information is inversely proportional to the number of patterns, as would be expected from theoretical information bounds.

**Figure S15.**
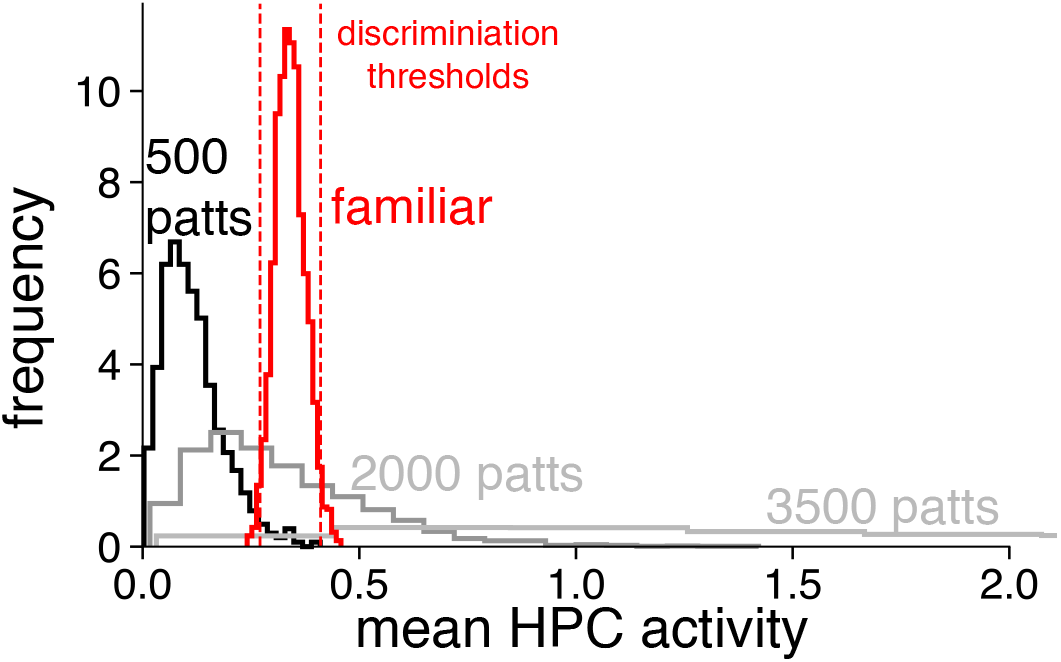
Mean activity in hippocampal layer can be used for novelty detection. The mean activity in the hippocampal layer for familiar patterns presents a narrow distribution. The mean hippocampal activity for novel patterns is strongly dependent on the number of stored patterns. The narrowness of the familiar pattern distribution allows for discrimination thresholds to be placed on either side (at two standard deviations away from the mean) to result in classification accuracy as shown in Fig. 3i

**Figure S16.**
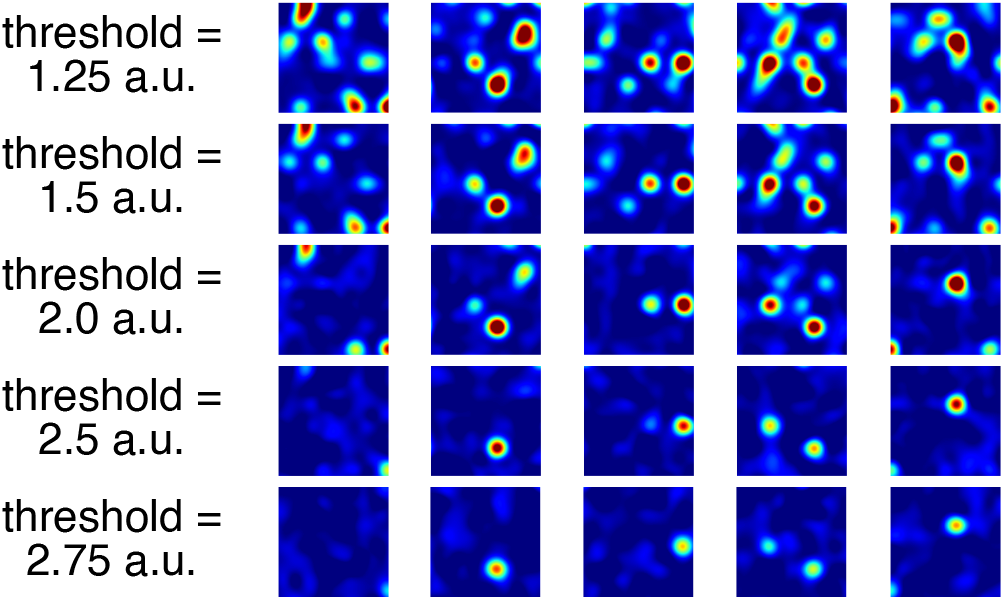
Additional examples of place fields, with varying thresholds. Random place cells generated with all parameters similar to those in Fig. 4 except for the threshold, *θ* . Varying the threshold varies the sparsity of the place fields. At low sparsity with several place fields within the same environment, fields do not demonstrate any grid like periodic tuning curve, despite receiving grid inputs, due to the mixture of grid inputs across cells with different periods.

**Figure S17.**
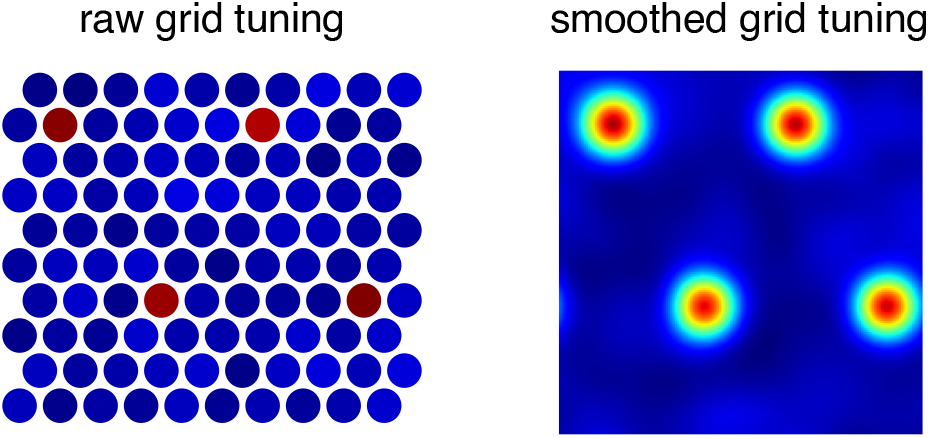
Grid tuning on discrete hexagonal lattice. *Left*: Tuning curve of a randomly selected grid cells in discrete space on a hexagonal lattice. *Right*: The same tuning curve after the smoothing procedure used for plotting, as described in *Methods*

**Figure S18.**
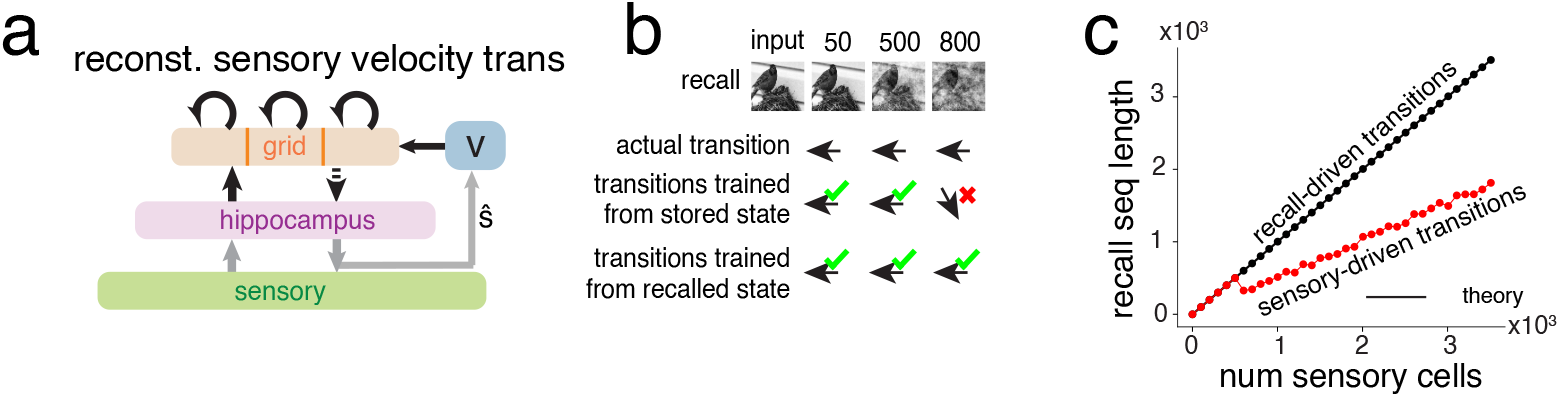
Sensory driven transitions must be reconstructed from recalled states. (a) Architecture for sensory based reconstruction of next-step transitions for sequence learning, Fig. 6a, *bottom*. (b) As the number of stored patterns increases, the recalled sensory state gradually degrades; as a result, reconstruction from mapping trained on ground truth sensory states can lead to inaccuracies. (c) More quantitatively, transitions trained on recalled sensory states result in sequence reconstruction of length up to the number of sensory cells (theory in SI Sec. D.7), whereas transitions trained on ground truth sensory states has a lower sequence capacity.

**Figure S19.**
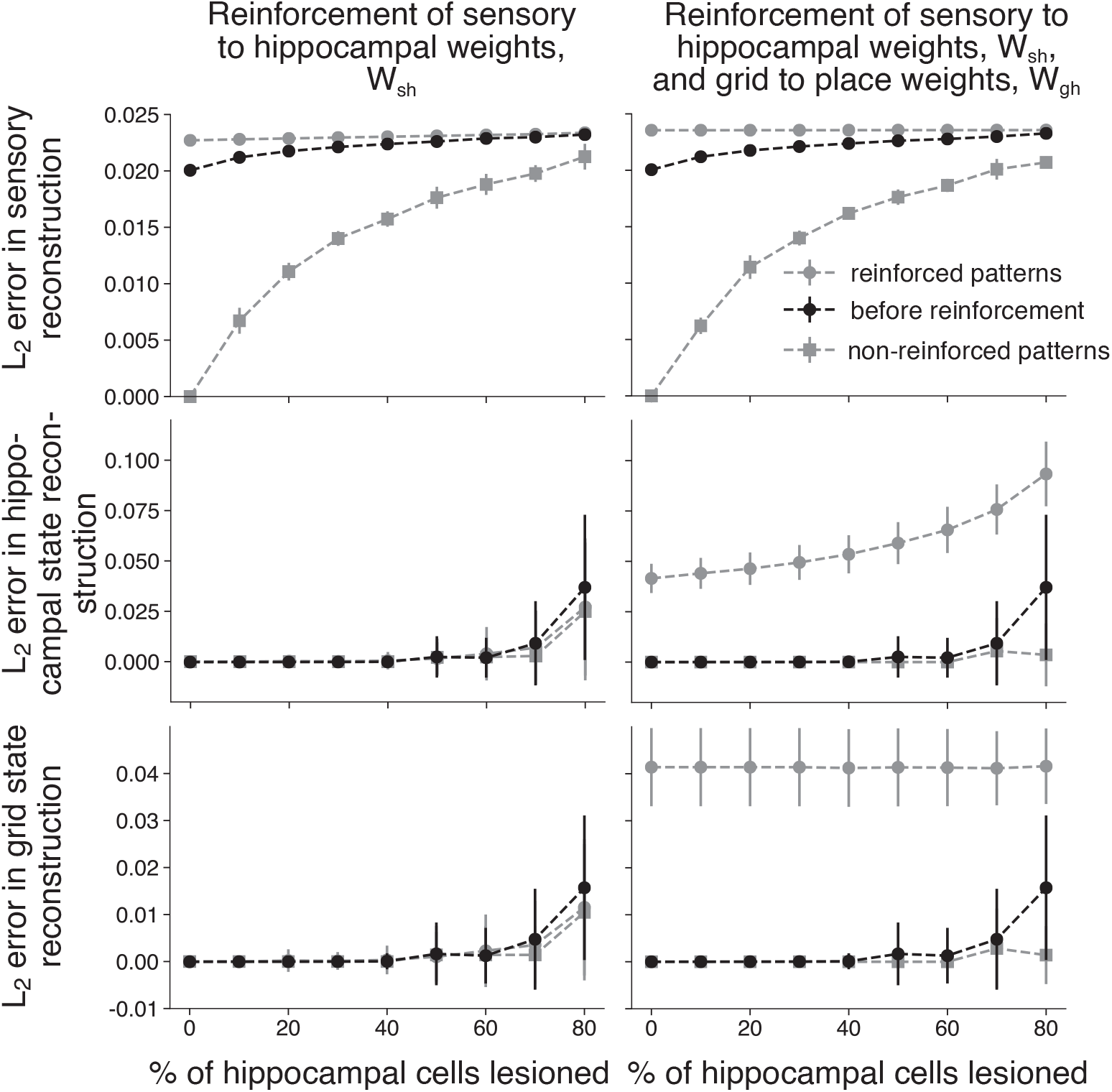
Reconstruction error in each layer of Vector-HaSH when tested for MTT by reinforcing the model weights for a subset of repeated patterns. Left: Results when only *W*_*sh*_ weights are reinforced, assuming pre-trained scaffold weights *W*_*gh*_. Right: Results when all of the learnable weights in Vector-HaSH *W*_*hs*_, *W*_*sh*_ and *W*_*gh*_ are reinforced. Note that *W*_*hs*_ reinforcement mathematically doesn’t change *W*_*hs*_ as describe in Sec. 1.

**Figure S20.**
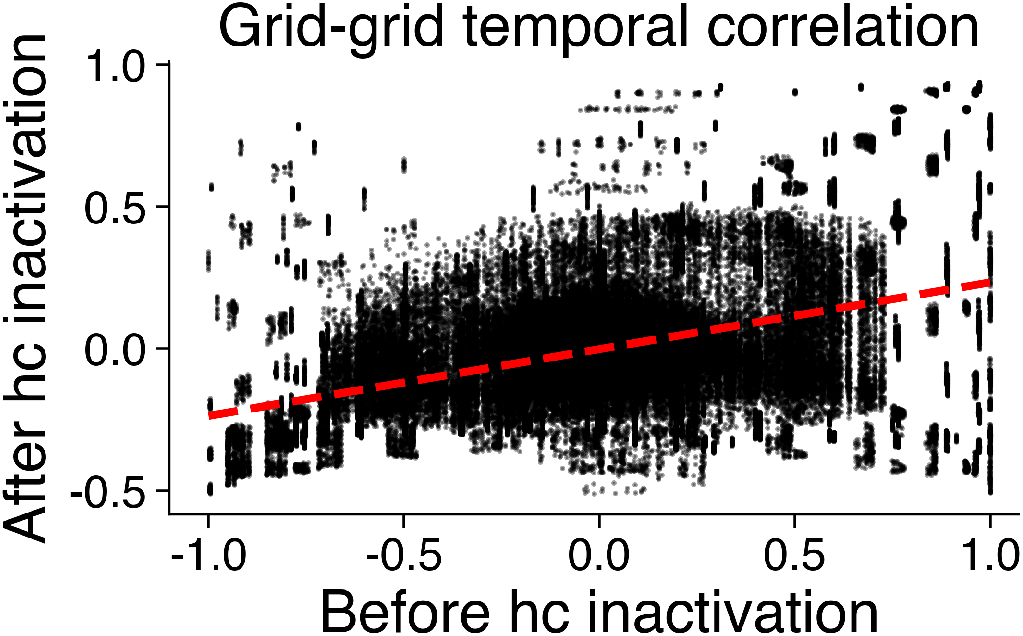
Correlation between grid states before and after hippocampal inactivation. Best-fit line (shown as red-dashed curve) used to compute correlation between cell-cell correlations, that is then compared with a null hypothesis based on shuffled correlations in Fig. 7f

**Figure S21.**
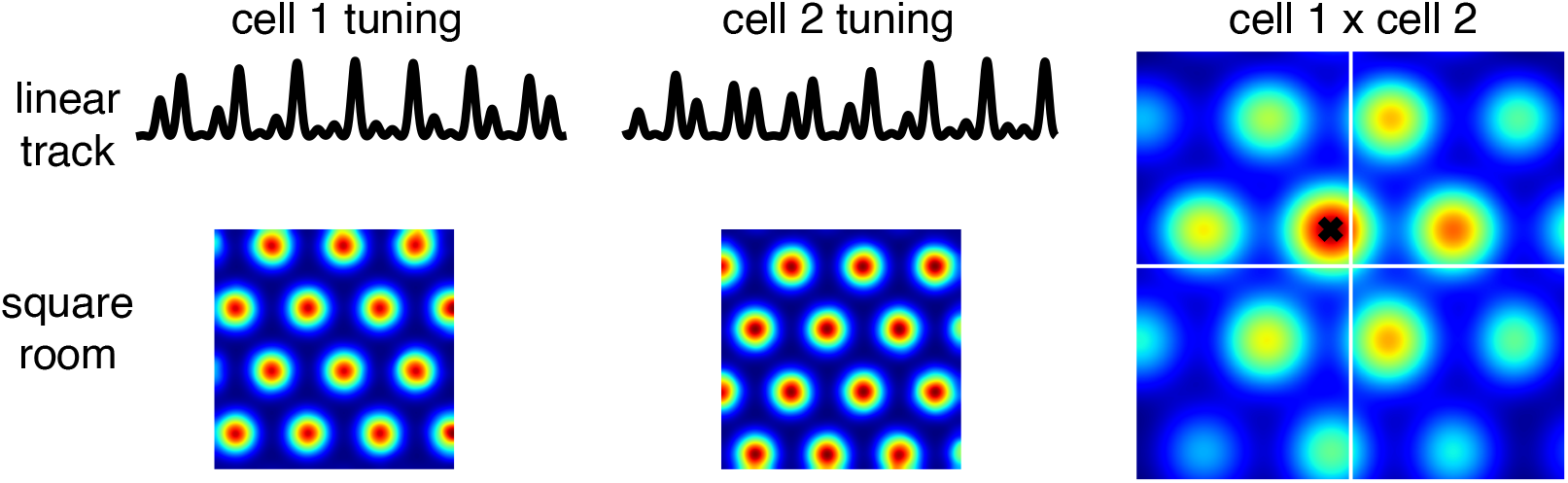
rigid correlation between grid states across environments of different geometries. *Left*: Spatial tuning curves of two grid cells on a one-dimensional linear track and in a two-dimensional square room. *Right*: relative phase estimated from the one-dimensional tuning curves (via the analysis in Ref. 34) shown as the black cross aligns with the peak of the spatial cross correlation, capturing the relative phase estimated from the two-dimensional tuning curves. See *Methods* for more details.

**Figure S22.**
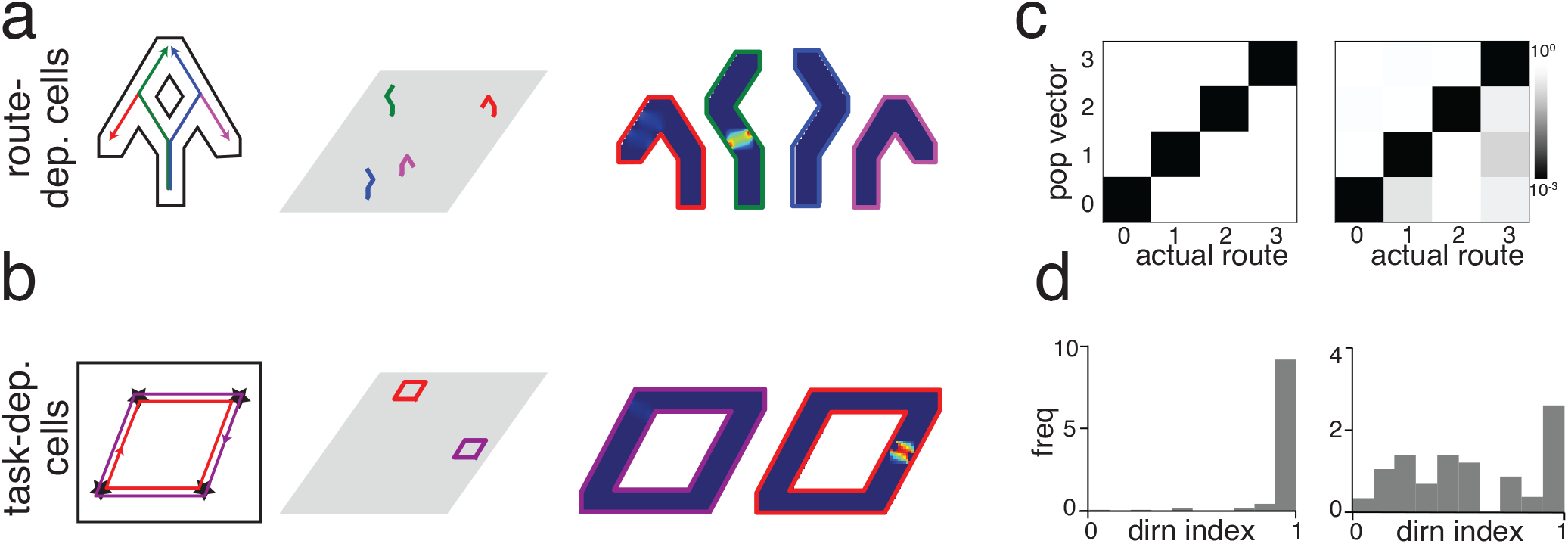
Vector-HaSH reproduces route-dependent and task-dependent cells. (a-b) Route-dependent spatial tuning^120^ and task-dependent cells in an open arena^122^ can be modeled by Vector-HaSH if different trajectories traverse different regions of the grid coding space (grid cell remapping), similar to Fig. 7 (c-d) Quantification of contextual/route selectivity of responses from the hippocampus in Vector-HaSH (left column) and experiments (right column), corresponding to (a-b). (c) Ensemble decoding of individual trajectories based on route population vectors, with color indicating the p-value of correct matches made by chance. (SI Sec.4.3 and SI Fig. S23)^120^). (d) The directionality index (a normalized metric for the difference in neural activity for different run directions^120,121^) shows that a majority of hippocampal cells have directional fields^121^ (Qualitatively similar results hold for a radial maze environment^122^, SI Fig. S24.)

**Figure S23.**
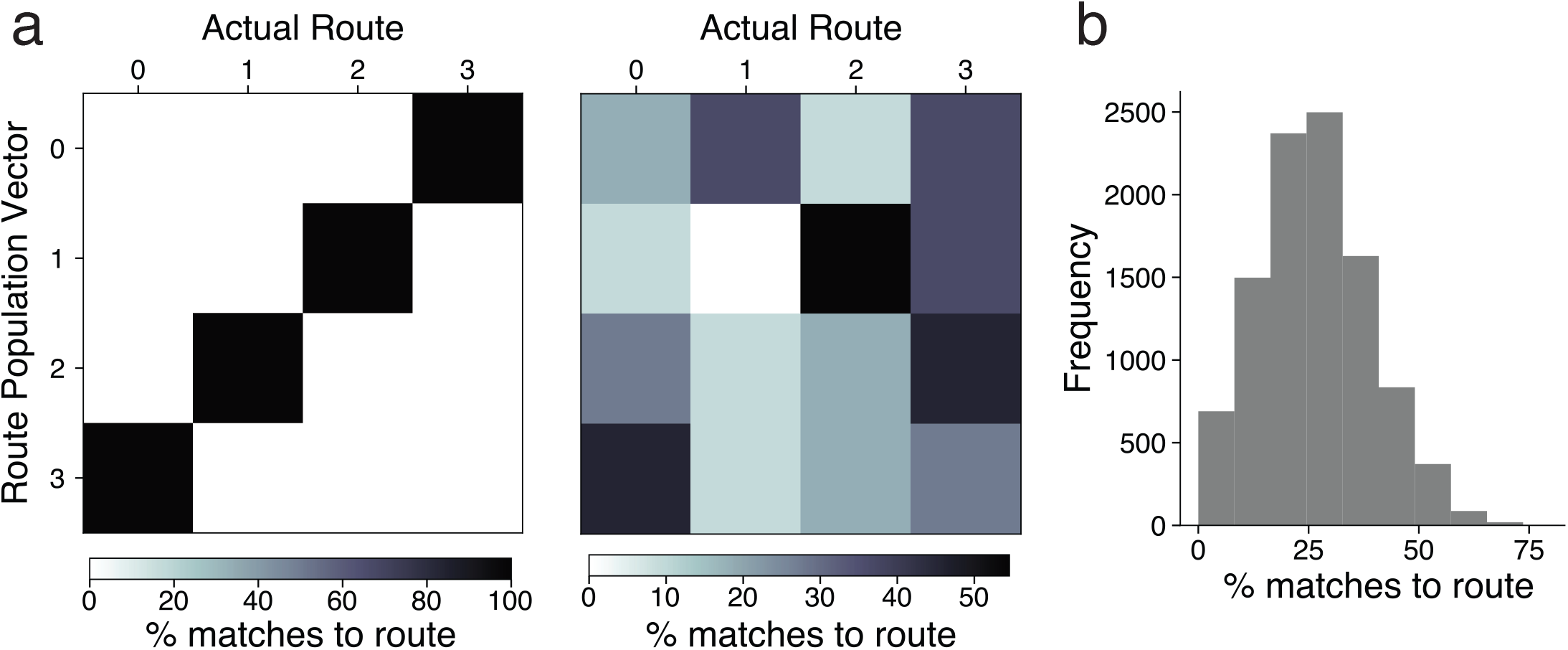
Analysis method for route encoding results in Fig. S22c. (a) Left: Trajectory population vectors (PVs) were compared to route-PVs and matched according to the highest cosine similarity score. Elements of the matrix show the percentage number of each trajectory matched to each of the four route-PVs. Right: Matches were also made using shuffled data, where each trajectory was randomly assigned to one of the four routes, thus shuffling the route identity of the trajectories. The matrix elements here show the same as *a* except that this data is for one representative shuffle (10000 were conducted in total). (b) Distribution of percentage correct matches for trajectory PV of Route 3 to its route-PV for all 10000 shuffles.

**Figure S24.**
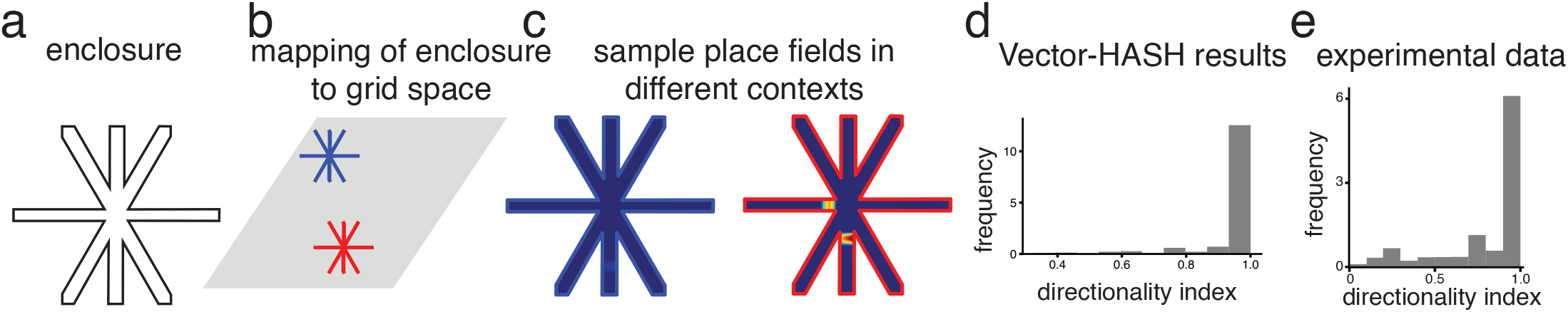
Vector-HaSH reproduces directional place fields on an 8-arm radial maze. (a) An 8-arm radial maze apparatus similar to the experiment^122^. (b) Inbound (towards the center) and outbound (away from the center) trajectories on the 8-arm radial maze represented separately in the grid coding space. (c) Fields of a representative hippocampal cell on inbound trajectories (left) and outbound trajectories (right) (d) Directionality index of place cells from Vector-HaSH showing that majority of the cells have directional fields. (e) Directionality index of place cells from the experimental data^122^.

**Figure S25.**
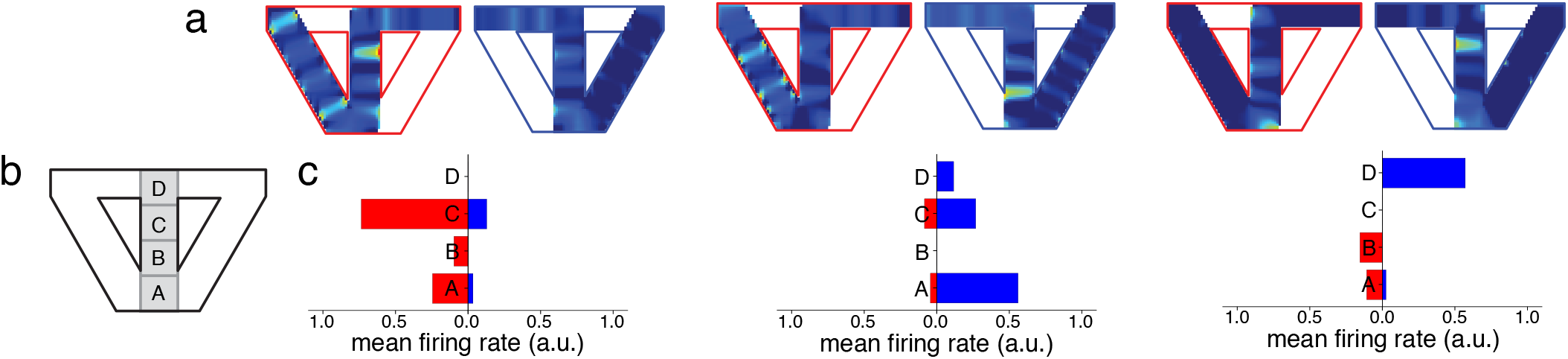
Splitter cells. (a) Fields of three representative hippocampal cells on the Right-Turn and Left-Turn trials. (b) The central stem of the continuous alternation task apparatus is divided into 4 equal regions for data analysis following the analysis conducted on the experimental data^119^. (c) Mean activation of the three hippocampal cells shown in *(a)* computed for each of the four regions defined in *(b)*. The cells show different activity patterns as Vector-HaSH traverses the central stem on Left-Turn and Right-Turn trials.

**Figure S26.**
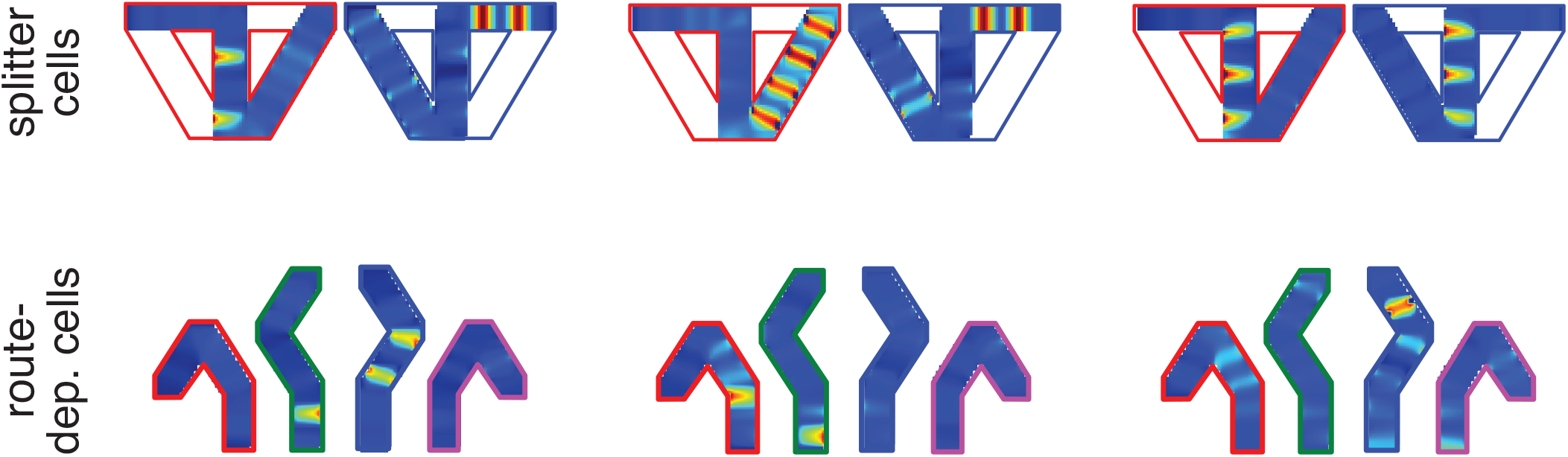
Vector-HaSH splitter cell-like phenomena in grid cells. Grid cells demonstrate different represntations in different contexts, due to contexts being represented through different regions of the grid coding space, Fig. 7g.

**Figure S27.**
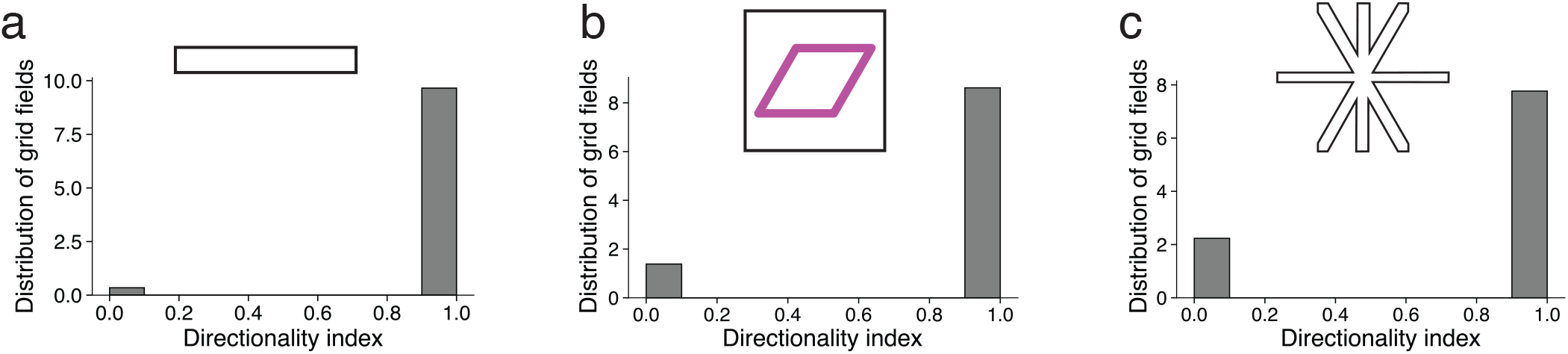
Vector-HaSH predicts directional grid fields. Directionality index of grid cells showing that in Vector-HaSH majority of the grid cells have directional fields in one dimensional environments (environment in (a)^121^), and on directed routes in two-dimensional environments (environments in b,c^122^).

## Footnotes

1 e.g., if *M* ∼ 10 (small number of grid modules), *K* = 10^2^ (resolution of 10 grid phases per dimension), the network would yield ∼ 10^20^ stable fixed points with only ∼ 10^3^ total hippocampal and grid cells combined.

2 We show in SI Sec. D.5 that one could simply use Hebbian learning instead of an iterative or standard pseudoinverse learning, while maintaining the same asymptotic capacity, with a smaller constant pre-factor, Fig. S8

3 Had we not earlier assumed that elements of W are drawn from a normal distribution we would have arrived at this same result with different intermediate distribution instead of *χ*^2^ and 𝒩𝒫

